# ASTRAL-X: Scaling Coalescent-Based Species Tree Inference to 300,000 Taxa

**DOI:** 10.64898/2026.07.31.742122

**Authors:** Anik Saha, Md. Shamsuzzoha Bayzid

## Abstract

Advances in genome sequencing have enabled phylogenomic studies involving tens or even hundreds of thousands of species. However, scalability remains a major computational challenge for statistically consistent species tree inference at this scale. ASTRAL, the most widely used coalescent-based species tree estimator, remains limited by computational and memory bottlenecks that make ultra-large analyses impractical. Here we present ASTRAL-X, a complete algorithmic redesign of the ASTRAL framework that overcomes these computational limitations. By fundamentally redesigning the underlying data representations, algorithms, and computational framework, ASTRAL-X dramatically reduces running time while lowering memory requirements to nearly the size of the input–the asymptotically optimal bound–thereby enabling statistically consistent species tree inference directly from unrooted gene trees at an unprecedented scale. ASTRAL-X preserves ASTRAL’s statistical guarantees and achieves accuracy comparable to state-of-the-art methods across simulated and empirical datasets while reconstructing species trees containing 200,000 and 300,000 taxa in only 5 hours and 12 hours, respectively, using modest computational resources. Notably, ASTRAL-X reconstructed the evolutionary history of 9,524 angiosperm species in only 16 minutes. These results enable statistically consistent coalescent-based species tree inference at the scale demanded by emerging Tree of Life initiatives. ASTRAL-X is publicly available at https://github.com/aaniksahaa/ASTRAL-X.

## 1 Introduction

The growing availability of genome-scale data is transforming evolutionary biology, with international biodiversity initiatives aiming to reconstruct different branches of Tree of Life across hundreds of thousands of species. Yet the ability to generate genomic data has begun to outpace our ability to analyze it, making scalable species tree inference one of the central computational challenges in modern phylogenomics.

Genome-scale species tree estimation is complicated by the fact that individual genes often have different evolutionary histories [1, 2]. Such discordance arises from several biological and methodological processes, including incomplete lineage sorting (ILS), gene duplication and loss, horizontal gene transfer, hybridization, and gene tree estimation error [3]. Among these processes, ILS is particularly pervasive and is naturally modeled by the multispecies coalescent (MSC) [4–7]. Because concatenation-based analyses can become statistically inconsistent under the MSC, coalescent-based summary methods, which infer species trees from collections of gene trees while explicitly accounting for gene tree discordance, have become a central framework for phylogenomic inference [8–11].

Among these methods, ASTRAL has emerged as the dominant framework for species tree estimation under the MSC [12]. By maximizing quartet agreement across unrooted gene trees, ASTRAL combines strong statistical guarantees with excellent empirical performance while operating directly on unrooted gene trees. Over the past decade, successive generations of ASTRAL have steadily expanded the framework through improvements in scalability, accuracy, computational efficiency, and functionality, establishing it as the most widely used framework for an increasingly broad range of phylogenomic settings [13–18]. Despite these remarkable advances in coalescent-based species tree estimation, computational scalability–not statistical methodology–has become the principal bottleneck in modern phylogenomics. Contemporary phylogenomic studies increasingly seek to reconstruct species trees containing tens or even hundreds of thousands of taxa, driven by global biodiversity initiatives and ambitious Tree of Life projects [19, 20]. Although the most scalable implementation of ASTRAL, named ASTRAL-MP [17], can analyze datasets containing approximately 10,000 taxa or more than 100,000 gene trees, analyses beyond this scale remain prohibitively expensive in memory and running time even on modern high-performance computing platforms. These limitations cannot be overcome simply by adding more processors or memory. Instead, they call for fundamentally new computational frameworks for statistically consistent species tree inference.

Divide-and-conquer frameworks have substantially extended the scale of phylogenomic analyses by partitioning taxa into smaller subsets, inferring local trees, and subsequently combining them into a global phylogeny [21, 22]. A prominent example is uDance [22], which uses backbone tree expansion to incrementally analyze approximately 200,000 microbial genomes. Although scalable, uDance does not perform a single global quartet-based optimization across all taxa. Instead, scalability is achieved by decomposing the inference problem into many local analyses followed by tree assembly. Moreover, uDance is not explicitly designed to ensure statistical consistency. As a result, the long-standing scalability barrier of inferring species trees directly from unrooted gene trees by globally optimizing quartet-based criteria across all taxa at this scale has remained unresolved.

Existing coalescent-based summary methods that perform global optimization of an objective across all input taxa and are provably statistically consistent have historically been viewed as computationally prohibitive for ultra-large datasets. Our recent work on STELAR-X [23] challenged this assumption by demonstrating that rooted triplet-based, statistically consistent summary methods [24] can be redesigned to scale beyond 100,000 taxa through a fundamentally new computational framework. Rather than relying on incremental implementation optimizations, STELAR-X showed that rethinking the underlying algorithmic representations and computational workflow could dramatically reduce memory consumption and running time while preserving statistical guarantees. However, STELAR-X operates on rooted gene trees, whereas estimated gene trees are typically unrooted in practice.

ASTRAL addresses this practical setting by inferring species trees directly from unrooted gene trees with statistical consistency. Whether the computational principles underlying STELAR-X could be extended to ASTRAL, however, remained an important open question. Unlike rooted triplet-based methods, ASTRAL operates on unrooted gene trees and relies on substantially more complex optimization involving tripartition representations, quartet-weight computation, and more sophisticated constrained search-space construction. These additional algorithmic challenges make the computational techniques developed for rooted gene trees inapplicable in their existing form and require fundamentally new algorithmic ideas.

Here we present ASTRAL-X, a complete redesign of the ASTRAL framework for ultra-large species tree inference. ASTRAL-X introduces a new computational foundation for quartet-based species tree inference that preserves ASTRAL’s strong statistical guarantees and accuracy while dramatically reducing computational cost. Unlike previous improvements to ASTRAL, which focused primarily on incremental algorithmic and implementation optimizations, ASTRAL-X fundamentally redesigns the computational foundations of quartet-based species tree inference, enabling phylogenomic analyses that were previously computationally inaccessible. It introduces novel compact representations of clusters and tripartitions, scalable candidate search-space construction using Euler-tour-based lowest-common-ancestor queries, and executes workload on GPUs through a block-tiling strategy that bounds GPU memory usage. Moreover, ASTRAL-X introduces scalable procedures for constructing and scoring candidate tripartitions, including GPU-batched weight computations and complementary prefix-sum-based intersection-counting strategies. Together, these innovations extend the practical limits of statistically consistent quartet-based species tree inference to an unprecedented scale.

We evaluate ASTRAL-X on extensive simulated and empirical datasets covering a broad range of evolutionary conditions and dataset sizes. Across these benchmarks, ASTRAL-X consistently maintains species tree accuracy comparable to existing state-of-the-art methods while dramatically improving scalability. Most notably, ASTRAL-X reconstructs species trees containing up to 300,000 taxa, approximately thirty times larger than the current practical limit of ASTRAL, in under 12 hours using modest computational resources (less than 100GB of memory). It also scales efficiently with the number of input gene trees, enabling analyses with ultra-large gene collections and substantially faster and more memory efficient inference than the current methods. Furthermore, ASTRAL-X reconstructed the evolutionary history of 9,524 angiosperm species in only 16 minutes. Thus, ASTRAL-X establishes a new level of scalability for statistically consistent species tree estimation from unrooted gene trees and enables highly accurate coalescent-based phylogenomic analyses of major branches of the Tree of Life.

## 2 Material and Methods

### 2.1 Problem Definition

Let *L* be a set of *n* taxa and let = {*g*_1_, *g*_2_, …, *g*_*k*_} be a collection of *k* unrooted binary gene trees, where each gene tree *g*_*i*_ is defined on a taxon subset *L*_*i*_ ⊆ *L*. Gene trees with *L*_*i*_ ⊊ *L* are called incomplete. For any four distinct taxa *a, b, c, d* ∈ *L*, there are exactly three possible unrooted binary tree topologies on {*a, b, c, d*}, known as quartet trees: *ab cd, ac*|*bd*, and *ad*|*bc*. The topology *ab cd* signifies that the unique internal edge of the quartet tree separates {*a, b*} from {*c, d*}. For a tree *t* whose leaf set contains {*a, b, c, d*}, the induced quartet topology *q*_*t*_({*a, b, c, d*}) is the unique quartet tree consistent with *t* on those four taxa. We write *q*(*t*) for the set of all quartet trees induced by *t*.

The species tree reconstruction problem by maximizing quartet consistency seeks to find a species tree *T* on *L* that maximizes the quartet score

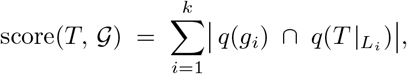

where 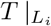 denotes the restriction of *T* to *L*_*i*_, and 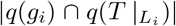 counts the number of four-taxon subsets of *L*_*i*_ whose induced quartet topology agrees between *g*_*i*_ and *T*. Under the multispecies coalescent model, this maximum quartet consistency estimator is statistically consistent [12, 14].

### 2.2 Background: Overview of ASTRAL and ASTRAL-MP

ASTRAL is a widely used summary method that estimates a species tree by maximizing the quartet consistency score defined in Section 2.1. Although the objective is expressed in terms of all induced quartet trees, ASTRAL avoids explicit quartet enumeration by using a dynamic-programming formulation over tripartitions of the taxon set [12]. Each internal node *u* of an unrooted species tree induces a tripartition *A*|*B*|*C*, obtained by removing *u* and considering the three resulting connected components. The contribution of such a tripartition can be scored independently against the input gene trees. Thus, the total quartet score of a species tree can be optimized by recursively combining locally scored tripartitions.

Since the unconstrained optimization problem is computationally intractable, ASTRAL restricts the search to a candidate set of clusters. Let *X* be a set of allowed clusters over *L*, closed under complementation, and let *w*_*G*_(*A*|*B*|*C*) denote the quartet consistency weight of a candidate tripartition with respect to the input gene trees.

Let *Y* denote the set of valid tripartitions that can be formed from the candidate cluster set *X*. That is,

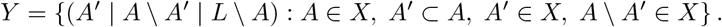

ASTRAL then uses the following dynamic-programming recurrence:

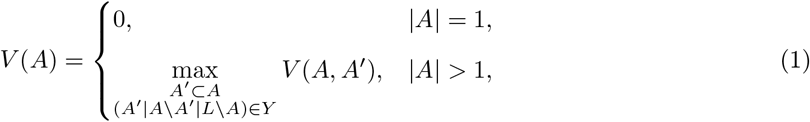

where

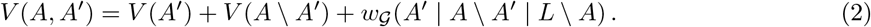

The optimal species tree is obtained by evaluating *V* (*L*) and backtracking through the maximizing choices. In the constrained setting, ASTRAL constructs *X* from bipartitions observed in the input gene trees. This constrained search remains statistically consistent under the multispecies coalescent model [12, 13].

The ASTRAL framework has been refined through several major versions including ASTRAL-II [13], and ASTRAL-III [14]. More recent implementations, including ASTRAL-MP [17] and ASTER [18], have pushed the scalability of the framework further through parallelization, randomization, and optimized engineering.

### 2.3 Overview of ASTRAL-X

ASTRAL-X is a complete redesign of the ASTRAL framework for ultra-large species tree inference. It preserves the same quartet-consistency objective, statistical guarantee, and constrained dynamic-programming formulation described above, but replaces several memory-intensive and computationally expensive components of existing ASTRAL implementations with new data structures and algorithms.

The main scalability barrier in ASTRAL-style methods comes from representing and operating on clusters and tripartitions over the full taxon set. Traditional implementations store clusters as bitsets of length n, so even a small cluster requires *O*(*n*) memory. STELAR-X [23] introduced the key idea of compact integer tuple representations for scalable coalescent-based species tree inference, dramatically reducing memory consumption in rooted triplet-based inference. ASTRAL-X extends this principle to the considerably more challenging setting of unrooted quartet-based species tree inference by representing clusters and tripartitions through compact integer tuples derived from postorder leaf arrays of the gene trees (discussed in Section 2.4).

ASTRAL-X first extracts all gene-tree clusters and tripartitions in a single bottom-up traversal of the input trees (Section 2.5). These clusters form the initial candidate set *X* for the dynamic program. To preserve the accuracy of inference, we also use parallel and optimized algorithms for enriching the cluster set *X* with heuristics to cover a broader search space [13, 14] (Section 2.6).

Since equivalent clusters and tripartitions may appear in different gene trees with different tuple encodings, ASTRAL-X uses a multi-seed double-hashing scheme to assign each cluster a permutation-invariant signature (Section 2.7). These signatures allow efficient deduplication through hash-table lookups and make hash collisions negligible in practice. They are also used to construct the DP state space efficiently.

After forming the candidate cluster set *X*, ASTRAL-X builds the valid transitions required by the dynamic program (Section 2.8). It first recovers tree-local transitions directly from the input gene tree. In contrast to STELAR-X, it also identifies additional cross-tree transitions using a GPU-accelerated hash-subtraction procedure. This preserves the richer ASTRAL search space while avoiding expensive bitset-based set comparisons.

Finally, ASTRAL-X precomputes the quartet-consistency weights of candidate tripartitions before running the dynamic program (Section 2.10). This eliminates repeated weight calculations during DP execution and offloads the main computational workload to GPU parallelization. ASTRAL-X supports four alternative methods of intersection counting, providing complementary trade-offs depending on the use-case (Section 2.10.3). The final species tree is obtained by running the ASTRAL dynamic program using table lookups for these precomputed weights.

### 2.4 Compact integer-tuple representation of clusters and tripartitions

Since ASTRAL-X evaluates quartet consistency through the clusters and tripartitions induced by the gene trees, their representation directly determines the memory requirements of the subsequent computations. ASTRAL-MP [17] represents each cluster as a bitset of length *n*, requiring *O*(*n*) memory per cluster and *O*(*n*^2^*k*) space for the *O*(*nk*) clusters across all gene trees. STELAR-X [23] showed that subtree bipartitions of rooted gene trees can instead be encoded using compact integer tuples, reducing the total storage to *O*(*nk*). Building on this insight, ASTRAL-X introduces a new tuple representation for the clusters and tripartitions of unrooted gene trees, likewise reducing the overall memory requirement to *O*(*nk*).

To construct a compact tuple representation of the clusters and partitions of the gene trees, we first root each unrooted gene tree *g*_*i*_ at an arbitrary edge. We then form the postorder traversal array *A*_*i*_ of the leaf labels *L*_*i*_ of *g*_*i*_, and let *A*_*i,l,r*_ denote the subarray of *A*_*i*_ from index *l* to index *r* inclusive (i.e., *A*_*i*_[*l* : *r*]). A fundamental property of postorder traversal is that the leaves of any subtree of *g*_*i*_ occupy a single contiguous subarray of *A*_*i*_. Consequently, for any internal node *u*, the clusters *A* and *B* comprising its left and right subtrees satisfy 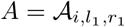 and 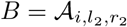 for some indices *l*_1_ ≤ *r*_1_ *< l*_2_ ≤ *r*_2_. In fact, *l*_2_ = *r*_1_ + 1 since the two subtrees are adjacent in postorder. These two clusters are thus encoded by the integer tuples (*i, l*_1_, *r*_1_) and (*i, l*_2_, *r*_2_), respectively, resembling the bipartition encoding of STELAR-X.

Unlike STELAR-X, ASTRAL-X must also represent the third cluster *C* = *L*_*i*_\(*A*∪*B*), since it operates on tripartitions of unrooted gene trees. Unlike *A* and *B*, however, *C* does not occupy a contiguous range in *A*_*i*_. We observe that *C* comprises all taxa lying outside the range [*l*_1_, *r*_2_], and so corresponds to the complement of the subarray *A*_*i*_[*l*_1_ : *r*_2_]. To accommodate this within the same framework, we augment the representation with a boolean complement flag, indicating whether a cluster corresponds to a contiguous subarray of *A*_*i*_ or to its complement. Under this convention, *C* is encoded as (*i, l*_1_, *r*_2_, 1), while *A* and *B* are encoded as (*i, l*_1_, *r*_1_, 0) and (*i, l*_2_, *r*_2_, 0), respectively. Every cluster of an unrooted gene tree is thus encoded by a 3-tuple of integers and a single boolean flag. Bipartitions and tripartitions are represented as sets of their constituent clusters. This representation encodes each cluster and tripartition in *O*(1) space, independent of the number of taxa it contains, thereby reducing the total memory complexity to *O*(*nk*). We present an illustrative example of this tuple representation in Supplementary Section 1.

### 2.5 Extraction of clusters and tripartitions

ASTRAL-X extracts all clusters and tripartitions across the gene trees with a single bottom-up traversal of each gene tree. At each non-root internal node *u* of the rooted gene tree *g*_*i*_, the traversal records two subtree clusters: the left and right child ranges encoded as (*i, l*_1_, *r*_1_, 0) and (*i, l*_2_, *r*_2_, 0), together with the complement cluster (*i, l*_1_, *r*_2_, 1) representing the remainder of *L*_*i*_. These three clusters jointly define the tripartition induced by *u*. The root node is excluded, as its two children partition *L*_*i*_ into only two non-empty parts, yielding a degenerate tripartition that contributes zero weight to ASTRAL’s quartet score. Iterating over all *k* gene trees in this manner takes *O*(*nk*) time in total and produces the complete sets of gene tree clusters *GC* and tripartitions *GT*.

### 2.6 Formation of the candidate cluster set *X*

The clusters extracted directly from the gene trees form a baseline search space for the dynamic program. When all gene trees are complete (*L*_*i*_ = *L* for all *i*), these clusters alone suffice as the initial *X*. However, when gene trees are incomplete (*L*_*i*_ ⊊ *L* for some *i*), ASTRAL-X has two alternative options available, trading computational cost for accuracy.

The first and more accurate path completes each incomplete gene tree over the full taxon set *L* and adds the clusters of the resulting trees to *X* (see Section 2.6.2). It further augments *X* with the bipartitions of a UPGMA guide tree built from the pairwise similarity matrix, following ASTRAL-MP [17]. This step relies on a precomputed *n* × *n* pairwise similarity matrix as described in Section 2.6.1.

The second, lightweight path, adds the clusters of each incomplete gene tree directly to *X* without completing it. This path is used in the tree-local mode (see Section 2.9) to avoid building the similarity matrix, thereby reducing the CPU memory requirement. In this case, however, a subtlety arises: clusters *A* ⊆ *L*_*i*_ ⊊ *L* from an incomplete gene tree are not directly reachable from the all-taxa DP root state *L* via tree-local transitions alone (as established in Section 2.10). To bridge this gap, for every cluster *A* added from an incomplete gene tree, ASTRAL-X also inserts its super-complement *L* \ *A* into *X*. The transition *L* → *A* | (*L* \ *A*) is then trivially valid from the DP root, thereby guaranteeing reachability of *A*.

We now present our method of computing the distance matrix and similarity matrix and subsequent steps to form the candidate cluster set *X*.

#### 2.6.1 GPU-accelerated distance and similarity matrix computation

##### Definitions

For a gene tree *g*_*i*_ on taxa set *L*_*i*_ ⊆ *L*, the distance *d*_*i*_(*x, y*) between two taxa *x, y* ∈ *L*_*i*_ is the number of edges on the unique *x*-*y* path in *g*_*i*_, and the similarity *s*_*i*_(*x, y*) is the number of resolved quartets of *g*_*i*_ in which *x* and *y* appear on the same side. For two taxa *x, y* ∈ *L*, let *N* (*x, y*) denote the number of gene trees containing both *x* and *y*. The aggregate distance matrix *D* and similarity matrix *S* are then the *n* × *n* matrices

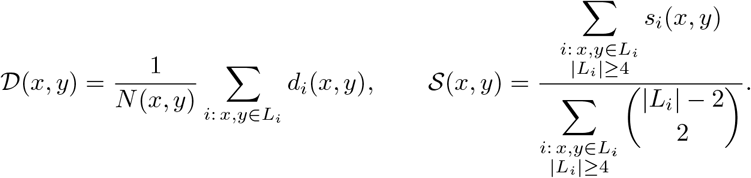

Here, the normalization in *S*(*x, y*) accounts for the fact that gene trees with different numbers of taxa contribute different numbers of quartets containing *x* and *y*. Specifically, a gene tree *g*_*i*_ with |*L*_*i*_| taxa contributes 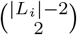 such quartets. We set *D*(*x, y*) = 0 whenever *N* (*x, y*) = 0, and *S*(*x, y*) = 0 whenever its denominator is zero. ASTRAL-MP constructs these matrices via a CPU-based multi-threaded approach, which does not scale to large *n*. We reformulate the computation for efficient GPU execution.

##### LCA-based reformulation

The efficient construction of these matrices requires careful management of repeated computation across taxon pairs. ASTRAL-MP [17] follows a bottom-up traversal of each gene tree and accumulates node-level contributions into the corresponding matrix entries. We observe that effectively leveraging GPU parallelism requires reorganizing this node-centric computation into a much larger collection of mutually independent work items. We therefore reformulate the computation using LCA-based queries so that each taxon pair can be processed in parallel. Each GPU thread computes one matrix entry by accumulating contributions across the relevant gene trees, yielding *O*(*n*^2^) independent work items that can be efficiently distributed across the available GPU threads.

The key observation is that both *d*_*i*_(*x, y*) and *s*_*i*_(*x, y*) are determined entirely by a fixed set of precomputed node-level values and the lowest common ancestor lca_*i*_(*x, y*). Concretely, storing the root-to-node distance dp(*u*) for all nodes *u* during a standard bottom-up traversal allows *d*_*i*_(*x, y*) to be recovered in *O*(1) as

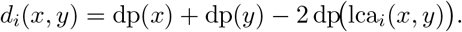

An analogous *O*(1) expression is used for *s*_*i*_(*x, y*). Its formulation is provided in the Supplementary Material (Section 3). This LCA reformulation turns the computation of each matrix entry into a single *O*(1) evaluation per (*x, y, i*) triple, once lca_*i*_(*x, y*) is known. Since these evaluations are entirely independent across all taxa pairs and all gene trees, the full *O*(*kn*^2^) workload is embarrassingly parallel and well-suited to GPU execution.

##### Euler tour and sparse RMQ for *O*(1) LCA queries

To support constant-time LCA queries on the GPU, we precompute two standard data structures per gene tree *g*_*i*_ on the CPU: an Euler tour of *g*_*i*_, recording each node at every visit during a depth-first traversal, and a sparse range minimum query (RMQ) table over the tour depths. For a gene tree *g*_*i*_ with *m*_*i*_ = |*L*_*i*_| taxa, these structures are constructed in *O*(*m*_*i*_ log *m*_*i*_) time and space and uploaded to GPU memory once, before any pairwise evaluation begins. Across all gene trees, these structures require *O*(*nk* log *n*) GPU memory. Given the first-occurrence positions of *x* and *y* in the tour, lca_*i*_(*x, y*) is the node of minimum depth in the subarray between those positions, retrieved in *O*(1) via the sparse table.

##### Block tiling for bounded GPU VRAM

Allocating the full *n* × *n* output matrix on the GPU would require *O*(*n*^2^) GPU VRAM, which becomes infeasible for large *n*. We therefore adopt a block-tiling strategy. We partition the *n* taxa into blocks of size *B* and process the matrix as *B* × *B* output tiles, iterating over all ⌈*n/B*⌉ ^2^ upper-triangular tile pairs (mirroring each tile to fill the lower triangle). For each tile, a kernel launches *B*^2^ threads, one per output cell, and each thread independently loops over all *k* gene trees to accumulate its entry. Setting

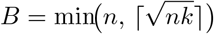

keeps the tile VRAM at *O*(*B*^2^) = *O*(*nk*). Thus, including the resident Euler-tour and sparse-RMQ structures, the total GPU VRAM required by this step is *O*(*nk* log *n* + *nk*) = *O*(*nk* log *n*). The CPU RAM requirement for the full matrix is *O*(*n*^2^). After all tiles are processed, the assembled matrices *D* and *S* reside in CPU RAM and are ready for subsequent use. Figure 2a illustrates our CPU–GPU heterogeneous approach for computing the distance and similarity matrices.

**Figure 1.**
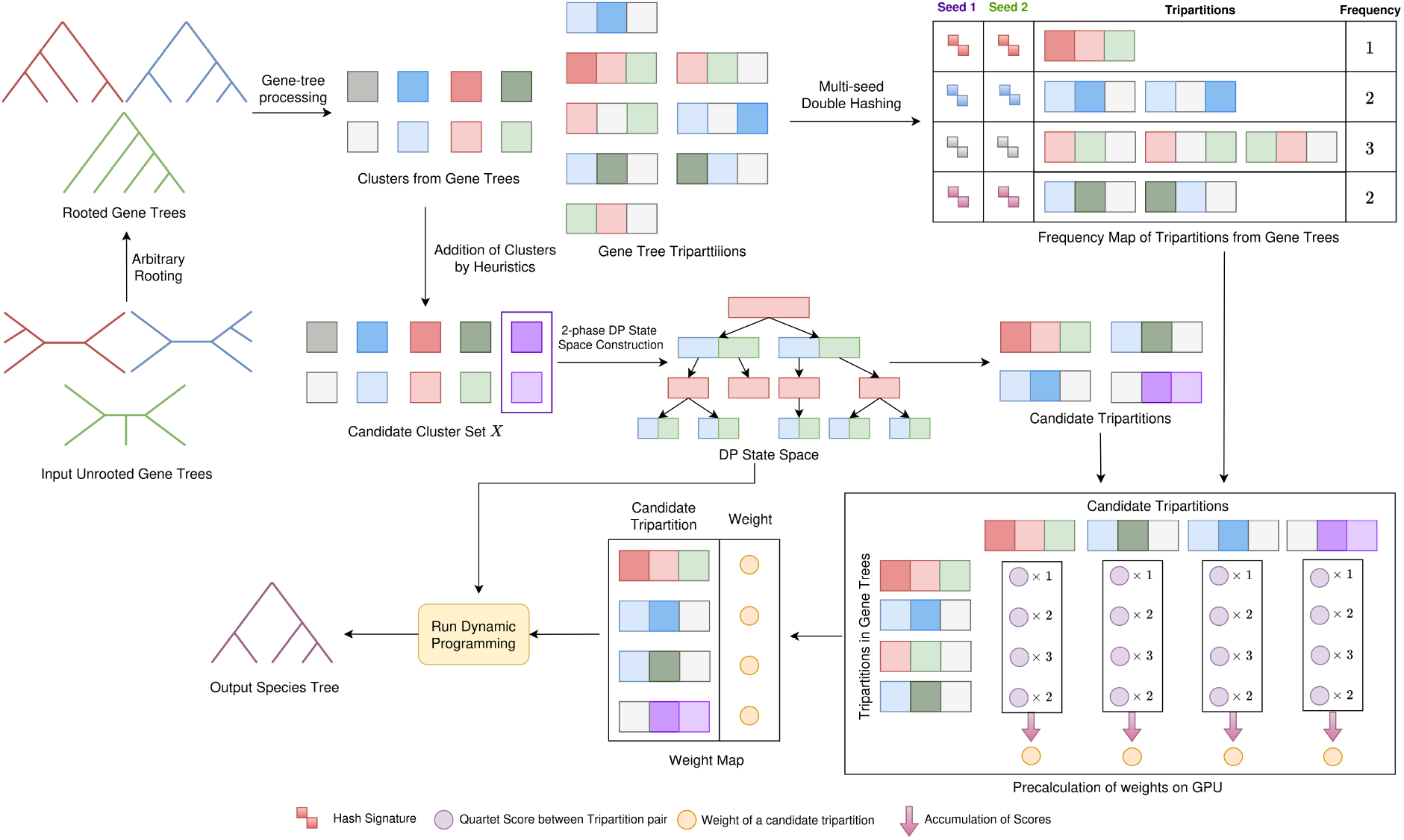
Algorithmic workflow of ASTRAL-X. Input unrooted gene trees are first arbitrarily rooted for traversal, and their clusters and tripartitions are extracted in our proposed compact representation. The candidate cluster set *X* is formed from these clusters together with heuristic additions, and is then used to construct the DP state space and candidate tripartitions. Multi-seed double hashing deduplicates gene-tree tripartitions and produces their frequency map. ASTRAL-X then precomputes candidate tripartition weights on the GPU and runs the dynamic program using this weight map to produce the output species tree.

**Figure 2.**
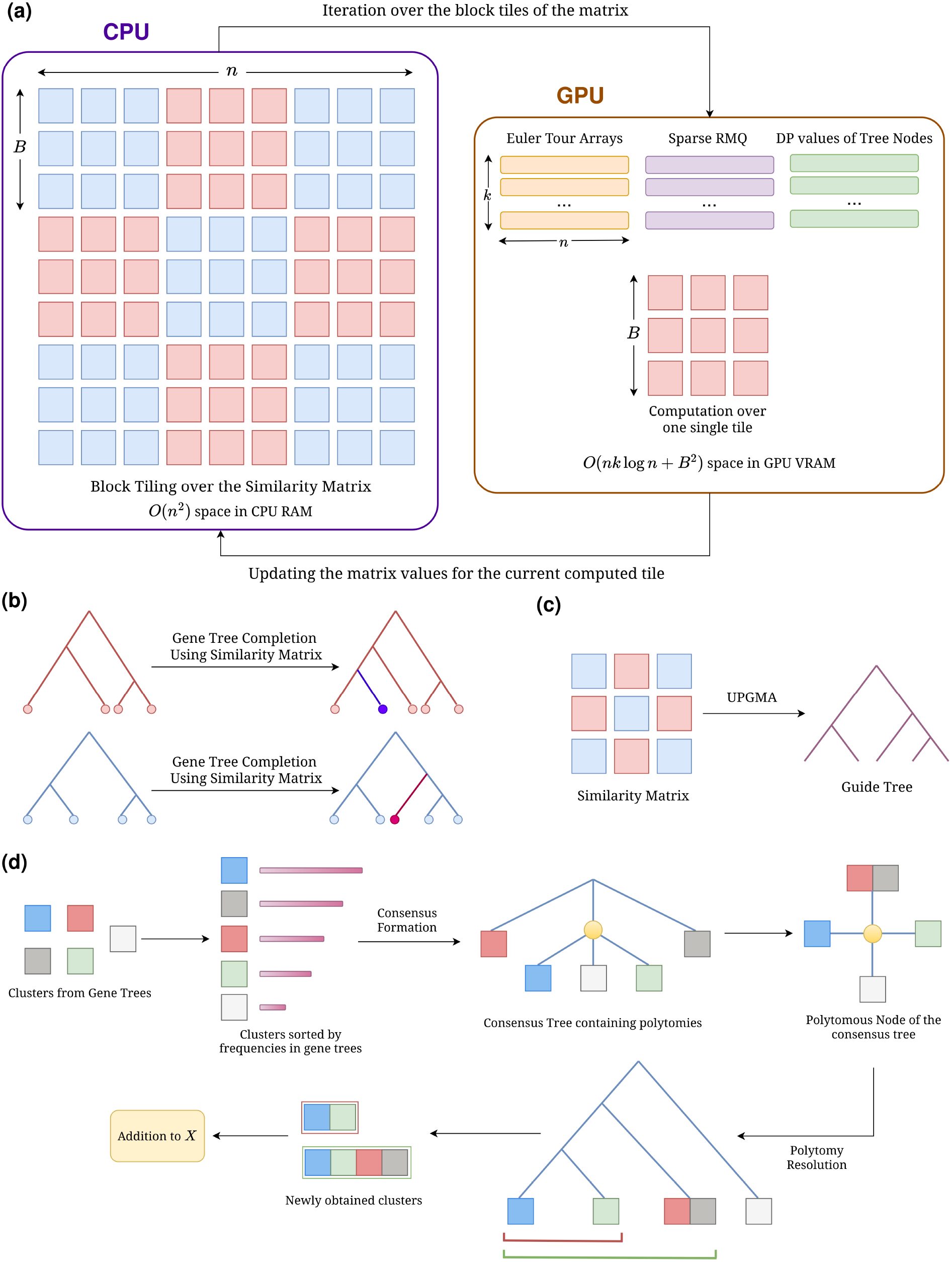
Heuristic addition of candidate clusters in ASTRAL-X. (a) GPU-accelerated computation of the pairwise similarity matrix using block tiling. The full *n* × *n* matrix is stored in CPU memory, while one *B* × *B* tile is processed at a time on the GPU. The GPU keeps the Euler tour arrays, sparse RMQ structures, and node-level DP values of the gene trees resident, and computes each tile using *O*(*nk* log *n* + *B*^2^) GPU memory. (b) The similarity matrix is used to complete the incomplete gene trees by inserting missing taxa through four-point comparisons. (c) A UPGMA guide tree is constructed from the similarity matrix, and the clusters induced by this guide tree are added to *X*. (d) Clusters observed in the gene trees are ranked by frequency and used to construct greedy consensus trees. Polytomies in these consensus trees are then resolved to generate new clusters that are added to *X*.

#### 2.6.2 Gene tree completion and addition to *X*

When gene trees are incomplete, containing only a subset *L*_*i*_ ⊊ *L* of all taxa, their clusters alone leave gaps in the candidate cluster set *X*. Following ASTRAL-MP, we complete each such gene tree by inserting every missing taxon *x* ∈ *L \ L*_*i*_ one at a time. For each *x*, we first identify its anchor: the taxon *a* ∈ *L*_*i*_ closest to *x* according to *D*, found in *O*(1) via pre-sorted distance rows. The tree is then rerooted at the edge incident to *a*, placing *x*’s likely subtree on the opposite side of the root. From there, a top-down navigation applies the four-point condition at each internal node: given the two child representatives *c*_1_, *c*_2_, the six pairwise matrix values among {*x, a, c*_1_, *c*_2_} determine whether *x* belongs in the subtree of *c*_1_, the subtree of *c*_2_, or at the current node, in *O*(1) per step. Insertion terminates when the landing position is identified. After all missing taxa are inserted, the completed gene trees are fully resolved over *L*, and their clusters are extracted and added to *X*. Figure 2b presents the completion of gene trees using similarity matrices.

#### 2.6.3 Addition to *X* from UPGMA guide tree

With the similarity matrix *S* computed as described in Section 2.6.1, we construct a UPGMA guide tree by iteratively merging the pair of active clusters with the highest average similarity, updating the similarity estimates after each merge. This runs in *O*(*n*^2^ log *n*) time and yields a fully resolved binary tree over all *n* taxa. Each of its *n* − 2 internal nodes contributes two clusters to *X*, one per side of the induced bipartition, extracted similarly as done for the gene tree clusters. The UPGMA tree enlarges the search space by capturing the averaged pairwise signal across gene trees and supplying bipartitions that may not appear in any individual gene tree. Figure 2c illustrates the expansion of the candidate search space using clusters from the UPGMA guide tree.

#### 2.6.4 Consensus-guided expansion of the cluster set *X*

To further enrich the candidate cluster set *X*, ASTRAL-X constructs a collection of greedy consensus trees from the input gene trees, following the earlier ASTRAL versions [13, 14, 17]. We first count the frequency of every cluster observed across the gene trees. Because complementary clusters describe the same edge of an unrooted tree, *A* and *L\A* are treated as equivalent during this counting. The distinct clusters are then considered in decreasing order of frequency.

Starting from a completely unresolved tree, each cluster is inserted only if it is compatible with all clusters accepted earlier. Concretely, the accepted clusters must form a laminar family. For every pair *A* and *B*,

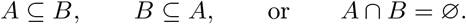

Thus, a cluster is retained when it can refine the consensus while being compatible with the current consensus hierarchy. Otherwise, it is discarded.

Maintaining compatibility efficiently is challenging because every accepted cluster changes the structure of the consensus tree. For each proposed cluster, we must identify the smallest region of the current tree containing all its taxa and verify that the cluster consists of complete child subtrees. If it cuts through an existing subtree, it is incompatible and must be rejected.

ASTRAL-MP performs this test by rebuilding a lowest-common-ancestor structure for every proposed cluster. Since each reconstruction examines the full *n*-taxon tree and at most *O*(*nk*) distinct gene-tree clusters are considered, this step requires *O*(*n*^2^*k*) time across the constant number of consensus thresholds. ASTRAL-X instead maintains the accepted clusters as an evolving laminar hierarchy and follows only the ancestor paths relevant to each proposed cluster. The total size of all clusters extracted from the *k* gene trees is *O*(*n*^2^*k*) in the worst case and *O*(*nk* log *n*) when the gene trees are balanced. Consequently, when the evolving consensus hierarchy is also reasonably balanced, the ancestor-path traversal requires *O*(*nk* log^2^ *n*) time, in addition to local scans of the affected nodes. After a successful insertion, only the corresponding part of the hierarchy is updated. Thus, ASTRAL-X avoids reconstructing a global ancestor structure for every proposal and makes the computation depend primarily on the portions of the hierarchy affected by the candidate clusters.

ASTRAL-X performs this construction incrementally and records consensus trees at the frequency thresholds

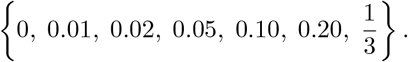

These thresholds capture compatible gene-tree signal at progressively different levels of support.

The clusters displayed directly by these consensus trees, however, are generally already present in *X*, since each accepted consensus cluster originates from a cluster observed in the input gene trees. The main value of the consensus trees, therefore, lies in their unresolved nodes or polytomies. Consider a polytomy whose incident arms contain the taxon sets *G*_1_, …, *G*_*d*_. ASTRAL-X first obtains a similarity-guided resolution by applying UPGMA to the average pairwise similarities among these arms. It also explores alternative resolutions by repeatedly sampling one representative taxon from each arm, restricting the gene trees to the sampled taxa, and constructing a local greedy consensus of the restricted trees.

A cluster recovered from such a representative consensus identifies a subset *J* ⊆ {1, …, *d*} of the polytomy arms. It is then expanded to the corresponding cluster over the original taxa,

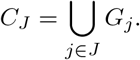

For each resulting binary resolution of the polytomy, ASTRAL-X considers all clusters induced by the resolution tree and adds their corresponding expanded clusters to *X* if they are not already present. Thus, the complete binary refinement hierarchy is represented in *X*, including the parent and sibling clusters associated with each newly introduced cluster. In this way, resolving consensus polytomies introduces new unions of well-supported groups and enlarges the candidate cluster set. This consensus-guided expansion of the cluster set *X* is illustrated in Figure 2d.

We now establish the time and memory complexity bounds of this consensus-guided cluster-expansion procedure.

##### Theorem 2.1

(Complexity of consensus-guided cluster expansion). *Suppose that ASTRAL-X uses a constant number of greedy-consensus frequency thresholds and a constant number of representative-sampling rounds for resolving each consensus polytomy. Then the consensus-guided expansion of the candidate cluster set X introduces at most O*(*n*) *additional clusters and requires O*(*nk*) *additional memory*.

*The running time of this procedure is*

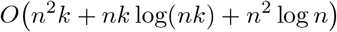

*in the worst case. When the input gene trees and the evolving consensus hierarchies have logarithmic height, the running time becomes*

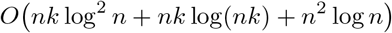

The proof is provided in the Supplementary Material (Section 2, Theorem 2.1).

### 2.7 Multi-seed double hashing for cluster and tripartition deduplication

Clusters and tripartitions extracted from different gene trees may represent identical taxon sets while having different tuple encodings. Efficiently identifying such equivalences, without explicitly traversing the underlying taxa, is therefore essential for scalability. Following STELAR-X [23], we use a permutation-invariant double-hashing scheme that maps each cluster to a compact signature, reducing equivalence testing to an expected *O*(1) hash-table lookup. Since any hashing-based deduplication scheme is susceptible to collisions, we also analyze and bound the probability of hash collisions. Unlike STELAR-X, ASTRAL-X strengthens the hash signature using multiple independent seeds, allowing the collision probability to be reduced arbitrarily by increasing the number of seeds.

#### Multi-seed double hashing

We build on the double-hashing framework introduced in STELAR-X [23]. Let *ϕ*_1_ and *ϕ*_2_ be two independent permutation-invariant and associative set-level hash functions (concretely, modular addition and bitwise XOR over 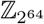, respectively). Each taxon *t* ∈ *L* carries an integer identifier id(*t*) ∈ {0, …, *n*−1}. Consider a per-element hash function ℋ that maps each identifier sparsely into 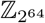, dispersing the contiguous integers 0, …, *n* − 1 across a much larger space to suppress combinatorial collisions. Given ℋ, STELAR-X defined the double-hash signature of a taxon set *S* as

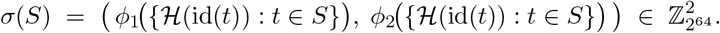

We strengthen this scheme by introducing *m* independent random seeds to further reduce the probability of hash collisions. Specifically, we replace ℋ with a seeded hash function ℋ′ that takes a seed *s* ∈ {*s*_1_, …, *s*_*m*_} alongside the taxon identifier and produces an independent sparse mapping 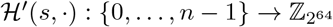 for each seed. The per-seed signature of a set *S* under seed *s* is then

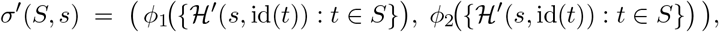

and the final multi-seed signature of *S* is the concatenation

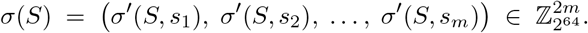

Two clusters are treated as equivalent if and only if their multi-seed signatures agree in all 2*m* components. For a tripartition (*A*|*B*|*C*), the signature is the unordered triple {*σ*(*A*), *σ*(*B*), *σ*(*C*)}. Since gene trees may carry different taxa sets *L*_*i*_, *C* = *L*_*i*_ *\A \B* is dependent on the tree of origin and not recoverable from *σ*(*A*) and *σ*(*B*) alone. Therefore, we hash all the three parts *A, B*, and *C* explicitly.

#### Prefix-scan arrays and O(1) hash evaluation

Since every cluster is either a contiguous subarray of some *A*_*i*_ or its complement, any required hash value reduces to a range query on the array *A*_*i*_. To support such queries in *O*(1), we precompute, for each gene tree *g*_*i*_ and each seed *s*, two prefix-scan arrays over *A*_*i*_:

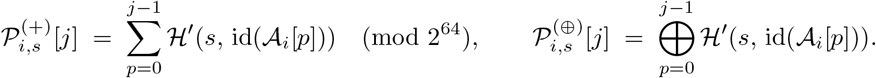

For any contiguous cluster *A*_*i,l,r*_ corresponding to the inclusive range [*l, r*], the two components of *σ*^′^(*A*_*i,l,r*_, *s*) are retrieved as

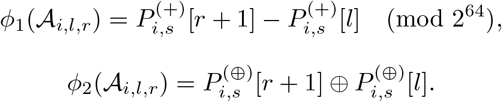

Here, we use the fact that both *ϕ*_1_ (addition) and *ϕ*_2_ (XOR) are associative and admit cancellation operators: subtraction modulo 2^64^ and XOR itself, respectively. Complement clusters also benefit from the same mechanism. For the cluster *C* = *L*_*i*_ −*A*_*i,l,r*_, represented by the tuple (*i, l, r*, 1), the hash follows by cancellation against the precomputed total 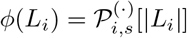:

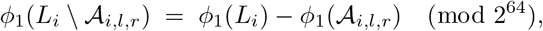

and analogously for *ϕ*_2_. Every cluster signature is thus evaluated in *O*(1) time regardless of the taxon count it represents.

#### Probability of hash collisions

The multi-seed double-hashing scheme enables us to obtain arbitrarily small collision probability by simply increasing the number of seeds. We now establish a formal bound for the collision probability by quantifying the failure probability of a single signature comparison and then apply a union bound over all clusters.

##### Theorem 2.2

(Collision Probability Bound). *Let σ*(*S*) = (*σ*^′^(*S, s*_1_), …, *σ*^′^(*S, s*_*m*_)) *be the multi-seed signature of a taxon set S, where each per-seed signature*

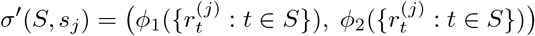

*employs two independent permutation-invariant hash functions ϕ*_1_, *ϕ*_2_ *over the modulus M, and each* 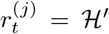 (*s*_*j*_, id(*t*)) *is independent and uniformly distributed in* ℤ_*M*_. *Among C distinct clusters, the probability that at least one pair admits an identical signature satisfies*

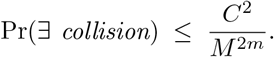

*Proof*. The central observation is that, for any two distinct sets, their signatures must agree simultaneously across all 2*m* independent hash components, an event whose probability we bound via four successive steps.

#### Setup

For each seed *s*_*j*_, the seeded hash ℋ′ maps each taxon *t* to a value 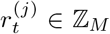, idealized as independent and uniform over ℤ_*M*_. The functions *ϕ*_1_ and *ϕ*_2_ are permutation-invariant, so *σ*(*S*) depends only on the taxon set *S*.

##### Step 1: collision probability of a single hash function

Fix a seed *s*_*j*_ and a function *ϕ*_*a*_ for *a* ∈ {1, 2}. Let *S*≠ *T*, and let *U* = *S* \ *T*, *V* = *T* \ *S*. Since *ϕ*_*a*_ is associative, it admits a cancellation operator ⊖ (subtraction for *ϕ*_1_, XOR for *ϕ*_2_), satisfying

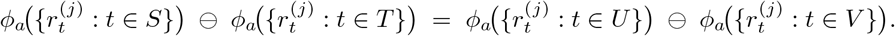

Since *S*≠ *T*, the symmetric difference *U* ∪ *V* is non-empty; let *t*^∗^ be any taxon appearing in exactly one of *U* or *V*. Conditioning on all values 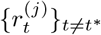, the equation above reduces to a constraint on the single uniform variable 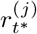. Because both addition and XOR are bijective in each argument, this constraint admits exactly one solution in ℤ_*M*_, giving

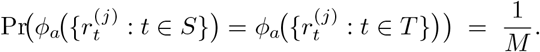

##### Step 2: collision probability of a per-seed signature

Since *ϕ*_1_ and *ϕ*_2_ are structurally independent, their collision events are independent, and

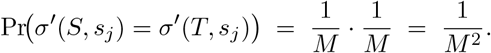

##### Step 3: collision probability of the multi-seed signature

Since distinct seeds produce independent random mappings ℋ′(*s*_*j*_, ·), the per-seed collision events are mutually independent. Multiplying across all *m* seeds,

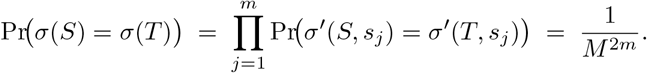

##### Step 4: union bound over all clusters

Two distinct clusters, each a single taxon set, collide precisely when their multi-seed signatures coincide, an event of probability 1*/M* ^2*m*^. Applying a union bound over all 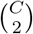 pairs,

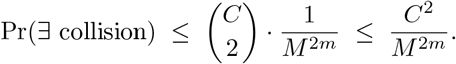

The bound extends directly to tripartitions: two distinct tripartitions collide only if all three constituent cluster signatures agree under some pairing, an event bounded above by 1*/M* ^2*m*^. Deduplicating tripartitions is therefore no less safe than deduplicating clusters.

To maintain Pr(∃ collision) ≤ *δ*, it suffices to choose *M* such that *M* ^2*m*^ ≥ *C*^2^*/δ*, equivalently 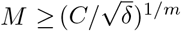. With *M* = 2^64^, *m* = 2, and *C* = |*X*| = Θ(*nk*), the denominator *M* ^2*m*^ = 2^256^ exceeds *C*^2^ = Θ(*n*^2^*k*^2^) by many orders of magnitude for any realistic dataset, rendering collisions negligible in practice. In addition, the inclusion of the seeded hash function enabled us to reduce the probability arbitrarily by simply increasing the number of seeds *m*. In our implementation, we use *m* = 2.

#### Implementation of the seeded single-element hash

We implement the seeded hash function as

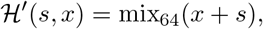

where mix_64_ is the 64-bit output-mixing function used by SplitMix [25]. It applies a short sequence of XOR-shift and odd-multiplication operations to obtain a well-dispersed 64-bit value. Adding a different seed before mixing produces a distinct seeded mapping for each component of the multi-seed signature. All calculations use unsigned 64-bit arithmetic. Consequently, additions overflow modulo 2^64^, as required by *ϕ*_1_, while *ϕ*_2_ uses the corresponding bitwise XOR operation.

#### Deduplication of clusters and tripartitions

Once the multi-seed signatures have been computed, equivalent clusters and tripartitions can be identified through hash-table lookups. Each cluster in *X* is inserted into a cluster hash table keyed by its signature, and all entries with the same signature are represented by a single canonical cluster. Applying the same procedure to the clusters and gene-tree tripartitions yields the unique sets *UX* and *UGT*. For every tripartition retained in *UGT*, we additionally store its frequency across the input gene trees. This allows repeated tripartitions to be processed only once during weight precomputation, with their contribution multiplied by the recorded frequency, as presented in Section 2.10.

### 2.8 Construction of the DP State Space

The dynamic program recurses over the cluster set *X*: at each cluster *C* ∈ *X*, it identifies all ways to split *C* into two disjoint sub-clusters *P, Q* ∈ *X* with *P* ∪ *Q* = *C*, scores the induced candidate tripartition (*P* | *Q* | *L* \ *C*), and recurses on *P* and *Q*. The collection of all such valid candidate tripartitions

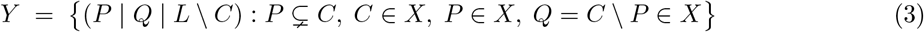

constitutes the DP state space. Since |*X*| = *O*(*nk*) by construction, the Kane–Tao bound [26] gives |*Y* | = *O* |*X*|^1.726^ = *O* (*nk*)^1.726^. Constructing *Y* efficiently is non-trivial, particularly because *X* is enriched with clusters drawn from multiple gene trees, UPGMA, and completed trees. For instance, valid splits can combine a cluster from one gene tree with a cluster from another, a cross-tree recombination that no single-tree traversal can discover.

We observe that the transitions sought by the dynamic program naturally fall into two categories. First, a split *C* → *P* | *Q* may appear directly in one of the gene trees. We refer to such a split as a tree-local transition. These transitions constitute a substantial part of the search space and can be recovered in *O*(*nk*) time through a single traversal of all input gene trees. Second, the clusters *P* and *Q* may originate from different gene trees, while their union *C* = *P* ∪ *Q* appears elsewhere in the candidate set. We refer to these as cross-tree transitions. Unlike tree-local transitions, they cannot be identified by traversing any single gene tree and therefore require a broader search across pairs of clusters from different trees. ASTRAL-MP [17] uses a randomized CPU-parallelized procedure to discover such transitions, which becomes difficult to scale to very large numbers of taxa and gene trees.

In ASTRAL-X, therefore, we adopt a two-phase approach. In the first phase, we recover all tree-local transitions directly from the input gene trees. For the second phase, we first establish that an algebraic property of the hash signatures introduced in Section 2.7 can be used to search for valid cross-tree splits with an arbitrarily small probability of hash collision. We then design a GPU-parallelized algorithm that evaluates candidate cluster pairs as independent work items distributed across GPU threads. Note, however, that the hash-subtraction algorithm can also discover tree-local transitions. Separating the computation into two phases allows ASTRAL-X, when appropriate, to restrict the DP state space to tree-local transitions only, thereby reducing the search space and improving computational efficiency. We now present these two phases of the state space construction in detail.

#### Phase 1: tree-local transitions

During the same bottom-up traversal already performed for cluster extraction (Section 2.5), we read off all transitions directly encoded in the structure of each gene tree *g*_*i*_. Two families of tree-local transitions arise (Figure 3a). For any internal node *u*, let sub(*u*) denote the cluster comprising the leaves of the subtree rooted at *u*. If *u* has children *x* and *y*, the subtree split

**Figure 3.**
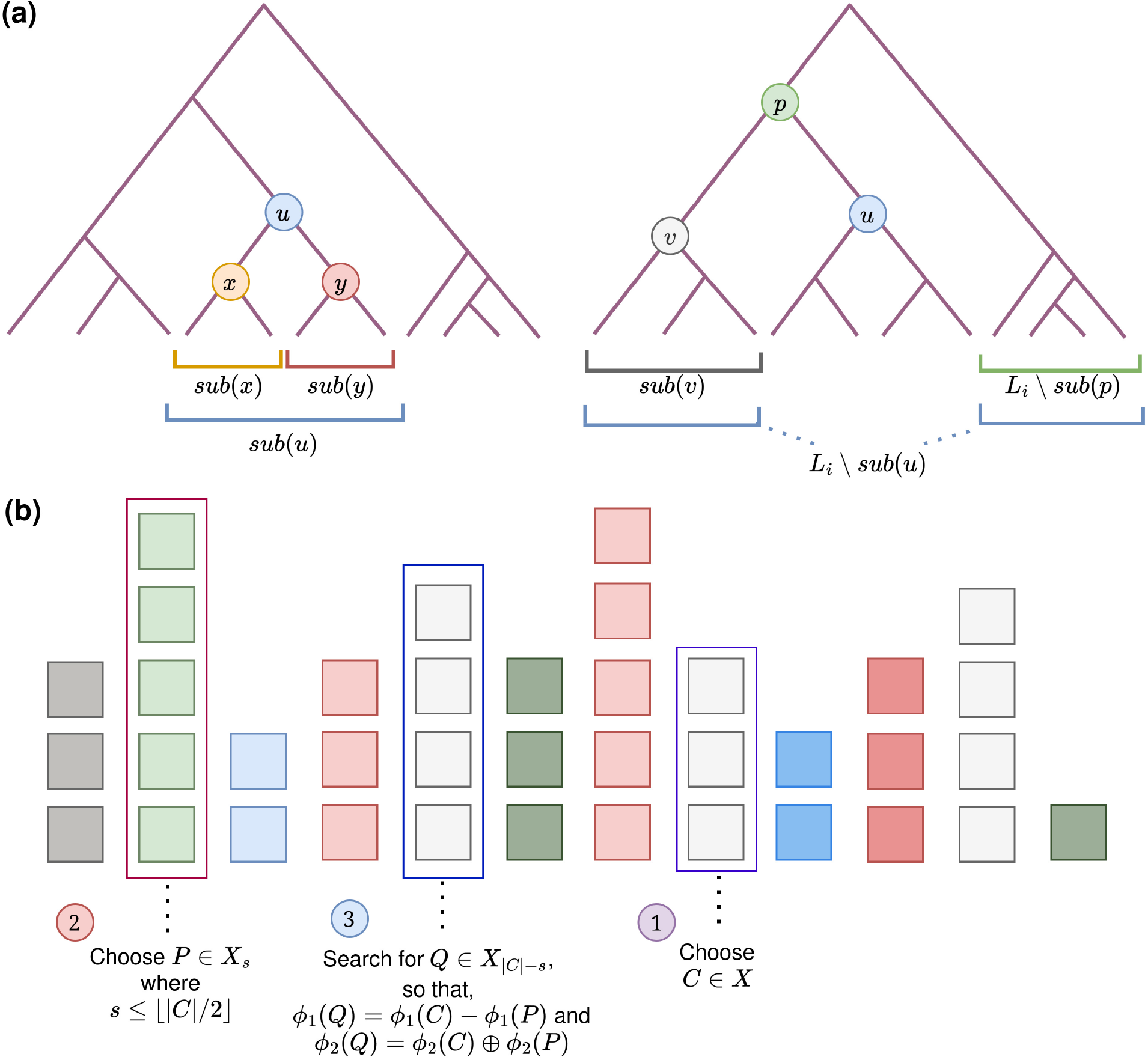
Construction of the DP state space in ASTRAL-X. (a) Tree-local transitions are recovered by direct traversal of each rooted gene tree. For an internal node *u* with children *x* and *y*, ASTRAL-X records the split sub(*u*) → sub(*x*) | sub(*y*) (left); complement-side transitions are recovered similarly from parent–sibling relationships (right) (b) Cross-tree transitions are discovered on the GPU using hash subtraction over size-binned candidate clusters. For each candidate cluster *C* ∈ *X* and each possible child cluster *P* ∈ *X*_*s*_ with *s* ≤ |*C*|*/*2, ASTRAL-X searches for the residual cluster *Q* ∈ *X*_|*C*|−*s*_ whose hash signature satisfies *σ*(*Q*) = *σ*(*C*) − *σ*(*P*), yielding the valid transition *C* → *P* | *Q*.

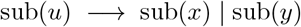

holds by construction and contributes a transition. Furthermore, for any non-root internal node *u* with parent 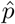 and sibling *v* (where 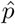 is itself not the root), the complement split

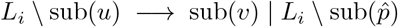

follows from the identity 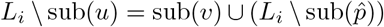. Both families are recovered during cluster extraction at no additional asymptotic cost, yielding *O*(*nk*) tree-local transitions in total. These local transitions from different gene trees are independent and are collected in parallel through multithreaded CPU execution.

#### Phase 2: cross-tree transitions via hash subtraction

Tree-local transitions may miss valid splits whose two children originate from different gene trees. In fact, the cluster set *X* is progressively expanded as described in Section 2.6. As a result, two disjoint clusters in *X* may form a valid DP transition whenever their union is also present in *X*. The second phase recovers these transitions by exploiting a fundamental algebraic property of the multi-seed hash signature *σ*.

##### Lemma 2.3

*Let P, Q, C* ⊆ *L be taxon sets with multi-seed hash signatures σ*(*P*), *σ*(*Q*), *σ*(*C*) *as defined in Section 2*.*7*.

1. **(Completeness)** *If P* ∩ *Q* = ∅ *and C* = *P* ∪ *Q, then for every seed s*_*j*_,

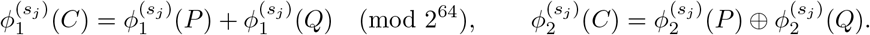
2. **(Converse)** *If the above component-wise equalities hold for all seeds s*_1_, …, *s*_*m*_ *and* |*P* |+|*Q*| = |*C*|, *then with probability at least* 1 − |*X*|^2^*/M* ^2*m*^, *we have P* ∩ *Q* = ∅ *and C* = *P* ∪ *Q*.

*Proof*. **Completeness**. For each seed *s*_*j*_, the additive hash evaluates as 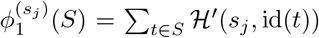 (mod 2^64^) and the XOR hash as 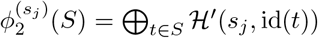. When *P* ∩ *Q* = ∅, the summation over *P* ∪ *Q* separates exactly into the sum over *P* and over *Q*, giving the first equality. A similar argument for XOR yields the second.

#### Converse

For arbitrary (possibly intersecting) *P* and *Q*,

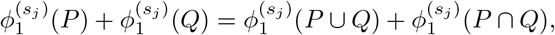

since elements in *P* ∩ *Q* are counted once in each summand on the left but only once in *P* ∪ *Q*. The analogous identity holds for 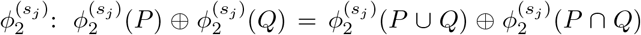. The assumed equalities therefore imply, for every seed *s*_*j*_,

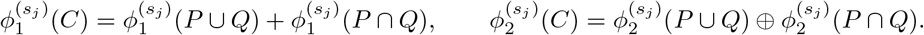

Suppose for a contradiction that *P*∩*Q* ≠ ∅ or *C* ≠ *P*∪*Q*. If *P*∩*Q* ≠ ∅, the size condition |*P*| + |*Q*| = |*C*| forces |*P*∪*Q*| *< C*, so |*C*| ≠ *P*∪*Q* in either case. The equalities above would then require the signature of *C* to agree component-wise with a specific combination of the signatures of two other distinct taxon sets, an event bounded per pair by 1*/M* ^2*m*^ by the same argument as Theorem 2.2. A union bound over all *O*(|*X*|^2^) candidate pairs yields the stated probability.

In practice, the probability of an incorrect match is negligible. Since |*X*| = *O*(*nk*), the union bound gives an error probability of *O*(*n*^2^*k*^2^)*/M* ^2*m*^. With *M* = 2^64^ and *m* = 2, the denominator becomes *M* ^2*m*^ = 2^256^, which exceeds *n*^2^*k*^2^ by many orders of magnitude for any realistic dataset. Thus, the probability of introducing a spurious transition through hash subtraction is vanishingly small.

Lemma 2.3 turns the discovery of a valid split into a constant-time residual lookup. Given a cluster *C* and a candidate subcluster *P*, we compute a residual signature *r*. For each seed *s*_*j*_, its additive component is

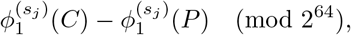

while its XOR component is

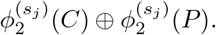

We then search for a cluster *Q* ∈ *X* such that *σ*(*Q*) = *r* and |*Q*| = |*C*| − |*P* |. A successful lookup identifies the transition *C* → *P* | *Q* with arbitrarily high confidence.

#### Algorithm and complexity

To support these lookups efficiently, we first group the clusters in *X* by size (Figure 3b). Let

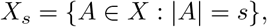

where each *X*_*s*_ is stored as a hash table keyed by cluster signature. For every *C* ∈ *X*, we consider sizes *s* ≤ ⌊|*C*|*/*2⌋. Larger values need not be examined because the two sides of a split are symmetric. For each *P* ∈ *X*_*s*_, we compute the residual signature in *O*(1) time and perform an *O*(1) lookup in *X*_|*C*|−*s*_ for a matching cluster *Q*. The total number of candidate pairs examined is

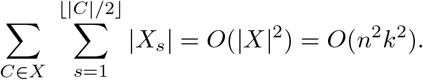

By contrast, the number of valid transitions produced is bounded by

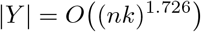

from the Kane–Tao bound [26]. Since each candidate pair can be evaluated independently, this search naturally exposes a large collection of parallel work items. The complete state-space construction procedure is summarized in Algorithm 1 and illustrated in Figure 3.

#### GPU execution

We assign each candidate pair (*C, P*) with |*P*| ≤ |*C*|*/*2 to an independent GPU thread. The thread computes the residual signature and searches the preloaded hash table for a matching cluster in *X*_|*C*|−|*P* |_. The cluster signatures and size-binned hash tables occupy *O*(*nk*) GPU memory and remain resident throughout the computation.

The output, however, requires additional care. Although the resident input data fit within the target memory budget of *O*(*nk*), the number of discovered transitions may grow to *O(*(*nk*)^1.726^), making it impractical to retain all results in GPU memory at once. We therefore process the candidate pairs in adaptive batches. Each batch launches a fixed number of GPU threads, stores the transitions found in a temporary output buffer, and transfers them to CPU memory before the next batch begins. The batch size is selected according to the available GPU memory and is reduced automatically if an allocation cannot be satisfied. Thus, the peak GPU memory usage is

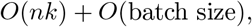

rather than depending on the total number of transitions discovered. This design allows the full cross-tree search to proceed while keeping GPU memory bounded.

### 2.9 Search-space modes of ASTRAL-X

The heuristic expansion and DP state space construction strategies described in Sections 2.6 and 2.8 allow ASTRAL-X to vary the breadth of its search space according to the characteristics of the input dataset. We therefore define three search-space modes of ASTRAL-X, ranging from a relatively constrained and efficient mode to a more extensively enlarged search space. Each successive mode retains the candidates considered by the preceding mode while introducing additional transitions in the search space.

#### Tree-local mode

The tree-local mode uses a relatively constrained search space. The candidate set *X* contains the clusters appearing directly in the input gene trees. For incomplete gene trees, it additionally includes the full-taxon super-complements *L\A* required to make the corresponding gene-tree clusters reachable from the DP root. The DP state space includes the tree-local transitions recovered directly from the input gene trees, together with the corresponding root transitions required for incomplete gene-tree clusters. This mode is particularly suitable when the number of gene trees is sufficiently large relative to the number of taxa and the extent of missing data is limited, so that the input trees themselves provide sufficient coverage of the search space.

#### Cross-tree-augmented mode

The cross-tree-augmented mode enlarges both the candidate cluster set and the DP state space. In this mode, gene trees with missing taxa are completed over the full taxon set, and the clusters obtained from the completed trees and the UPGMA guide tree are added to *X* as described in Section 2.6. The DP state space includes both tree-local and cross-tree transitions, allowing valid pairs of disjoint clusters in *X* to form a transition whenever their union is also present in *X*. This broader search space is useful when missing taxa or variation among the gene trees makes the tree-local search space inadequate, although the additional clusters and transitions increase the computational complexity.

##### Algorithm 1

Construction of the DP State Space

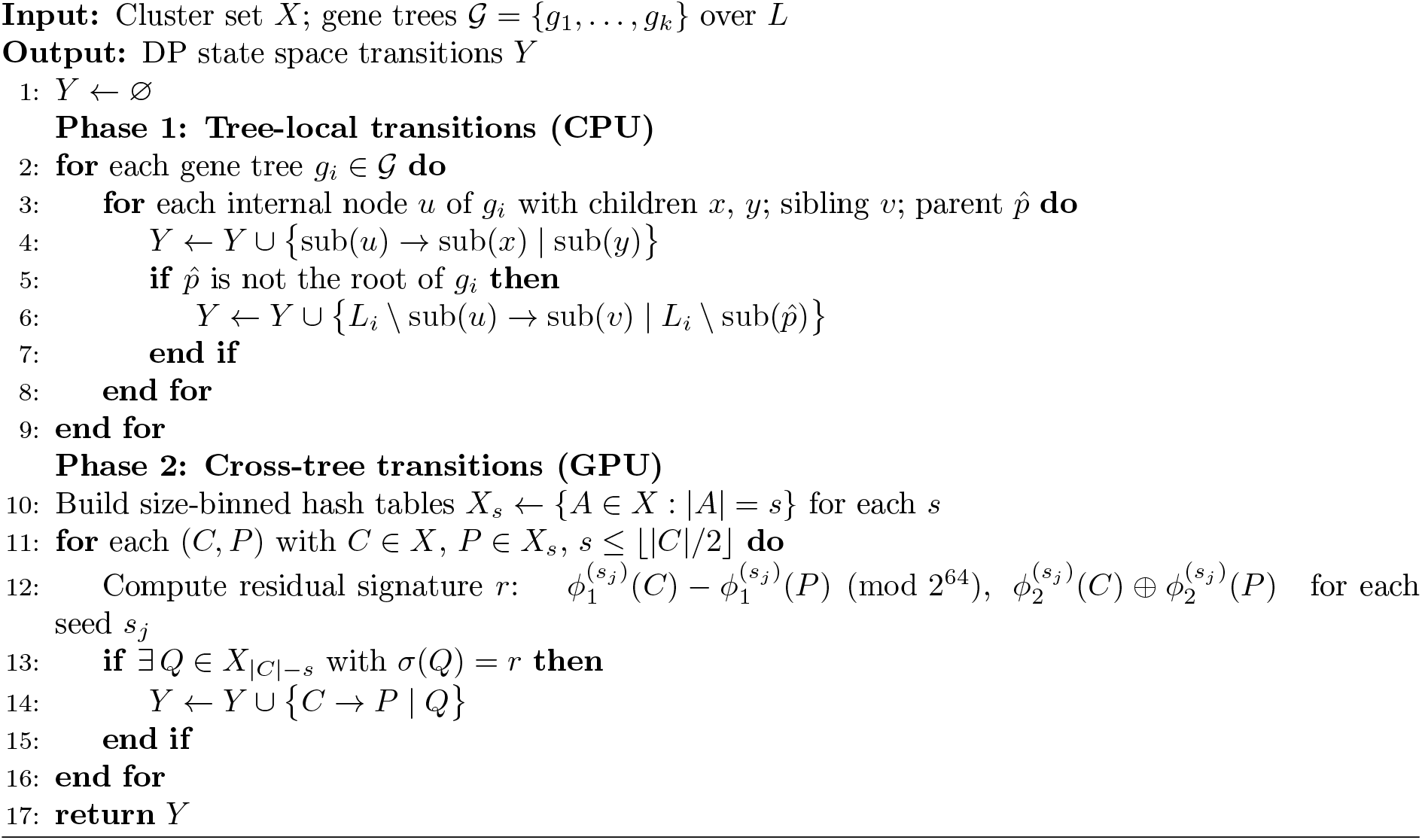

#### Expanded mode

The expanded mode provides the broadest search space in ASTRAL-X. It begins with the cross-tree-augmented formulation and further enriches *X* through the consensus-guided expansion procedure described in Section 2.6.4. The resulting cluster set *X* is then used to construct the DP state space, considering all valid transitions formed by pairs of disjoint clusters whose union is also present in *X*. This mode is appropriate when relatively few gene trees are available compared with the number of taxa, or when substantial incompleteness or discordance may leave important candidate relationships absent from the input trees. It offers the most comprehensive search space at the cost of greater running time and memory usage.

We now establish the statistical consistency of all three search-space modes of ASTRAL-X.

##### Theorem 2.4

(Statistical consistency of ASTRAL-X search modes). *Under the standard ASTRAL assumptions of complete, error-free gene trees generated under the MSC, all three search-space modes of ASTRAL-X are statistically consistent estimators of the species-tree topology*.

*Proof*. The original ASTRAL consistency argument shows that, under complete, error-free gene trees generated under the MSC, the true species-tree topology appears among the input gene trees with probability approaching one as the number of gene trees increases [12, 13]. Consequently, all tree-local transitions required to reconstruct the true species tree become available to the DP with probability approaching one. Therefore, the tree-local mode preserves the statistical consistency guarantee of ASTRAL.

Since the cross-tree-augmented and expanded modes only enlarge the tree-local search space, they retain the same statistical consistency guarantee.

### 2.10 Precomputation of tripartition weights

The dynamic program recurses over the cluster set *X*. At each cluster *C*, it evaluates every valid split *C* → *P*|*Q* by combining the recursive scores of *P* and *Q* with the weight of the induced species-tree tripartition (*P* | *Q* | *L* | *C*), and selects the split that maximises the total quartet score. Computing these tripartition weights forms the principal computational bottleneck of our workflow and determines the highest asymptotic contribution to the running time, as established later in this section. ASTRALMP [17] computes the weights on demand during the dynamic program, generating the required tasks incrementally. In ASTRAL-X, however, we observe that, as in STELAR-X [23], precomputing the weights of all candidate tripartitions introduces no unnecessary work because every such weight is eventually required by the dynamic program (Theorem 2.5). We therefore redesign the workflow to precompute all tripartition weights beforehand. This exposes a large collection of independent weight-computation tasks that can be efficiently distributed across GPU threads. We first define the quartet-consistency weight of a tripartition and establish that precomputing all weights introduces no redundant work. We then describe how these weights are evaluated efficiently.

#### 2.10.1 Quartet-consistency weight of a tripartition

Let *T* = (*P* | *Q* | *R*) be a candidate tripartition arising from the split *P* | *Q* of cluster *C* = *P* ∪ *Q*, so that *R* = *L* \ *C*. The weight of *T* counts the number of quartet trees in the input gene trees whose topology is consistent with *T* [12]. For a candidate tripartition *T* = (*P* | *Q* | *R*) and a gene tree tripartition *M* = (*M*_1_ | *M*_2_ | *M*_3_), the quartet contribution is

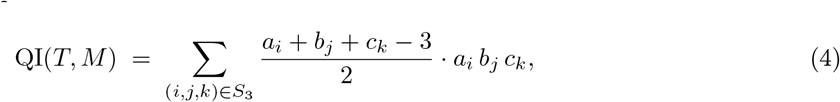

where *S*_3_ denotes the set of all six permutations of (1, 2, 3), and *a*_*i*_ = |*P*∩*M*_*i*_|, *b*_*j*_ = |*Q*∩*M*_*j*_|, *c*_*k*_ = |*R*∩*M*_*k*_|.

For a fixed assignment of the candidate sides *P*, *Q*, and *R* to the gene-tree sides *M*_*i*_, *M*_*j*_, and *M*_*k*_, respectively, a quartet consistent with both tripartitions is obtained by choosing two taxa from one of the three corresponding sides and one taxon from each of the other two. Thus, the number of such quartets is

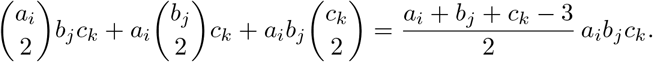

The factor 1*/*2 in Equation (4) therefore arises from the 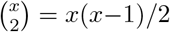 terms in this counting expression.

The total weight of *T* across all gene trees is then expressed efficiently as a weighted summation over the set *UGT* of unique gene tree tripartitions as

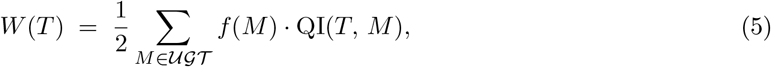

In an unrooted binary gene tree, each induced quartet is associated with the two internal nodes incident to its central edge and is therefore counted twice when the contributions are summed over all internal-node tripartitions. The outer factor of 1*/*2 in this equation corrects for this double counting across gene-tree tripartitions. Here, *f* (*M*) is the frequency of *M* across all gene trees recorded during deduplication (Section 2.7). Iterating over unique tripartitions rather than over every internal node of every gene tree helps avoid redundant computation when many gene trees share common tripartitions.

#### 2.10.2 Justification for precomputation

Precomputing the weights of all candidate tripartitions is efficient only if the dynamic program will eventually require all of them. The following theorem establishes that this is indeed the case.

##### Theorem 2.5

*The dynamic programming of ASTRAL-X eventually visits every cluster C* ∈ *X. Consequently, since the DP evaluates all valid transitions from every visited cluster, precomputing W* (*P* | *Q* | *R*) *for every valid split C* → *P* | *Q introduces no redundant computation*.

*Proof*. We show that every *C* ∈ *X* is reachable from the DP’s starting cluster *L* ∈ *X*. The DP evaluates all valid transitions at each visited cluster, so once a cluster is reached, all its children are visited in turn. We consider four cases according to the origin of *C* in *X*.

##### Case 1: *C* arises from a complete gene tree

Suppose *C* = sub(*u*) for some internal node *u* of a gene tree *g*_*i*_ with *L*_*i*_ = *L*. The cluster extraction procedure adds sub(*v*) to *X* for every internal node *v*, so all clusters along the path from *u* to the root of *g*_*i*_ lie in *X*. This path defines a chain of valid tree-local transitions (Section 2.8) from sub(root) = *L* down to *C*. Since the DP visits *L* and evaluates all valid transitions at each step, it descends this chain and visits *C*.

##### Case 2: *C* arises from the UPGMA guide tree

The UPGMA tree is a fully-resolved tree constructed over the full taxon set *L*, so its root cluster is *L* itself. The same chain argument of Case 1 applies verbatim, with the UPGMA tree replacing the gene tree.

##### Case 3: *C* arises from an incomplete gene tree

Suppose *C* = sub(*u*) for some internal node *u* of a gene tree *g*_*i*_ with *L*_*i*_ ⊊ *L*. The extraction procedure adds, for each internal node of such a tree, not only the subtree cluster *A* = sub(*u*) ⊆ *L*_*i*_ but also its super-complement *L* \ *A* to *X*, where the complement is taken with respect to the full taxon set *L* rather than *L*_*i*_ alone. Since *A* ∪ (*L* \ *A*) = *L* and *A* ∩ (*L* \ *A*) = ∅, with both *A* and *L* \ *A* in *X*, the transition

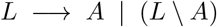

is a valid split from the DP root. The DP therefore reaches *A* = *C* directly in one step from *L*.

##### Case 4: *C* arises from consensus-guided polytomy resolution

Consider a polytomy in a greedy consensus tree whose incident arms have taxon sets *G*_1_, …, *G*_*d*_, and let

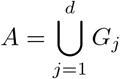

denote the cluster corresponding to the polytomy. The consensus-guided expansion replaces this polytomy by a binary refinement of its *d* arms (Section 2.6.4). Every cluster introduced by this refinement is therefore a union

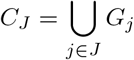

for some subset *J* ⊆ {1, …, *d*}.

ASTRAL-X adds the clusters induced by the complete binary refinement to *X*. Hence, for every non-root cluster *C*_*J*_ in this refinement, its parent cluster *C*_*J*_′ and its sibling cluster *C*_*J*_′_\*J*_ are also in *X*, and

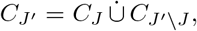

where 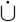 denotes disjoint union. Therefore,

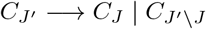

is a valid DP transition.

The cluster *A* at the root of the local refinement is already present in *X*: it is either the full taxon set *L* or a cluster of the greedy consensus hierarchy, and every such consensus cluster originates from a gene-tree cluster already included in *X*. By Cases 1 and 3, *A* is reachable from *L*. Following the transitions of the binary refinement from *A* downward therefore reaches every cluster introduced by the consensus-guided resolution, including *C*.

In all four cases, *C* is reachable from *L*. Therefore the DP visits every cluster in *X*, and every weight the precomputation produces is consumed.

#### 2.10.3 Computation of the intersection counts

Equations (4) and (5) show that computing quartet-consistency weights primarily requires evaluating intersection sizes between clusters belonging to candidate and gene-tree tripartitions. We first show that only a small subset of these intersection counts must be computed directly, while the remaining values can be derived from simple counting identities. Computing even these core intersections, however, is non-trivial under our compact tuple representation. In ASTRAL-MP [17], clusters are stored as length - *n* bitsets, so an intersection size can be easily obtained through a bitwise intersection followed by a population count in *O*(*n*) time. In ASTRAL-X, clusters are instead represented by integer tuples denoting ranges, or complements of ranges, in gene-tree traversal arrays (Section 2.4). A direct approach would therefore require iterating over the taxa represented by each pair of clusters, substantially increasing the running time. Therefore, later in this section and in Section 2.10.4, we develop efficient procedures for computing the required intersection counts while retaining the running time and memory advantages of the compact representation.

Equation (4) depends on the nine entries of a 3 × 3 intersection matrix, where the (*r, c*)-th entry records the number of taxa shared between the *r*-th side of the candidate tripartition (*P, Q, R*) and the *c*-th side of the gene-tree tripartition (*M*_1_, *M*_2_, *M*_3_) (Figure 4). It is unnecessary, however, to compute all nine entries independently. Four carefully chosen intersection counts are sufficient, and the remaining five can be recovered from the row and column sums of the matrix.

**Figure 4.**
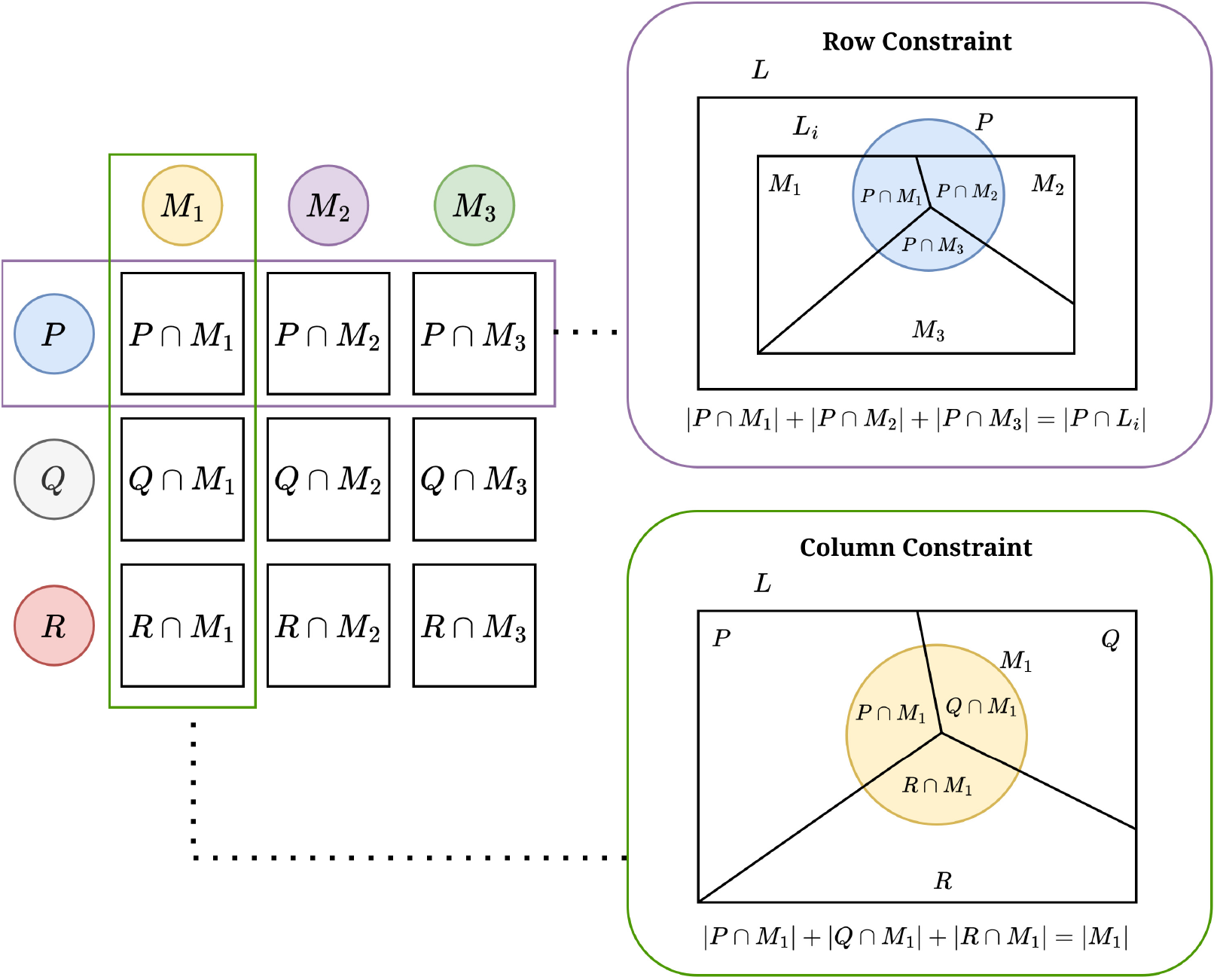
Computation of the 3 × 3 intersection matrix. Only four core intersection counts are evaluated directly. The remaining five are recovered from the row and column constraints induced by the candidate tripartition (*P* | *Q* | *R*) and the gene-tree tripartition (*M*_1_ | *M*_2_ | *M*_3_).

We first use the column sums (Figure 4). Since *P*, *Q*, and *R* form a partition of the full taxon set *L*, every taxon in *M*_*j*_ ⊆ *L*_*i*_ ⊆ *L* belongs to exactly one of these three candidate sides. Therefore,

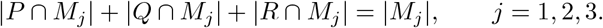

Let

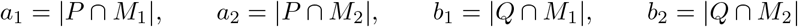

denote the four core intersection counts. The column constraints then immediately give the first two entries of the third row:

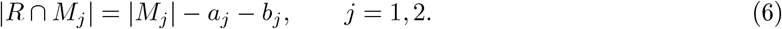

We next use the row constraints (Figure 4). The gene tree sides *M*_1_, *M*_2_, and *M*_3_ partition *L*_*i*_, rather than the full taxon set *L*. Thus, a candidate side such as *P* may contain taxa absent from gene tree *g*_*i*_, and its row sum is |*P* ∩ *L*_*i*_| rather than |*P* |. Hence,

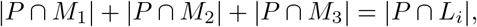

with an analogous identity for *Q*. Rearranging yields the remaining entries in the first two rows:

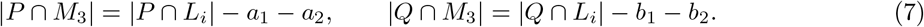

Finally, the last unknown entry follows from the column sum for *M*_3_:

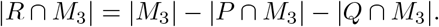

Thus, evaluating the four core intersections, together with the two row totals |*P* ∩ *L*_*i*_ | and |*Q* ∩ *L*_*i*_|, determines the complete intersection matrix.

The two row totals |*P* ∩ *L*_*i*_| and |*Q* ∩ *L*_*i*_ | depend only on the candidate sides *P* and *Q* and on the leaf set *L*_*i*_ of gene tree *g*_*i*_. They therefore need to be obtained only once for each candidate–gene-tree pair and can then be reused for all tripartitions induced by the internal nodes of *g*_*i*_. For complete gene trees, these quantities reduce directly to |*P*| and |*Q*|. For incomplete gene trees, they count only the taxa of *P* and *Q* that are present in *L*_*i*_. The four different intersection-computation methods obtain these row totals using their respective representations and computational procedures as described later in this section.

Computing the four core intersection counts between each candidate tripartition and each unique gene tree tripartition is therefore the central computational task. We now present four complementary methods of computing the intersection counts (Section 2.10.4, Section 2.10.5, Section 2.10.6, and Section 2.10.7) that are implemented in ASTRAL-X and subsequently discuss when each is preferable (Section 2.10.8). Importantly, all these four methods are mathematically equivalent and differ only in how the required computations are organized, leading to different running-time and memory requirements. We also present an example showing the intersection computation using the four methods in Supplementary Material (Section 1).

#### 2.10.4 Intersection computation via smaller-side traversal

As the first method, we adopt the intersection counting approach introduced in STELAR-X [23]. For two clusters 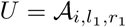 and 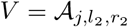 from distinct gene trees, the intersection |*U*∩*V*| is computed by iterating over the smaller cluster and checking, for each taxon *t*, whether the inverse index *π*_*j*_(*t*) falls within [*l*_2_, *r*_2_]. Assume without loss of generality that |*U* | ≤ |*V* |.

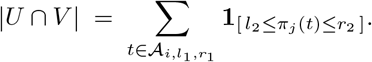

Each such count costs *O*(min(|*U*|, |*V*|)).

The same procedure also handles complement-flagged clusters. If 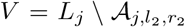, membership is determined by inverting the interval test: a taxon *t* belongs to *V* if it is present in *L*_*j*_ and *π*_*j*_(*t*) */*∈ [*l*_2_, *r*_2_]. If the cluster selected for traversal is complement-flagged, its taxa are enumerated from the two ranges outside its encoded interval. The total number of taxa traversed still equals the size of that cluster, so the *O*(min(|*U*|, |*V*|)) bound is unchanged.

Since the weight formula (Equation (5)) sums over the deduplicated set *UGT*, this method iterates over |*UGT* | unique gene tree tripartitions per candidate tripartition, exploiting cross-tree deduplication whenever gene trees share common tripartitions.

We now analyze the complexity of this approach in our setting of ASTRAL-X.

##### Theorem 2.6

(Smaller-side precomputation complexity). *Let Y be the set of valid candidate tripartitions with* |*Y* | ≤ *O* (*nk*)^1.726^, *and let UGT be the set of unique gene tree tripartitions with* 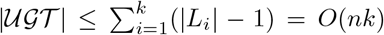. *Denoting the two explicit sides of tripartition y* ∈ *Y by P*_*y*_ *and Q*_*y*_, *the total precomputation cost of the smaller-side method is*

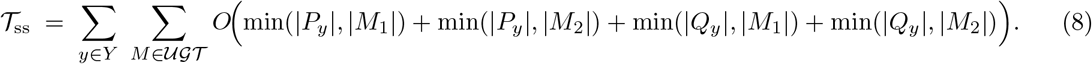

*This satisfies the following bounds:*

- *Tree-local transitions only (*|*Y* |_local_ = *O*(*nk*)*), balanced gene trees: T*_ss_ = *O*(*n*^2^*k*^2^).
- *Tree-local transitions only (*|*Y* |_local_ = *O*(*nk*)*), caterpillar gene trees: T*_ss_ = *O*(*n*^3^*k*^2^).
- *Full set of transitions including cross-tree ones (Y* = *O* (*nk*)^1.726^ *): T*_ss_ = *O(*(*nk*)^1.726^ · *n*^2^*k) in the worst case*.

*Proof*. **Cases 1 and 2 (tree-local)**. With |*Y*|_local_ = *O*(*nk*), both candidate tripartitions and gene tree tripartition clusters arise from the same gene tree family. Since tripartitions are unordered, the ordinary subtree transition and the corresponding complement-side transition induce the same tripartition up to a permutation of its three sides. Thus, after deduplication, each tree-local candidate tripartition can be represented using the two child-subtree sides of its inducing internal node. Thus, complement-side transitions do not introduce additional tripartition-weight computations.

The double sum in (8) decomposes over ordered pairs of gene trees (*g*_*p*_, *g*_*q*_): for each internal node *u* of *g*_*p*_ contributing candidate tripartition *y* with child-subtree sizes |*P*_*y*_| and |*Q*_*y*_|, and each internal node *v* of *g*_*q*_ contributing tripartition cluster sizes |*M*_1_| and |*M*_2_|, the cost of the corresponding term is *O*(min(|*P*_*y*_|, |*M*_1_|) + min(|*P*_*y*_|, |*M*_2_|) + min(|*Q*_*y*_|, |*M*_1_|) + min(|*Q*_*y*_|, |*M*_2_|)).

For balanced gene trees, subtree sizes take values in *S* = {*n/*2, *n/*4, …, 1} with *n/*(2*s*) nodes of child-size *s* per tree. Under this child-subtree representation, for balanced nodes |*P*_*y*_| = |*Q*_*y*_| = *s* and |*M*_1_| = |*M*_2_| = *t*, so the per-pair cost is *O*(min(*s, t*)). Summing over all node pairs in a single tree pair (*g*_*p*_, *g*_*q*_):

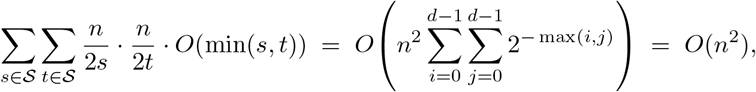

where the double sum is bounded by the same convergent series as in STELAR-X [23]. Summing over all *k*^2^ ordered tree pairs gives *O*(*n*^2^*k*^2^).

For caterpillar gene trees, internal node *u* has child-subtree sizes 1 and |sub(*u*) | − 1, with |sub(*u*) | ranging over 2, …, *n*. The per-pair cost is *O*(min(*s, t*)) where *s* = |sub(*u*) | and *t* = |sub(*v*) |. Summing over all node pairs in one tree pair:

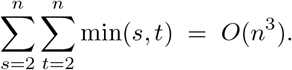

Over *k*^2^ tree pairs: *O*(*n*^3^*k*^2^).

##### Case 3 (full cross-tree)

For an arbitrary *y* ∈ *Y*, we bound the inner sum in (8) independently of the sizes |*P*_*y*_| and |*Q*_*y*_| using min(*a, b*) ≤ *b*:

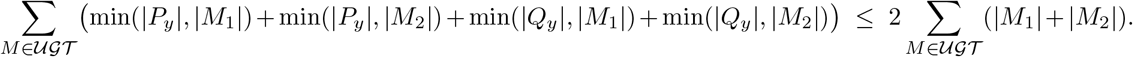

Since |*M*_1_| + |*M*_2_| = |sub(*u*)| for the node *u* inducing *M* :

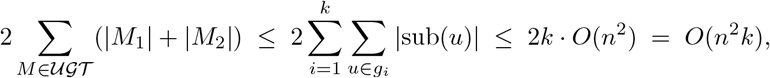

where ∑_*u*_ |sub(*u*)| = *O*(*n*^2^) is the worst-case (caterpillar) total subtree mass per tree. This bound is uniform across all *y* ∈ *Y*, so

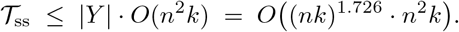

The topology-dependence in Cases 1 and 2 is a limitation of this method. While the balanced-tree bound *O*(*n*^2^*k*^2^) is very efficient in practice, the caterpillar worst case can be significant on highly unbalanced inputs. The prefix-sum method below removes this topology dependence entirely.

#### 2.10.5 Intersection computation via prefix sums

By the tuple representation described earlier, we know that a gene tree tripartition *M* = (*M*_1_ |*M*_2_| *M*_3_) arising from internal node *u* of *g*_*i*_ corresponds to either contiguous subarrays or complement of a subarray of *A*_*i*_ as follows:

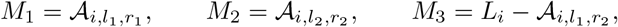

for some indices *l*_1_ ≤ *r*_1_ *< l*_2_ ≤ *r*_2_. Computing |*P*_*y*_ ∩ *M*_1_| therefore reduces to counting how many taxa of *P*_*y*_ occupy positions [*l*_1_, *r*_1_] in A_*i*_, a range counting query that admits an *O*(1) computation via prefix sums after an *O*(|*L*_*i*_|) preprocessing.

##### Prefix-sum construction

Let us fix a candidate tripartition *y* = (*P*_*y*_ | *Q*_*y*_ | *R*_*y*_) ∈ *Y* and a gene tree *g*_*i*_ with postorder array *A*_*i*_ of length |*L*_*i*_|. Define the leaf membership indicators for *P*_*y*_ and *Q*_*y*_ over *g*_*i*_’s leaf positions:

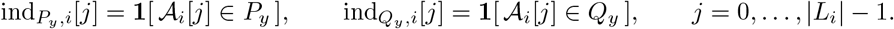

Thus, 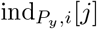 is a boolean indicator variable stating whether the *j*-th taxon in *g*_*i*_ belongs to *P*_*y*_. We may easily build this indicator array by traversing the taxa in *P*_*y*_ and setting the corresponding indices to 1 using the inverse-index lookup. Next, we build the prefix-sum arrays 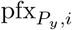 and 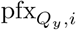, where

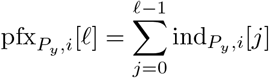

counts taxa of *P*_*y*_ in positions 0 through ℓ − 1 of _*i*_, and 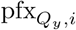 is defined identically for 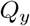. This construction takes *O*(*n*) time.

Every core intersection count for every internal node of *g*_*i*_ is then retrieved in *O*(1):

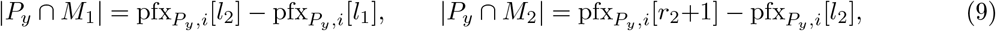

where the first formula exploits the adjacency *l*_2_ = *r*_1_ + 1 directly. The identities for *Q*_*y*_ follow symmetrically. Importantly 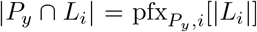 and 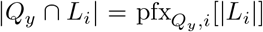, so the row sums needed for Equations (7) also emerge as by-products of the prefix arrays, handling incomplete gene trees at no extra cost under this setting.

###### Theorem 2.7

(Prefix-sum precomputation complexity). *Using the prefix-sum formulation, the total weight precomputation time is*

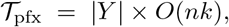

*unconditionally independent of gene tree topology. For the two modes of DP state space construction:*

1. *Tree-local transitions only (*|*Y* |_local_ = *O*(*nk*)*): T*_pfx_ = *O*(*n*^2^*k*^2^).
2. *Full set of transitions including cross-tree ones (*|*Y* | = *O* ((*nk*)^1.726^)): *T*_pfx_ = *O*((*nk*)^1.726^ · *nk*) = *O*((*nk*)^2.726^).

*Proof*. For each candidate tripartition *y* ∈ *Y* and each gene tree *g*_*i*_, building the two prefix arrays 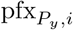 and 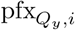 over _*i*_ costs *O*(|*L*_*i*_|). Each of the |*L*_*i*_| −1 internal nodes of *g*_*i*_ then contributes four *O*(1) prefix-difference lookups together with a constant-time QI evaluation. The per-(*y, g*_*i*_) cost is therefore *O*(|*L*_*i*_|) exactly, regardless of tree topology. Summing first over all *k* gene trees and then over all |*Y*| candidate tripartitions:

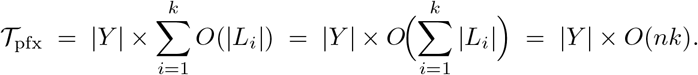

Substituting |*Y* |_local_ = *O*(*nk*) and |*Y* | = *O*((*nk*)^2.726^) yields Cases 1 and 2 respectively.

##### GPU execution via shared memory

Since the prefix-sum computation for each *y* ∈ *Y* is independent of every other, the workload is embarrassingly parallel. One GPU thread-block is assigned per candidate tripartition *y*. The block’s threads cooperate to build 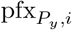 and 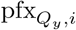 for each gene tree *g*_*i*_ in sequence via a parallel prefix scan in shared memory, evaluate all |*L*_*i*_| −1 internal nodes in a striped loop, accumulate partial QI scores per thread, and reduce to a single output value. The two prefix arrays occupy *O*(*n*) entries of shared memory per block and are discarded after each gene tree, so the global VRAM footprint scales with the number of active blocks, not with |*Y*|. For gene trees too large to fit both arrays in shared memory, these arrays are stored in the resident global-memory.

#### 2.10.6 Intersection computation by postorder propagation

ASTRAL-X also provides a direct tree-walk method for computing the intersection counts required by the quartet-scoring function. Rather than processing each gene-tree tripartition independently, this method exploits the nested structure of the tree: the counts associated with an internal node are obtained by combining the information already computed for its children.

Fix a candidate tripartition *y* = (*P*_*y*_ | *Q*_*y*_ | *R*_*y*_) and a gene tree *g*_*i*_ with leaf set *L*_*i*_. For each node *u* of *g*_*i*_, let *L*_*i*_(*u*) denote the taxa below *u*. During a postorder traversal, we maintain the state

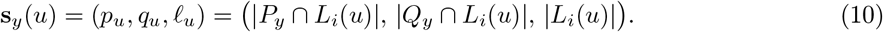

A separate count for *R*_*y*_ is unnecessary because the three candidate sides partition the full taxon set, giving

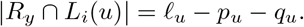

At a leaf labelled by taxon *x*, the state records whether *x* belongs to *P*_*y*_ and *Q*_*y*_, together with the subtree size 1. In this method, these membership queries are answered from bitset representations of the candidate clusters.

Now consider a non-root internal node *u* with children *v* and *w*. Its two child subtrees form the first two sides of the induced gene-tree tripartition, while the taxa outside *L*_*i*_(*u*) form the third. The four core intersections are therefore available directly as *p*_*v*_, *p*_*w*_, *q*_*v*_, and *q*_*w*_. Intersections with *R*_*y*_ follow from the child-subtree sizes. For example,

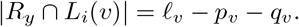

The intersections with the outside side are obtained from the row totals

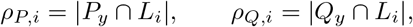

by subtracting the contributions of the two child subtrees. Thus, once the child states are known, all intersection counts needed to score the tripartition at *u* follow through constant-time arithmetic.

The state propagated to the parent is simply

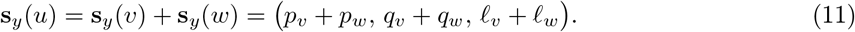

Starting from the leaves, this recurrence therefore recovers the required intersection information exactly at every internal node.

For a complete gene tree, the row totals reduce to *ρ*_*P,i*_ = |*P*_*y*_| and *ρ*_*Q,i*_ = |*Q*_*y*_|. For an incomplete gene tree, they are computed once for the candidate–gene-tree pair by intersecting the candidate sides with the set of taxa present in *g*_*i*_, and are then reused at every internal node of that tree.

ASTRAL-X stores the gene trees as compact postorder traversal streams. During evaluation, leaf states are pushed onto a stack, while each internal node combines the states of its children, scores the induced partition, and pushes the resulting state upward. This directly accumulates the contribution of every internal-node occurrence. Consequently, repeated gene-tree partitions need not be deduplicated or stored in a separate frequency table.

Let

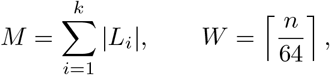

and let *k*_inc_ denote the number of incomplete gene trees. For one candidate tripartition, traversing all gene trees requires *O*(*M*) time, while computing the row totals for incomplete trees contributes *O*(*k*_inc_*W*) word operations. The per-candidate running time is therefore

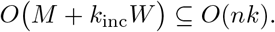

Hence, scoring all candidate tripartitions requires

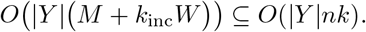

The resulting bound depends on the search-space mode. In tree-local mode, the DP state space contains only transitions occurring directly in the input gene trees, and hence |*Y*| = *O*(*nk*). The total running time for scoring all candidate tripartitions is therefore

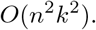

In both the cross-tree-augmented and expanded modes, the candidate cluster set satisfies |*X*| = *O*(*nk*), while cross-tree transitions are considered among clusters in *X*. Using the bound |*Y*| = *O*(|*X*| ^1.726^), the number of candidate tripartitions is at most *O*((*nk*)^1.726^) in either mode. Consequently, the corresponding scoring time is

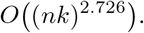

The expanded mode generally contains more candidate clusters and transitions than the cross-tree-augmented mode in practice, but both retain the same worst-case asymptotic bound.

The resident postorder representation of the gene trees requires *O*(*nk*) memory, while the shared candidate-cluster bitsets require

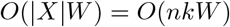

additional machine words, where *W* = ⌈*n/*64⌉. To bound the candidate-dependent working memory, ASTRAL-X processes candidate tripartitions in batches. Let *J* denote the number of candidates processed concurrently in a batch. Each active candidate maintains a private postorder stack whose size is determined by the maximum traversal frontier *H* of the gene trees. Thus, processing *J* candidates concurrently requires

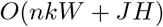

GPU memory. Since candidate tripartitions are scored independently, changing the batch size affects only the degree of parallelism and the peak memory usage, not the resulting quartet scores.

#### 2.10.7 Intersection computation via bitwise operations

Although the compact tuple representation is essential for large taxon sets, dense bitset representations can be more efficient when *n* is small. A bitset over the full taxon set requires 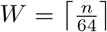 64-bit words. Thus, when *n* spans only a few machine words, the memory overhead remains modest and intersections can be evaluated using fast word-level bitwise operations.

In this method, each distinct candidate cluster is materialized once as a bitset over the global taxon set and shared by all candidate tripartitions that use it. Gene-tree partitions are stored similarly, together with their frequencies. For a binary gene-tree tripartition (*M*_1_ | *M*_2_ | *M*_3_), only the first two sides are stored explicitly while the third is recovered from the corresponding row and column totals.

For two clusters *P* and *Q* represented as bitsets, the intersection size |*P* ∩ *Q*| is obtained by taking their bitwise intersection and counting the set bits in the result. The four core intersection counts are computed in this way, while the remaining entries follow from the row and column constraints described in Section 2.10.3.

Each candidate cluster in *X* is materialized only once, so the candidate-cluster bitsets require *O*(*W*|*X*|) machine words. Across all *k* gene trees, the total number of explicitly stored partition sides is *O*(*nk*), contributing an additional *O*(*Wnk*) words. Since |*X*| = *O*(*nk*), the overall memory requirement becomes

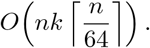

To score a candidate tripartition, ASTRAL-X computes its intersections with the *O*(*nk*) partition sides induced by the input gene trees. Each intersection examines *W* = ⌈*n/*64⌉ machine words, and therefore the total running time for scoring all candidate tripartitions is

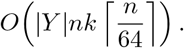

This bitset-based approach is consequently most effective when the taxon set occupies only a small number of machine words. As *n* increases, the additional factor ⌈*n/*64⌉ becomes increasingly costly in both time and memory, and ASTRAL-X instead uses the compact tuple-based methods.

Candidate tripartitions are scored independently and can be distributed across GPU threads. The shared bitset pools mentioned above remain resident in memory, while each worker maintains only the current intersection counts and accumulated weight. This yields a simple and highly efficient approach for computing intersection counts for datasets with relatively small numbers of taxa.

#### 2.10.8 Choice of the intersection computation method

The four intersection-counting methods described above compute the same quartet-consistency weights, but they present different trade-offs in running time, memory usage, and sensitivity to the input. Let

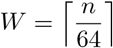

denote the number of machine words in a taxon bitset. In tree-local mode, |*Y* | = *O*(*nk*), whereas in the cross-tree-augmented and expanded modes, *Y* can contain up to *O* ((*nk*)^1.726^) candidate tripartitions. Table 1 summarizes the resulting bounds and the settings in which each method is most attractive. The memory bounds report the method-specific working memory and exclude the common *O*(|*Y*|) storage required for the precomputed weight map.

**Table 1.** Trade-offs among the intersection computation methods in ASTRAL-X. Here, *W* = ⌈*n/*64⌉, *J* is the number of candidates processed concurrently, and *H* is the maximum postorder stack frontier of the gene trees. The stated bounds report the time and memory required by the corresponding intersection-computation method, rather than the total time and memory usage of ASTRAL-X. TL, CTA, and E denote the tree-local, cross-tree-augmented, and expanded search modes, respectively.

| Method | Time (TL) | Time (CTA/E) | GPU Memory | Most suitable setting |
| --- | --- | --- | --- | --- |
| Smaller-side traversal | $O(n^2k^2)$ (balanced);<br>$O(n^3k^2)$ (worst case) | $O((nk)^{1.726}n^2k)$ (worst case) | $O(nk)$ | Approximately balanced gene trees, when many gene-tree tripartitions are repeated |
| Prefix sums | $O(n^2k^2)$ | $O((nk)^{2.726})$ | $O(nk + Jn)$ | Large taxon sets, unbalanced or unknown tree shapes |
| Postorder propagation | $O(n^2k^2)$ | $O((nk)^{2.726})$ | $O(nkW + JH)$ | Moderate-sized taxon sets with many gene trees, limited repetition among gene-tree partitions |
| Bitwise operations | $O(n^2k^2W)$ | $O((nk)^{2.726}W)$ | $O(nkW)$ | Small taxon sets |

The smaller-side method is suitable when the gene trees are approximately balanced, where it achieves the same *O*(*n*^2^*k*^2^) tree-local bound as the topology-independent methods. It can also benefit substantially from deduplication when many gene trees contain repeated tripartitions. Its performance, however, deteriorates on highly unbalanced trees, reaching *O*(*n*^3^*k*^2^) in the caterpillar case.

The prefix-sum method provides the most flexible choice for large datasets. Its *O*(*n*^2^*k*^2^) tree-local bound is independent of gene-tree topology, and in full search mode it improves upon the worst-case smaller-side bound by a factor of *n*. It also retains the compact tuple representation, making it particularly suitable when *n* is large or the balance of the gene trees is unknown.

Postorder propagation offers the same topology-independent time bounds while scoring gene-tree nodes directly, without constructing prefix arrays or a deduplicated partition table. It is therefore useful when there are many gene trees and the input contains many distinct partitions, although its bitset storage makes it less attractive for very large taxon sets.

Finally, direct bitwise intersection is the simplest and often fastest option when *n* is small and each cluster occupies only a few machine words.

In summary, ASTRAL-X favors bitwise intersection for small taxon sets, smaller-side traversal for balanced and highly repetitive gene trees, and prefix sums for large or potentially unbalanced datasets. Postorder propagation provides a complementary option for moderate-sized datasets with many gene trees and many distinct partitions.

### 2.11 Running time and memory complexity of ASTRAL-X

The overall running time and memory complexity of ASTRAL-X depend on both the search-space mode and the intersection-computation method. Tables 2 and 3 summarize these bounds for the tree-local mode and the cross-tree-augmented and expanded modes of ASTRAL-X, respectively. In tree-local mode, the number of candidate tripartitions is *O*(*nk*), which keeps the CPU memory requirement at *O*(*nk*) for all four intersection methods. The running time is then primarily determined by the cost of weight precomputation. Smaller-side traversal requires *O*(*n*^2^*k*^2^) time for balanced gene trees but can increase to *O*(*n*^3^*k*^2^) in the worst case. Prefix sums and postorder propagation both require *O*(*n*^2^*k*^2^) time, while bitwise intersection incurs an additional factor of *W* = ⌈*n/*64⌉ due to the bitset representation. The GPU VRAM requirement varies with the temporary data structures maintained by each intersection method (Table 2).

**Table 2.** Overall asymptotic complexity of ASTRAL-X in tree-local (TL) mode under the four intersection-computation methods. Here, *W* = ⌈*n/*64⌉, *J* is the number of candidate tripartitions processed concurrently during weight computation, and *H* is the maximum postorder stack frontier of the gene trees, used by the postorder-propagation method. The bounds include the complete ASTRAL-X workflow in TL mode, including input processing, construction of the tree-local DP state space, weight precomputation, storage of the precomputed weight map, and dynamic programming.

| Intersection method | Running time | CPU RAM | GPU VRAM |
| --- | --- | --- | --- |
| Smaller-side traversal | $O(n^2 k^2)$ for balanced gene trees; $O(n^3 k^2)$ in the worst case | $O(nk)$ | $O(nk)$ |
| Prefix sums | $O(n^2 k^2)$ | $O(nk)$ | $O(nk + Jn)$ |
| Postorder propagation | $O(n^2 k^2)$ | $O(nk)$ | $O(nkW + JH)$ |
| Bitwise operations | $O(n^2 k^2 W)$ | $O(nk)$ | $O(nkW)$ |

**Table 3.** Overall asymptotic complexity of ASTRAL-X in the cross-tree-augmented (CTA) and expanded (E) search modes under the four intersection-computation methods. Both modes satisfy *X* = *O*(*nk*) and |*Y*| = *O*((*nk*)^1.726^) and therefore have the same worst-case asymptotic bounds, although the expanded mode generally has a larger search space in practice. Here, *W* = ⌈*n/*64⌉, *J* is the number of candidate tripartitions processed concurrently during weight computation, and *H* is the maximum postorder stack frontier of the gene trees. The GPU bounds include the *O*(*nk* log *n*) VRAM required by the Euler-tour and sparse-RMQ structures during distance/similarity-matrix computation.

| Intersection method | Running time | CPU RAM | Peak GPU VRAM |
| --- | --- | --- | --- |
| Smaller-side traversal | $O((nk)^{1.726} n^2 k)$ (worst case) | $O(n^2 + (nk)^{1.726})$ | $O(nk \log n)$ |
| Prefix sums | $O((nk)^{2.726})$ | $O(n^2 + (nk)^{1.726})$ | $O(\max\{nk \log n, nk + Jn\})$ |
| Postorder propagation | $O((nk)^{2.726})$ | $O(n^2 + (nk)^{1.726})$ | $O(\max\{nk \log n, nkW + JH\})$ |
| Bitwise operations | $O((nk)^{2.726} W)$ | $O(n^2 + (nk)^{1.726})$ | $O(\max\{nk \log n, nkW\})$ |

The cross-tree-augmented and expanded modes have the same worst-case asymptotic complexity, although the expanded mode generally explores a larger search space in practice. The expanded mode additionally performs the consensus-guided cluster expansion described in Section 2.6.4. As established in Theorem 2.1, this phase adds only *O*(*n*) clusters, requires *O*(*nk*) additional memory, and runs in *O*(*n*^2^*k* + *nk* log(*nk*) + *n*^2^ log *n*) time in the worst case. This additional running time is asymptotically dominated by the subsequent search-space construction and weight-computation costs. Thus, it does not alter the overall complexity bounds of the expanded mode.

In both modes, *X* = *O*(*nk*) and the number of candidate tripartitions is bounded by *Y* = *O*((*nk*)^1.726^) [26]. Consequently, prefix-sum and postorder propagation each require *O*((*nk*)^2.726^) running time. Smaller-side traversal requires *O*((*nk*)^2.726^ *n*^2^*k*) time in the worst case, while bitwise intersection requires *O*((*nk*)^2.726^*W*) time, where *W* = ⌈*n/*64⌉. The larger transition space increases the CPU memory requirement to *O*(*n*^2^ +(*nk*)^1.726^), accounting for the pairwise similarity structures, candidate transitions, and precomputed weights. Peak GPU VRAM usage includes the *O*(*nk* log *n*) memory required by the Euler-tour and sparse-RMQ structures, together with the method-specific working memory used during weight precomputation, as summarized in Table 3.

## 3 Results

### 3.1 Experimental Setup

We performed extensive experiments to evaluate the scalability and accuracy of ASTRAL-X on both simulated and empirical datasets. For scalability analysis, we compared ASTRAL-X against STELARX (v2.0.0) [23], a highly scalable triplet-based summary method; ASTRAL-MP [17], a multi-parallel quartet-based implementation of ASTRAL; and ASTER [18], a faster recent implementation of the

ASTRAL framework. For accuracy analysis, we further compared ASTRAL-X with ASTRAL-MP [17], WQFM-TREE [27], and ASTER [18]. All experiments were conducted on a Linux machine equipped with an NVIDIA RTX 4090 GPU with 24 GB VRAM and an AMD EPYC 7742 processor using 32 CPU cores and 129 GB of RAM. Each analysis was given a maximum runtime of 48 hours.

### 3.2 Datasets

#### Simulated Datasets

To evaluate scalability, we simulated large phylogenomic datasets under the multispecies coalescent model using SimPhy [28]. Following the experimental design of STELAR-X [23], we first fixed the number of gene trees at *k* = 1,000 and varied the number of taxa from *n* = 1,000 to 300,000. We then fixed the number of taxa at *n* = 1,000 and varied the number of gene trees from *k* = 1,000 to 300,000. We used the same simulation parameters as in STELAR-X [23]. The details of the simulation parameters are presented in Supplementary Material (Section 4 and Table S1). To assess the robustness of scalability under different levels of gene tree discordance, we generated datasets under four model conditions, ranging from low (L1) to medium-high (L4) levels of incomplete lineage sorting (ILS). The average gene tree–gene tree (GT–GT) and gene tree–species tree (GT–ST) discordance, measured using RF rates, under the four ILS conditions is reported in Table 4. To further examine whether the scalability of ASTRAL-X is robust to gene tree estimation error, we used existing simulated datasets with 500, 1,000, and 10,000 taxa, each containing 1,000 gene trees. Finally, to evaluate accuracy, we used simulated datasets ranging from 37 to 1,000 taxa, since most existing methods cannot scale to the largest datasets used in our scalability experiments.

**Table 4.** Discordance levels under different ILS conditions analyzed in this study. We show the average gene tree-gene tree (GT–GT) and gene tree-species tree (GT–ST) discordance, measured by average RF rates, under various levels of ILS

| ILS Level | GT-GT Discordance (%) | GT-ST Discordance (%) |
| --- | --- | --- |
| ILS-L1 | 25.4 | 17.5 |
| ILS-L2 | 31.7 | 22.1 |
| ILS-L3 | 37.5 | 26.4 |
| ILS-L4 | 42.8 | 30.7 |

#### Empirical datasets

We reanalyzed three empirical phylogenomic datasets with ASTRAL-X: an angiosperm dataset comprising 9,524 taxa and 353 genes [19], an avian dataset comprising 48 taxa and 14,446 genes [29], and an extended avian dataset comprising 363 taxa and 63,430 genes [30]. We compared the trees inferred by ASTRAL-X with the corresponding published reference trees and with those inferred by STELAR-X, ASTRAL-MP, and ASTER, considering both topological agreement and computational performance in terms of running time and memory usage. Because STELAR-X requires rooted gene trees, its input trees were rooted using appropriate outgroup taxa [23]. For the 48-taxon avian dataset, *Struthio camelus* (ostrich) and *Tinamus guttatus* (white-throated tinamou) were used as outgroups. For the 363-taxon extended avian dataset, *Acanthisitta chloris*, a member of the outgroup clade Passeriformes in this dataset, was used as the outgroup.

### 3.3 Results on Scalability of Species Tree Inference

We evaluated the scalability of ASTRAL-X on simulated datasets with large numbers of taxa and gene trees, measuring average running time, CPU memory, and GPU memory usage. We compared ASTRAL-X against three scalable summary methods: STELAR-X, a highly scalable triplet-based method; ASTRAL-MP, the most scalable parallel implementation of ASTRAL; and ASTER, a recent fast implementation of the ASTRAL framework. For the first set of scalability experiments, we used true simulated gene trees with no missing taxa. We therefore ran ASTRAL-X in tree-local mode and computed the required intersection counts using the smaller-side traversal method. Later, we also evaluate the accuracy and scalability of ASTRAL-X across all three search-space modes and compare their performance with ASTRAL-MP (Section 3.3). We first fixed the number of gene trees at *k* = 1,000 and varied the number of taxa from *n* = 1,000 to 10,000 for comparison with all methods. Within our resource limits, ASTRAL-MP could be run up to 10,000 taxa, whereas ASTER could be run up to 7,500 taxa before exceeding the 48-hour limit, consistent with the scalability limits reported in previous studies [17, 18, 23].

ASTRAL-X consistently outperformed ASTRAL-MP and ASTER across all tested taxon scales, substantially reducing both running time and CPU memory usage while requiring only modestly higher GPU memory than ASTRAL-MP. ASTER was the slowest among the four methods in all completed cases, which could be due to the fact that it does not use GPU parallelization. The CPU memory usage of ASTER, however, was consistently lower than that of ASTRAL-MP, while ASTRAL-X and STELAR-X remained substantially more memory-efficient than both the other methods. For instance, on the 10,000-taxon dataset, ASTER exceeded the 48-hour time limit, while ASTRAL-MP, the previously most scalable multi-parallel implementation of ASTRAL, required 9.37 hours of runtime and 125.84 GB of CPU RAM. In contrast, ASTRAL-X completed the same analysis in only 16.45 seconds using 8.86 GB of CPU RAM, achieving an approximately 2051× speedup and a 14.20× reduction in CPU memory relative to ASTRAL-MP. The observed trends in running time and memory usage suggest that ASTRAL-MP would require substantially greater computational resources and multi-day runtimes even for moderately larger datasets, such as those with 25,000 taxa, whereas the resource requirements of ASTRAL-X increase modestly. This scaling pattern enables ASTRAL-X to analyze substantially larger datasets, as examined later in this section (Section 3.3). ASTRAL-X and STELAR-X showed comparable computational performance across the tested taxon scales. On the 10,000-taxon dataset, STELAR-X completed the analysis in 15.34 seconds using 9.09 GB of CPU RAM, compared with 16.45 seconds and 8.86 GB for ASTRAL-X. Thus, STELAR-X was slightly faster at this scale, while ASTRAL-X used slightly less CPU memory.

We next assessed scalability with respect to the number of gene trees by fixing the number of taxa at *n* = 1,000 and varying the number of gene trees from *k* = 1,000 to 50,000. In this setting, ASTRAL-MP could be executed up to 50,000 gene trees before reaching the CPU memory limit, while ASTER could be executed up to 25,000 gene trees before exceeding the runtime limit. ASTRAL-X again achieved substantial improvements over ASTRAL-MP and ASTER across all tested settings (Figure 5b). On the 50,000-gene-tree dataset, ASTRAL-X completed the analysis in 1.51 minutes using 37.89 GB of CPU RAM, whereas ASTRAL-MP required 76.18 minutes and 126.07 GB of CPU RAM. Thus, ASTRAL-X was approximately 50× faster and used 3.33× less CPU memory. Compared with STELAR-X, ASTRAL-X was faster on the datasets with up to 25,000 gene trees, although their relative CPU memory usage varied across these settings. On the largest 50,000-gene-tree dataset, however, STELAR-X was 2.43× faster while ASTRAL-X used approximately 1.35× less CPU memory. Overall, these experiments demonstrate that ASTRAL-X scales efficiently with both the number of taxa and the number of gene trees, removing the primary computational bottlenecks that limit existing ASTRAL implementations on ultra-large datasets.

**Figure 5.**
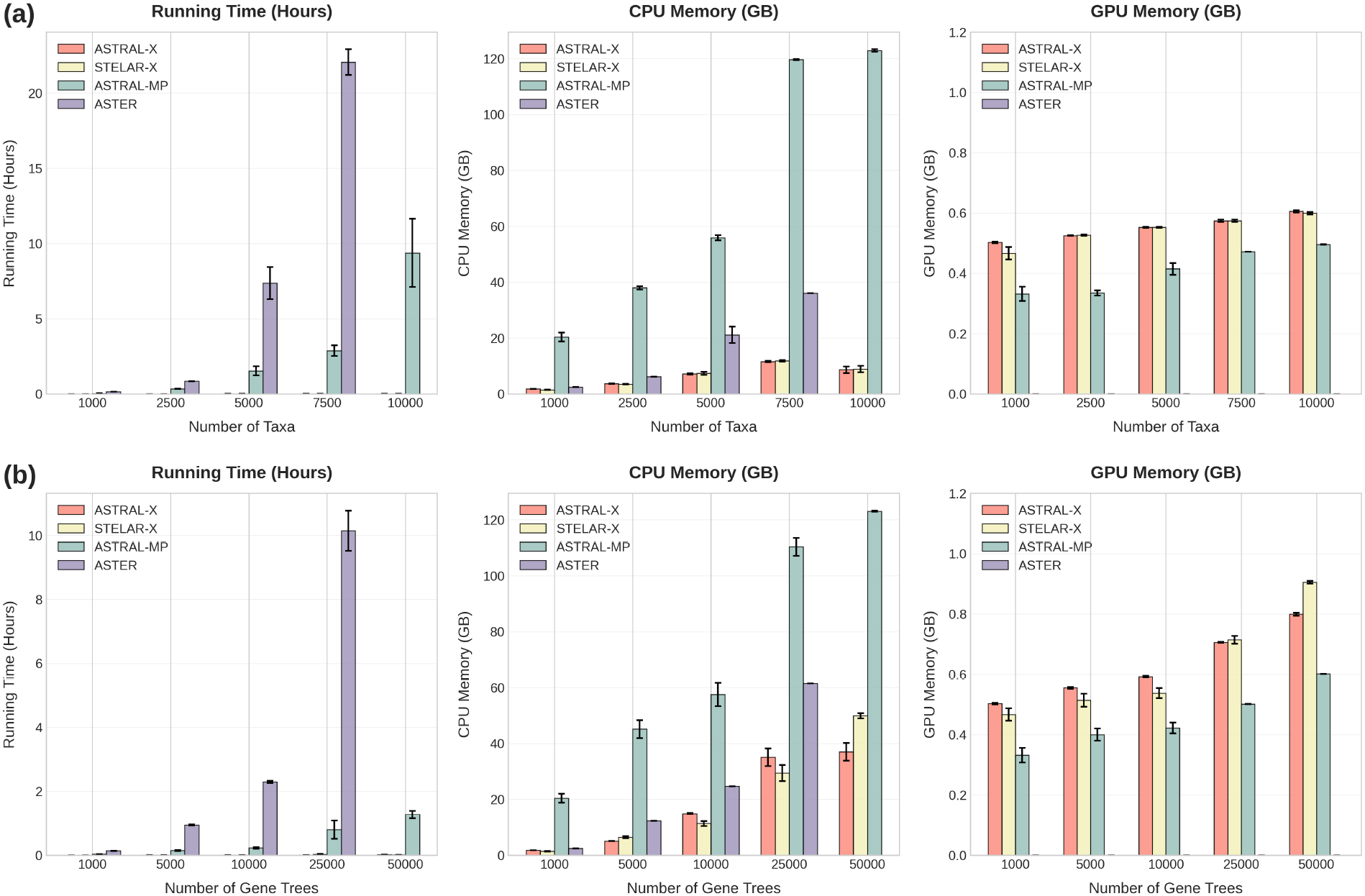
Average running time, CPU RAM, and GPU VRAM requirements of ASTRAL-X, STELAR-X, ASTRAL-MP, and ASTER over 5 replicates (except for 10,000-taxon and 50,000-gene conditions where ASTER could not be run due to its high computational demands). (a) 1,000 gene trees with 1,000–10,000 taxa, (b) 1,000 taxa and 1,000–50,000 genes. Error bars represent the standard error of the mean across replicates.

#### Scalability at larger scales of inference

We next evaluated ASTRAL-X beyond the operating range of existing ASTRAL implementations by fixing the number of gene trees at *k* = 1,000 and increasing the number of taxa to 300,000 (Figure 6a). ASTRAL-X successfully analyzed the 300,000-taxon dataset in 11.65 hours using 94.25 GB of CPU RAM and 3.87 GB of GPU VRAM. Its CPU and GPU memory requirements exhibited an approximately linear growth overall, consistent with the *O*(*nk*) memory complexity of ASTRAL-X in the tree-local search space. Notably, ASTRAL-X analyzed the 300,000-taxon dataset using only 94 GB of CPU RAM, whereas ASTRAL-MP could not scale beyond 10,000 taxa within our 128-GB memory limit. STELAR-X could be executed up to 250,000 taxa within our memory limits (128 GB). The two methods showed broadly comparable scalability, although their relative running times varied across taxon scales. On the 250,000-taxon dataset, ASTRAL-X completed the analysis in 6.44 hours using 82.95 GB of CPU RAM, compared with 6.70 hours and 112.39 GB for STELAR-X. Thus, ASTRAL-X was slightly faster while using approximately 1.35× less CPU memory at this scale. Beyond this point, ASTRAL-X continued to 300,000 taxa while remaining within the same 128-GB memory limit. GPU memory usage remained modest for both methods.

**Figure 6.**
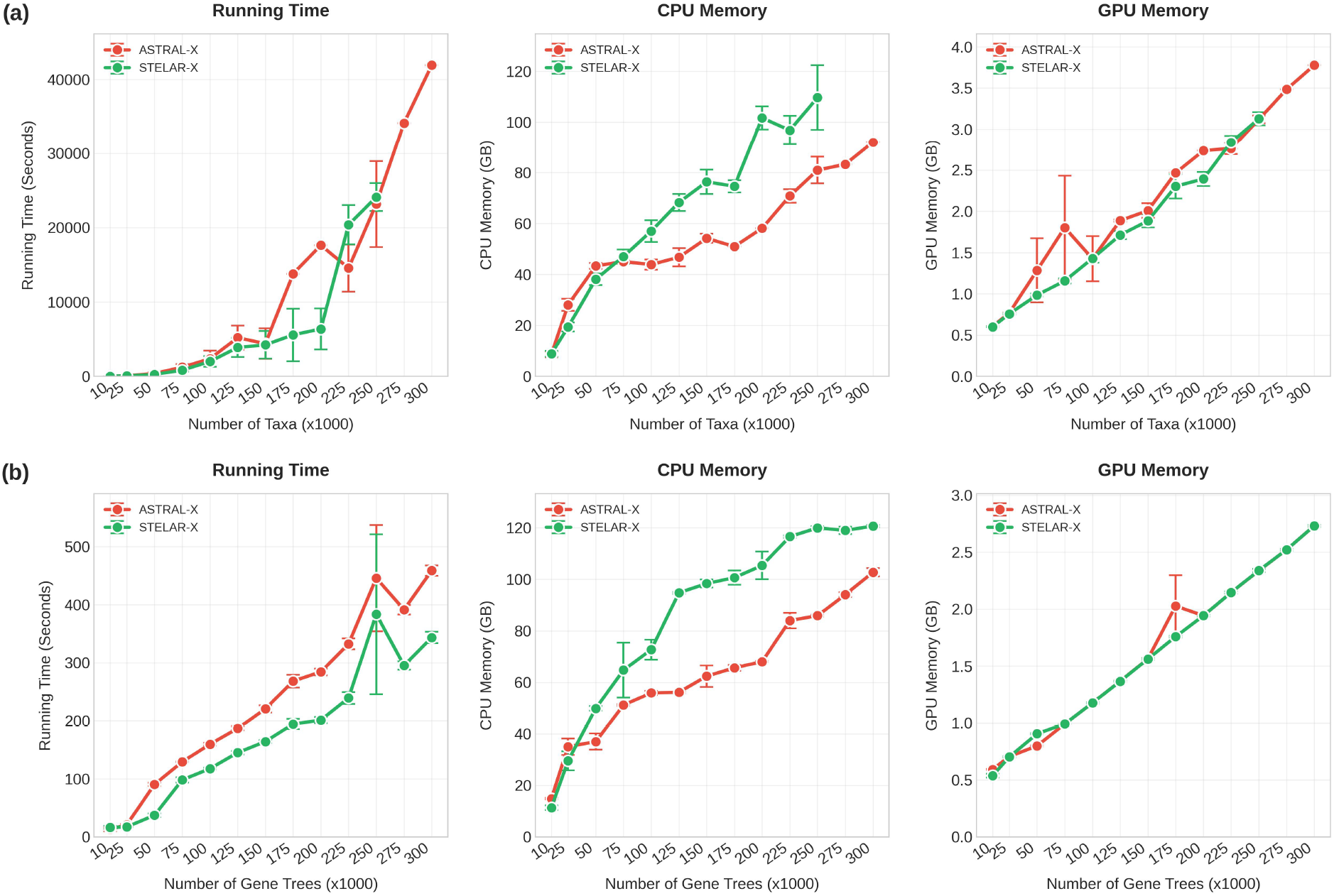
Average running time, CPU RAM, and GPU VRAM usage of ASTRAL-X and STELAR-X on large datasets over 5 replicates. (a) 10,000 to 300,000 taxa with 1,000 gene trees (STELAR-X could be run up to the 250,000 taxa dataset within our memory limit), (b) 1,000 taxa with 10,000 to 300,000 gene trees. Error bars show the standard error of the mean across replicates.

Next, we evaluated scalability with respect to the number of gene trees by fixing the number of taxa at *n* = 1,000 and increasing the number of gene trees to 300,000 (Figure 6b). ASTRAL-X analyzed the 300,000-gene-tree dataset in only 7.65 minutes using 105.23 GB of CPU RAM and 2.80 GB of GPU VRAM. In comparison, ASTRAL-MP required 76.18 minutes and 126.07 GB of CPU RAM for a dataset with only 50,000 gene trees, beyond which it exceeded our memory limit. Thus, even at a sixfold larger scale, ASTRAL-X completed the analysis in approximately one-tenth of the time while using less CPU memory. STELAR-X also scaled to 300,000 gene trees and was slightly faster than ASTRAL-X at the larger gene-tree scales, while ASTRAL-X required less CPU memory from 50,000 gene trees onward. On the 300,000-gene-tree dataset, STELAR-X completed the analysis in 5.72 minutes using 123.59 GB of CPU RAM, compared with 7.65 minutes and 105.23 GB for ASTRAL-X. GPU memory usage remained modest and nearly identical between the two methods.

#### Effect of varying levels of ILS on scalability

We next examined how the level of incomplete lineage sorting affects the scalability of ASTRAL-X. We considered four model conditions, ranging from low (ILS-L1) to medium-high (ILS-L4) ILS (see the discordance amounts in Table 4), using datasets with 1,000–40,000 taxa and 1,000 gene trees, as well as datasets with 1,000 taxa and 1,000–50,000 gene trees (Figure 7). Higher levels of ILS generally increased the running time, especially as the number of taxa grew. For example, on the 40,000-taxon datasets, the running time increased from 232.05 seconds under ILS-L2 to 848.08 seconds under ILS-L4, an approximately 3.7× increase. In comparison, memory usage remained much less sensitive to ILS: CPU RAM increased only from 24.79 GB to 28.34 GB, while GPU VRAM increased from 0.93 GB to 1.19 GB. When scaling the number of gene trees, the influence of ILS on running time was more moderate; even at 50,000 gene trees, ASTRAL-X completed all four model conditions within approximately one minute. CPU and GPU memory usage remained broadly comparable across ILS levels and continued to follow the same overall scaling trend. These results indicate that increased gene-tree discordance primarily affects computational time, while the memory efficiency of ASTRAL-X remains largely robust across different levels of ILS, supporting its asymptotically optimal memory complexity of *O*(*nk*).

**Figure 7.**
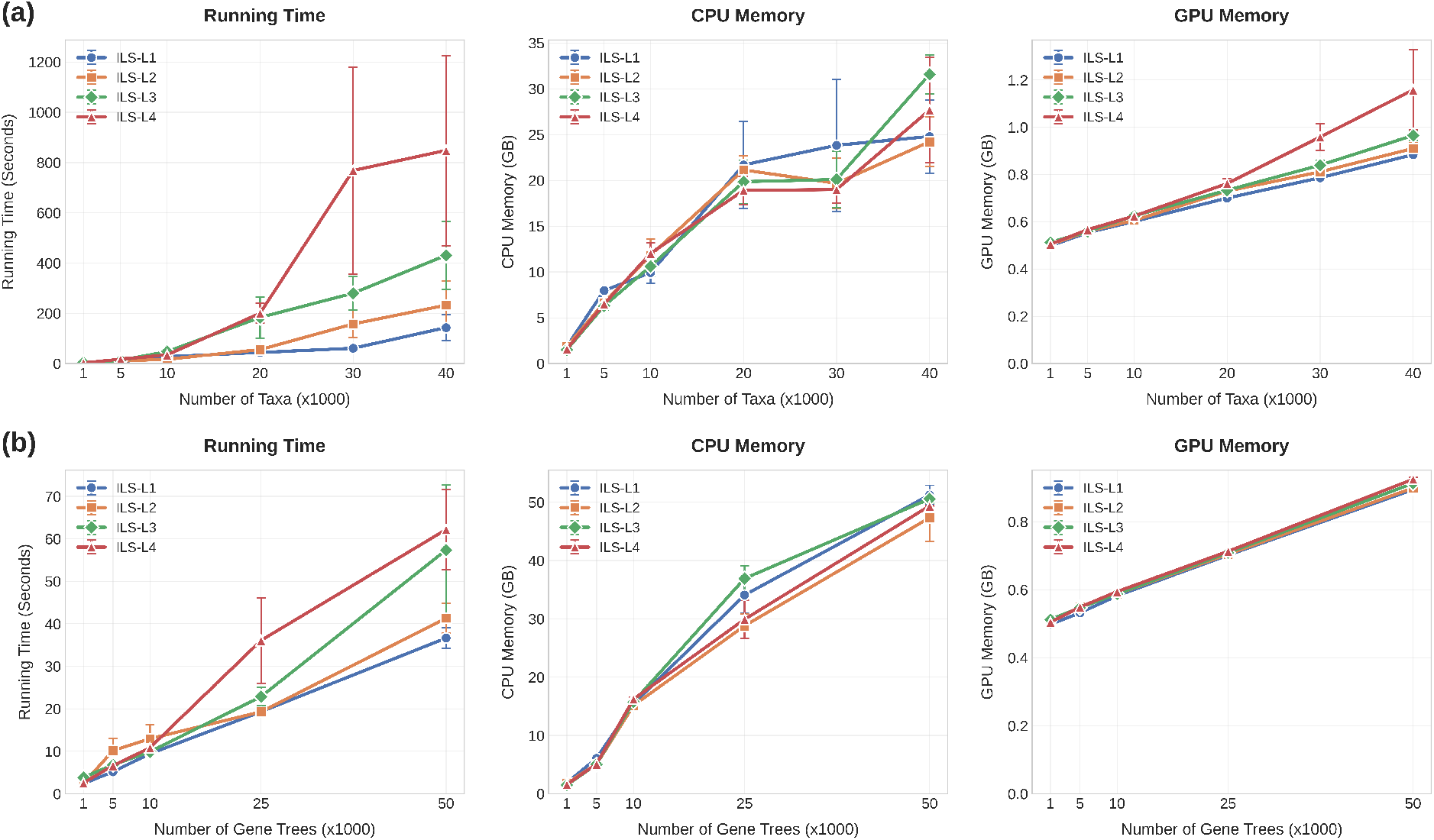
Effect of varying levels of ILS on the running time and memory usage of ASTRAL-X across 4 model conditions, using simulated datasets with (a) 1,000–40,000 taxa and 1,000 gene trees (b) 1,000 taxa and 1,000-50,000 gene trees. The error bars indicate the standard error of the mean across replicates.

#### Scalability of inference for estimated gene trees

We next examined how gene tree estimation error affects the scalability of ASTRAL-X using three publicly available simulated datasets. The first two datasets contain 500 and 1,000 taxa, respectively, with 1,000 gene trees each. They were originally studied in ASTRAL-II [13] and later reanalyzed in the WQFM-TREE study [27] and the STELAR-X study [23]. Their species trees were generated under a Yule process with a maximum tree length of 2M and a speciation rate of 1 × 10^−6^. We also included the larger simulated dataset from the ASTRAL-MP study [17], comprising 10,000 taxa and 1,000 gene trees generated with SimPhy [28] under the multispecies coalescent model. For inference on these datasets, we used the cross-tree-augmented search mode of ASTRAL-X and the smaller-side-traversal method for computing intersection counts. For each dataset, we compared ASTRAL-X, STELAR-X, and ASTRAL-MP using both the true gene trees and their corresponding estimated gene trees. Since STELAR-X operates on rooted input gene trees, for the STELAR-X analyses the estimated gene trees were rooted using the designated outgroup taxon for each dataset. Results were averaged over 10 replicates (Table 5).

**Table 5.**
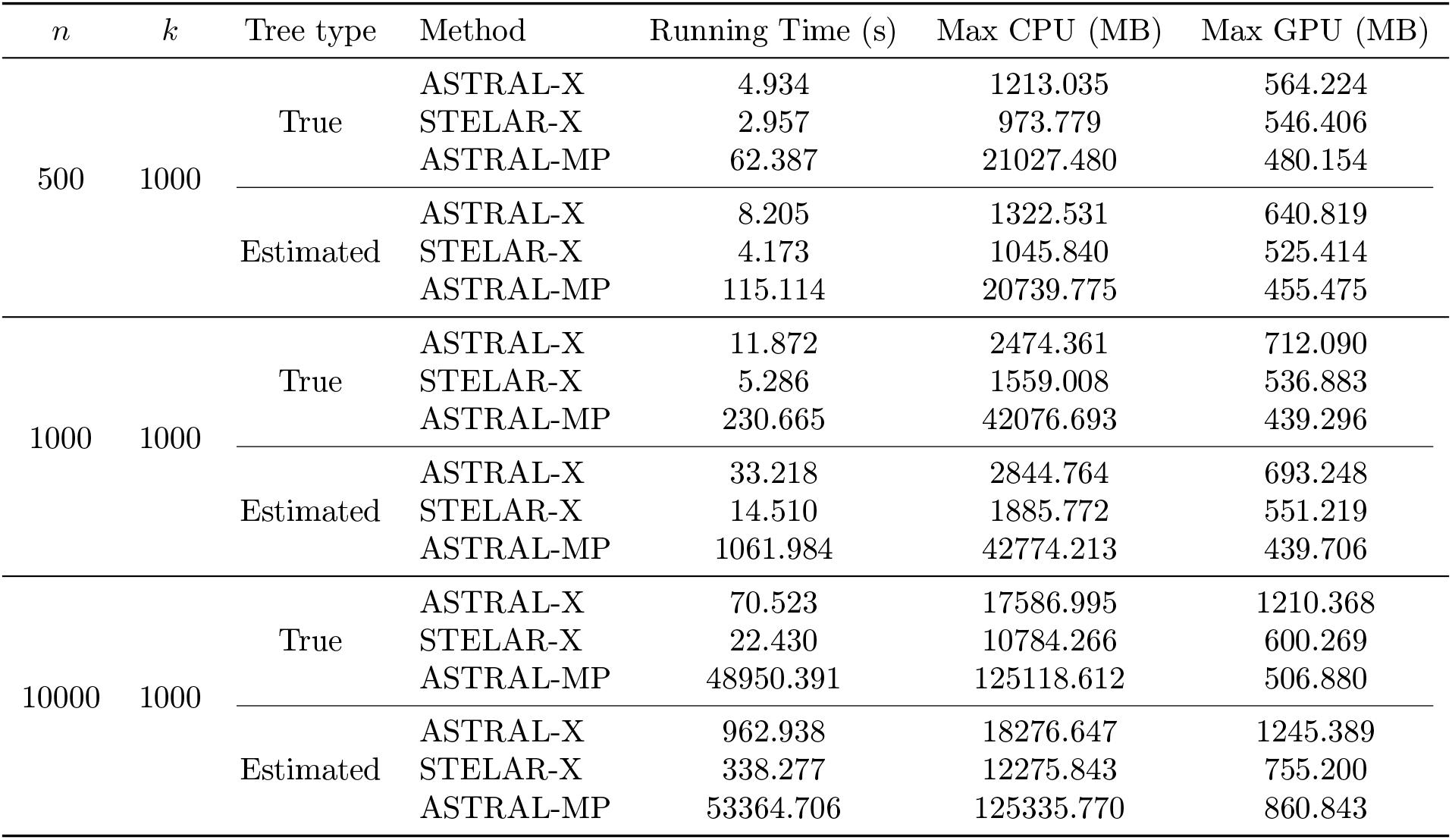
Comparison of running time and memory usage among ASTRAL-X, STELAR-X, and ASTRAL-MP for true vs. estimated gene trees in simulated datasets, averaged over 10 replicates.

| $n$ | $k$ | Tree type | Method | Running Time (s) | Max CPU (MB) | Max GPU (MB) |
| --- | --- | --- | --- | --- | --- | --- |
| 500 | 1000 | True | ASTRAL-X | 4.934 | 1213.035 | 564.224 |
|  |  |  | STELAR-X | 2.957 | 973.779 | 546.406 |
|  |  |  | ASTRAL-MP | 62.387 | 21027.480 | 480.154 |
|  |  | Estimated | ASTRAL-X | 8.205 | 1322.531 | 640.819 |
|  |  |  | STELAR-X | 4.173 | 1045.840 | 525.414 |
|  |  |  | ASTRAL-MP | 115.114 | 20739.775 | 455.475 |
| 1000 | 1000 | True | ASTRAL-X | 11.872 | 2474.361 | 712.090 |
|  |  |  | STELAR-X | 5.286 | 1559.008 | 536.883 |
|  |  |  | ASTRAL-MP | 230.665 | 42076.693 | 439.296 |
|  |  | Estimated | ASTRAL-X | 33.218 | 2844.764 | 693.248 |
|  |  |  | STELAR-X | 14.510 | 1885.772 | 551.219 |
|  |  |  | ASTRAL-MP | 1061.984 | 42774.213 | 439.706 |
| 10000 | 1000 | True | ASTRAL-X | 70.523 | 17586.995 | 1210.368 |
|  |  |  | STELAR-X | 22.430 | 10784.266 | 600.269 |
|  |  |  | ASTRAL-MP | 48950.391 | 125118.612 | 506.880 |
|  |  | Estimated | ASTRAL-X | 962.938 | 18276.647 | 1245.389 |
|  |  |  | STELAR-X | 338.277 | 12275.843 | 755.200 |
|  |  |  | ASTRAL-MP | 53364.706 | 125335.770 | 860.843 |

Gene tree estimation error generally increased the running time and CPU memory requirements of all methods, with the effect becoming more evident at larger taxon scales. Both ASTRAL-X and STELAR-X remained substantially more efficient than ASTRAL-MP across all three datasets. On the estimated 10,000-taxon dataset, ASTRAL-X completed the analysis in 962.94 seconds using 18.28 GB of CPU RAM, whereas STELAR-X required 338.28 seconds and 12.28 GB. In comparison, ASTRAL-MP required approximately 14.8 hours and 125.34 GB of CPU RAM. Thus, ASTRAL-X remained approximately 55.4× faster and used 6.86× less CPU memory than ASTRAL-MP. STELAR-X was faster and more CPU-memory-efficient than ASTRAL-X on these datasets, requiring approximately 2.85× less running time and 1.49× less CPU memory on the 10,000-taxon estimated-gene-tree dataset. GPU memory usage remained modest for all methods across these experiments.

#### Effect of Search-Space Expansion on Accuracy and Scalability

Since ASTRAL-X provides three search-space modes, we now examine how the choice of mode affects its accuracy, running time, and memory usage. Table 6 compares the tree-local, cross-tree-augmented, and expanded modes on three simulated datasets using both true and estimated gene trees, with results averaged over 10 replicates. We also include the results for ASTRAL-MP as a baseline for comparison. We further examine how the computational requirements of these modes scale to much larger simulated datasets by varying the number of taxa and gene trees using true gene trees (Figure 8).

**Table 6.**
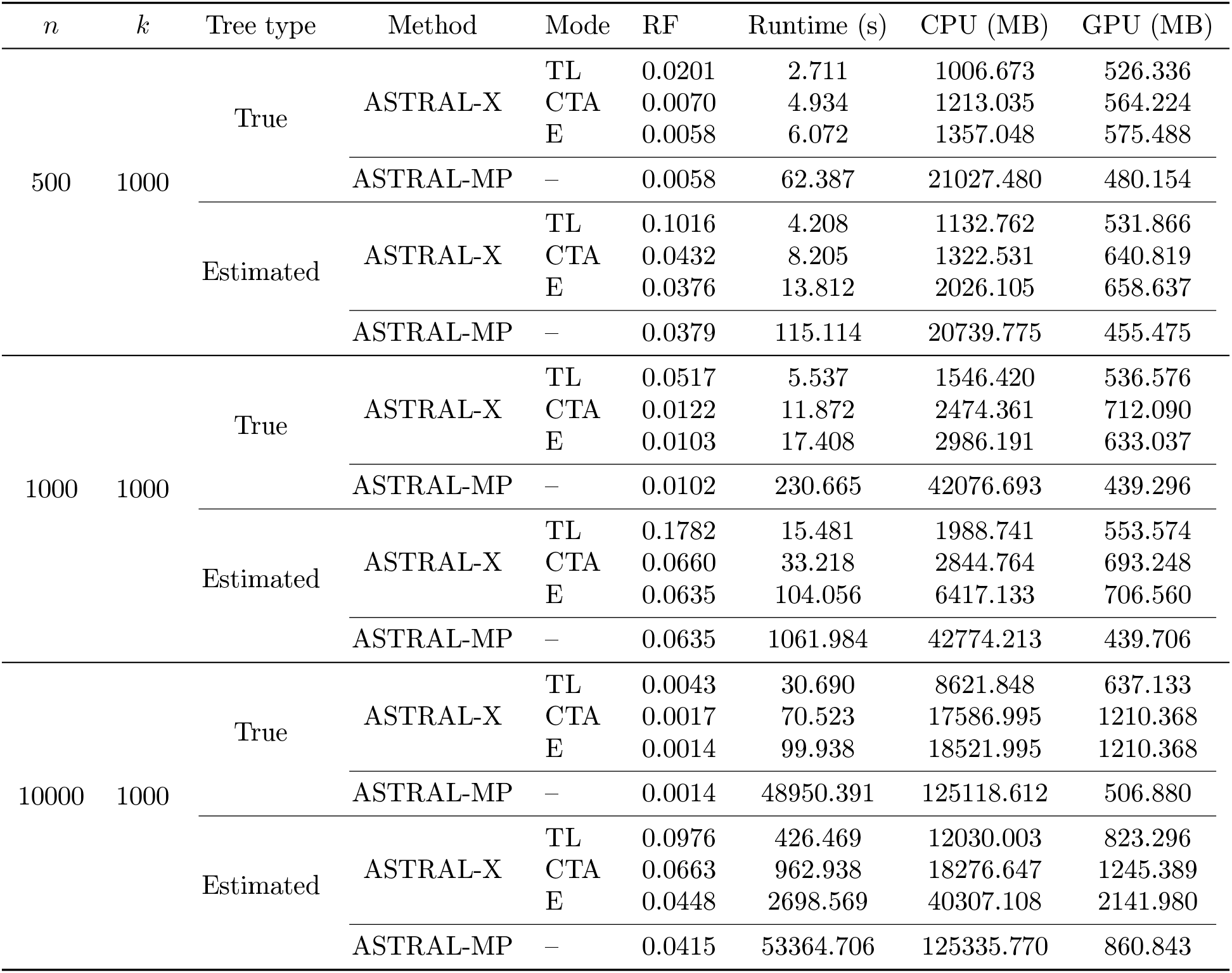
Comparison of accuracy, running time, and memory usage of ASTRAL-X under different search-space modes for true and estimated gene trees in simulated datasets. Results for ASTRAL-MP are also included as a baseline for comparison. Results are averaged over 10 replicates. TL, CTA, and E denote the tree-local, cross-tree-augmented, and expanded search-space modes of ASTRAL-X, respectively.

**Figure 8.**
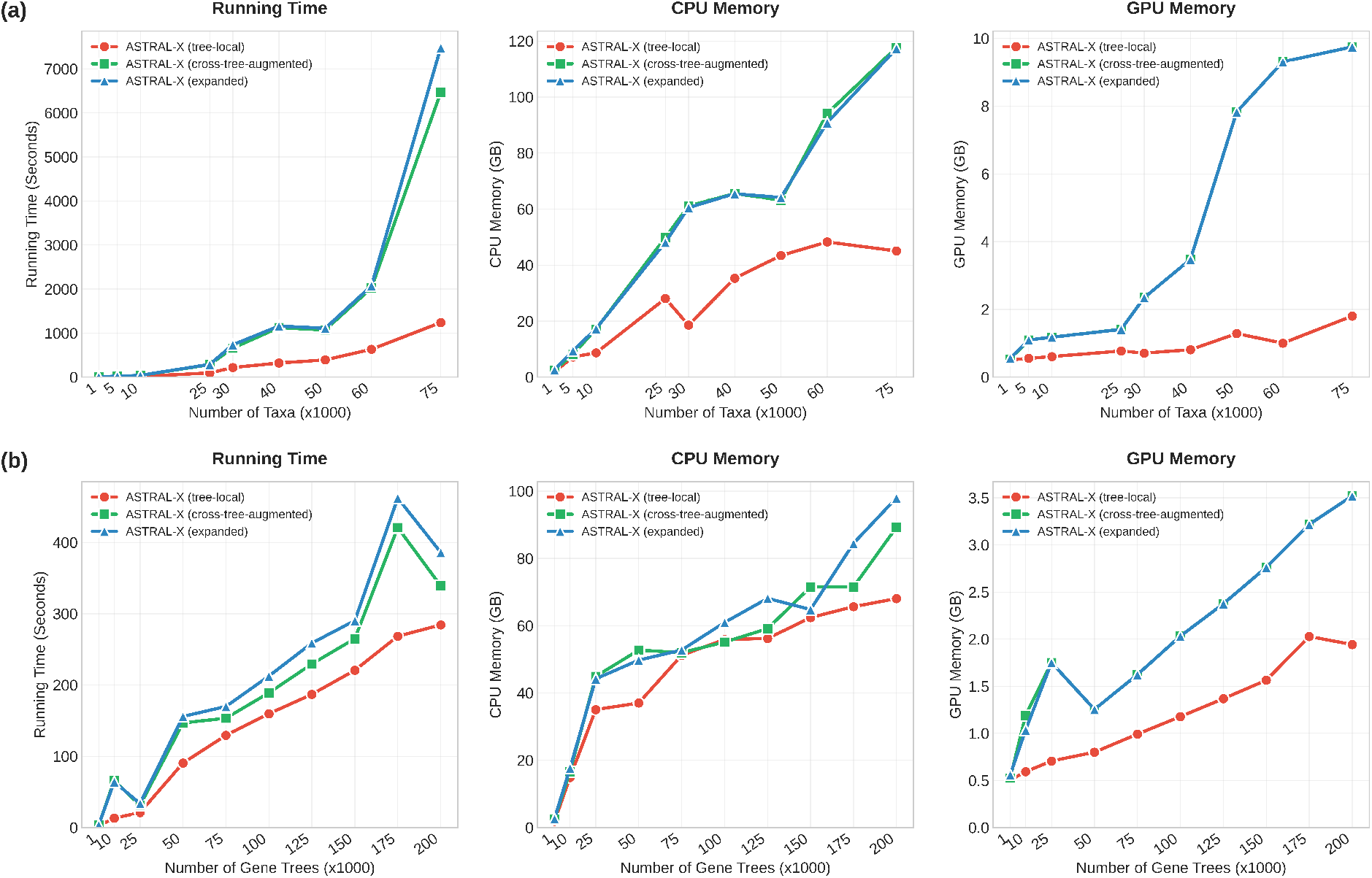
Comparison of the running time, CPU memory, and GPU memory usage of ASTRAL-X across its three search-space modes (tree-local, cross-tree-augmented, and expanded) on simulated datasets with (a) 1,000–75,000 taxa and 1,000 gene trees, and (b) 1,000 taxa and 1,000–200,000 gene trees. Results are averaged over 5 replicates.

As expected, enlarging the search space generally improves inference accuracy, while requiring additional computation (Table 6). This improvement is particularly evident for estimated gene trees, where the clusters observed in individual gene trees may provide an inadequate search space, especially when the number of gene trees is small or when the gene trees contain substantial missing taxa. On the 10,000-taxon, 1000-gene dataset with estimated gene trees, for example, the RF error rate decreased from 0.0976 in tree-local mode to 0.0663 in cross-tree-augmented mode and further to 0.0448 in expanded mode. The cross-tree-augmented and expanded modes were often close in accuracy, although the expanded mode generally achieved the lowest RF error. These gains come with the expected increase in running time and memory usage as broader sets of candidate clusters and DP transitions are considered. Nevertheless, all three ASTRAL-X modes remained substantially faster and more CPU-memory efficient than ASTRAL-MP. On the 10,000-taxon dataset with true gene trees, expanded-mode ASTRAL-X achieved the same RF error as ASTRAL-MP (0.0014), while running in about 100 seconds compared with 48,950 seconds for ASTRAL-MP, a nearly 490× speedup. It also used about 18.5 GB of CPU memory compared with 125.1 GB for ASTRAL-MP. With estimated gene trees, expanded-mode ASTRAL-X remained about 20× faster and used roughly one-third of the CPU memory of ASTRAL-MP, while obtaining a similar RF rate of 0.0448 compared to 0.0415. Similarly, on the 500- and 1000-taxon datasets with estimated gene trees, expanded-mode ASTRAL-X matched or slightly improved upon the accuracy of ASTRAL-MP, while being about 8–10× faster and using 7–10× less CPU memory.

Nevertheless, the broader search spaces of ASTRAL-X remain practical at very large scales. Notably, ASTRAL-X could analyze datasets with up to 75,000 taxa in expanded mode within our 128-GB CPU memory limit, completing the analysis in approximately 2.1 hours (Figure 8a). Scalability with respect to the number of gene trees was more favorable. All three modes could analyze datasets containing up to 200,000 gene trees within our resource limits (Figure 8b). The expanded mode completed this dataset in only 386 seconds using about 100 GB of CPU memory and 3.6 GB of GPU memory.

These results present the trade-off among the three search-space modes. The expanded mode is generally recommended when time and memory resources permit, particularly for estimated gene trees or datasets with relatively few gene trees or a high prevalence of missing taxa. The cross-tree-augmented mode often achieves similar accuracy with a lower computational cost. In contrast, the tree-local mode is well suited to datasets where the input gene trees already provide sufficient coverage of the search space, offering substantially lower running time and memory usage while often providing comparable accuracy.

### 3.4 Results on Accuracy of Inference

We next examined whether ASTRAL-X preserves the accuracy of existing ASTRAL-style methods while substantially improving scalability. We compared ASTRAL-X with ASTRAL-MP and ASTER, two scalable implementations of the ASTRAL framework, and with wQFM-TREE, a quartet-amalgamation-based method. We first evaluated the methods on a biologically based simulated dataset with 37 taxa under varying levels of ILS and gene tree estimation error, and then on simulated datasets with 100, 200, and 500 taxa and 1,000 gene trees (Figure 9a–c). For these datasets, we used the cross-tree-augmented mode of ASTRAL-X. Across all model conditions and dataset sizes, ASTRAL-X closely matched the accuracy of ASTRAL-MP and ASTER, with no statistically significant differences in RF error rates (*p >* 0.05). wQFM-TREE achieved the lowest RF rates in most cases, consistent with previous observations that wQFM-like quartet amalgamation methods can be highly accurate [27, 31], although ASTRAL-X remained competitive under all model conditions.

**Figure 9.**
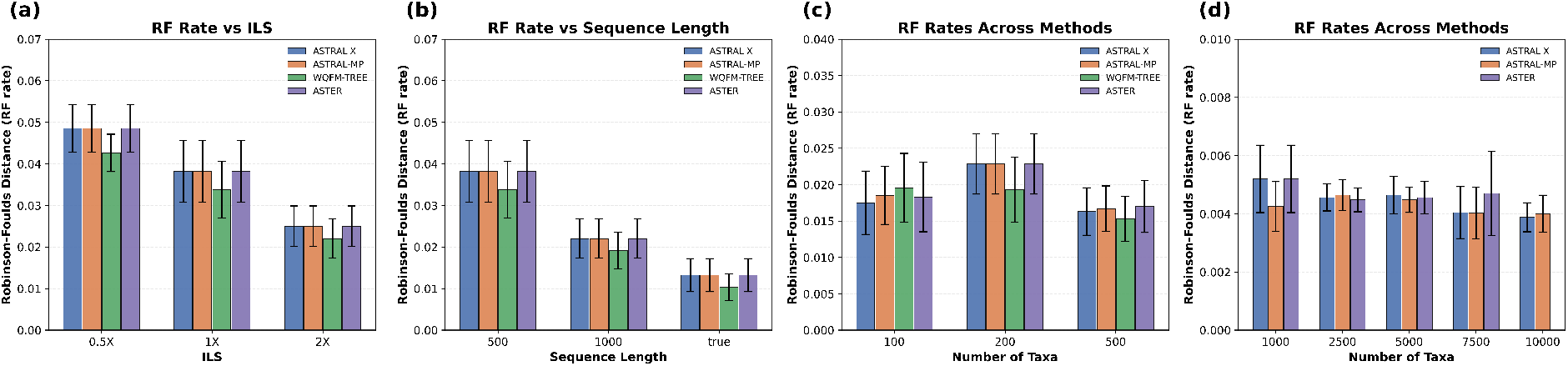
(a)-(c) Average RF rates of ASTRAL-X, ASTRAL-MP, WQFM-TREE, and ASTER over 10 replicates (a) varying ILS (37 taxa) (b) varying sequence lengths (37 taxa) (c) 100–500 taxa (d) RF rates on 1,000–10,000 taxa over 5 replicates except for ASTER on 10,000 taxa. Error bars correspond to the standard error of the mean across replicates.

To evaluate accuracy at larger scales, we further compared ASTRAL-X with ASTRAL-MP and ASTER on our simulated datasets containing 1,000 to 10,000 taxa and 1,000 gene trees. Since these datasets involved true gene trees, we used the tree-local search space of ASTRAL-X. wQFM-TREE was excluded from these experiments because it could not complete these analyses within our resource limits, and ASTER was excluded from the 10,000-taxon case due to the runtime limit. In these large-scale experiments, ASTRAL-X maintained accuracy comparable to ASTRAL-MP and ASTER, with only minor variations across dataset sizes (Figure 9d). Since these experiments used true gene trees rather than estimated gene trees, the RF rates were low for all methods. ASTRAL-MP obtained slightly lower RF rates in the 1,000- and 5,000-taxon cases, whereas ASTRAL-X outperformed ASTRAL-MP in the 2,500-, 7,500-, and 10,000-taxon cases. ASTER remained similar to the other two methods while exhibiting modestly higher RF rates in the 7,500-taxon case.

### 3.5 Results on Biological Datasets

#### 3.5.1 Analysis of the angiosperms dataset

We reanalyzed the angiosperm dataset of Zuntini et al. [19], which contains 9,524 taxa and 353 gene trees. ASTRAL-X reconstructed the angiosperm phylogeny in only 16 minutes, using 34.6 GB of CPU RAM and 3.0 GB of GPU VRAM. For comparison, the ASTRAL-MP run log provided in the GitHub repository of the reference study reports a runtime of approximately 48 hours, making the analysis about 180× slower than ASTRAL-X on the same dataset. We also attempted to run ASTRAL-MP under our experimental setup, but the analysis exceeded our available CPU memory limit (129 GB). Importantly, the species tree inferred by ASTRAL-X (Figure 10) was broadly consistent with the phylogeny reported by Zuntini et al. [19] and with several relationships supported by previous angiosperm studies.

**Figure 10.**
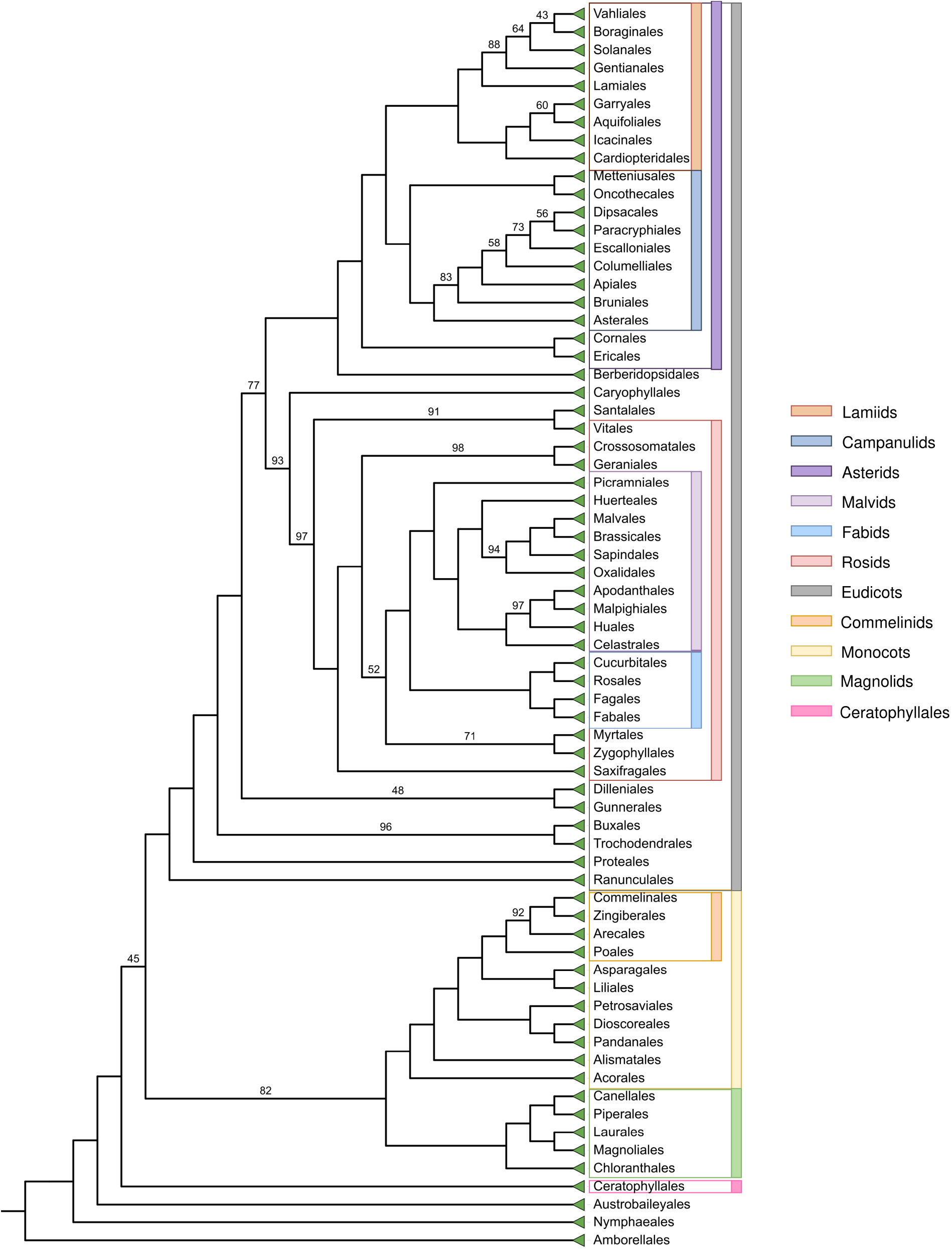
ASTRAL-X inferred tree on the angiosperms dataset. Branch supports (BS) based on quartet-based local posterior probabilities [32] are shown on branches. All BS values are 100% except where noted.

For this dataset, ASTRAL-X was run in expanded search-space mode because it contained only 353 genes and the gene trees had missing taxa. The required intersection counts were computed using the postorder-propagation method. The tree inferred by ASTRAL-X achieved a quartet score of 118,945,675,842,574,646, which is within 0.001% of the quartet score of the published reference tree (118,946,562,764,431,780).

To further assess the quality of the inferred tree, we examined whether each recognized angiosperm order forms a monophyletic clade in the ASTRAL-X species tree. Among the 74 recognized orders represented in the dataset, 71 (95.9%) were recovered as monophyletic without any post-processing. The remaining three orders (Arecales, Escalloniales, and Malpighiales) each differed from monophyly because only a very small number of taxa were separated from an otherwise intact clade, affecting a total of only seven taxa across the entire dataset (Table 7).

**Table 7.** Orders that were not initially recovered as monophyletic in the angiosperm species tree inferred by ASTRAL-X due to a small number of misplaced taxa.

| Order | Total taxa | Misplaced taxa |
| --- | --- | --- |
| Arecales | 183 | 4 |
| Escalloniales | 5 | 2 |
| Malpighiales | 498 | 1 |

We next present several observations on the angiosperm phylogeny inferred by ASTRAL-X in comparison with previous studies.

##### ASTRAL-X recovered stable relationships among Asterids

The relationships among Asterids inferred by ASTRAL-X were largely consistent with previous studies. Ericales and Cornales formed a clade as sister to all other Asterids, matching the placement reported by the 1KP study [33]. The remaining orders were divided into the two major clades Lamiids and Campanulids, a split that has long been supported by floral ontogenetic evidence [19, 34]. Within Campanulids, the phylogeny inferred by ASTRAL-X agreed with Zuntini et al. [19], placing Metteniusales and Oncothecales together as a subclade, sister to the remaining Campanulids. Within Lamiids, however, ASTRAL-X placed Cardiopteridales as sister to the clade containing Garryales, Aquifoliales, and Icacinales, whereas Zuntini et al. [19] placed Cardiopteridales as sister to all other Lamiids.

##### Confirmatory placement of Amborellales

Consistent with Zuntini et al. [19], ASTRAL-X placed Amborellales as sister to all other angiosperms. The placement of Ceratophyllales also matched the tree reported by Zuntini et al. [19], although earlier studies proposed conflicting positions for this clade [33,35].

##### ASTRAL-X recovered the long-supported Fabids clade

Within Rosids, ASTRAL-X recovered the two major clades Malvids and Fabids. The Fabids clade inferred by ASTRAL-X, comprising Cucurbitales, Fabales, Fagales, and Rosales, agrees with the tree reported by Zuntini et al. [19]. This grouping has also long been supported by evidence associated with symbiotic nitrogen fixation [19, 36].

##### Placement of Oxalidales in a different subclade

Within Malvids, ASTRAL-X placed Oxalidales with Malvales, Brassicales, and Sapindales, in contrast to the primarily plastid-based reconstruction of APG IV [37], where Oxalidales was placed in the Celastrales–Oxalidales–Malpighiales (COM) clade. However, the placement inferred by ASTRAL-X is consistent with Zuntini et al. [19].

##### The tree inferred by ASTRAL-X supported evidence of whole-genome duplication and early hybridization

Similar to Zuntini et al. [19], ASTRAL-X placed Dilleniales and Gunnerales as sister lineages, although this relationship received low support. This weakly supported placement may reflect the influence of ancient whole-genome duplication events [33, 38]. In addition, the low support for the branch separating Eudicots from Magnoliids, Monocots, and Ceratophyllales may be consistent with early hybridization events, as proposed in previous studies [19, 39].

#### 3.5.2 Analysis of the avian dataset

We also analyzed the avian dataset containing 48 taxa and 14,446 gene trees [29]. The species tree inferred by ASTRAL-X was topologically identical to the tree inferred by ASTRAL-MP and ASTER and was largely consistent with the published reference tree and STELAR-X re-analysis (Figure 11). In particular, ASTRAL-X recovered several major avian clades, including Australaves, Afroaves, Core Waterbirds, Columbimorphae, and Caprimulgimorphae, in agreement with the reference study [29]. However, none of the methods fully recovered the Cursores and Otidimorphae clades.

**Figure 11.**
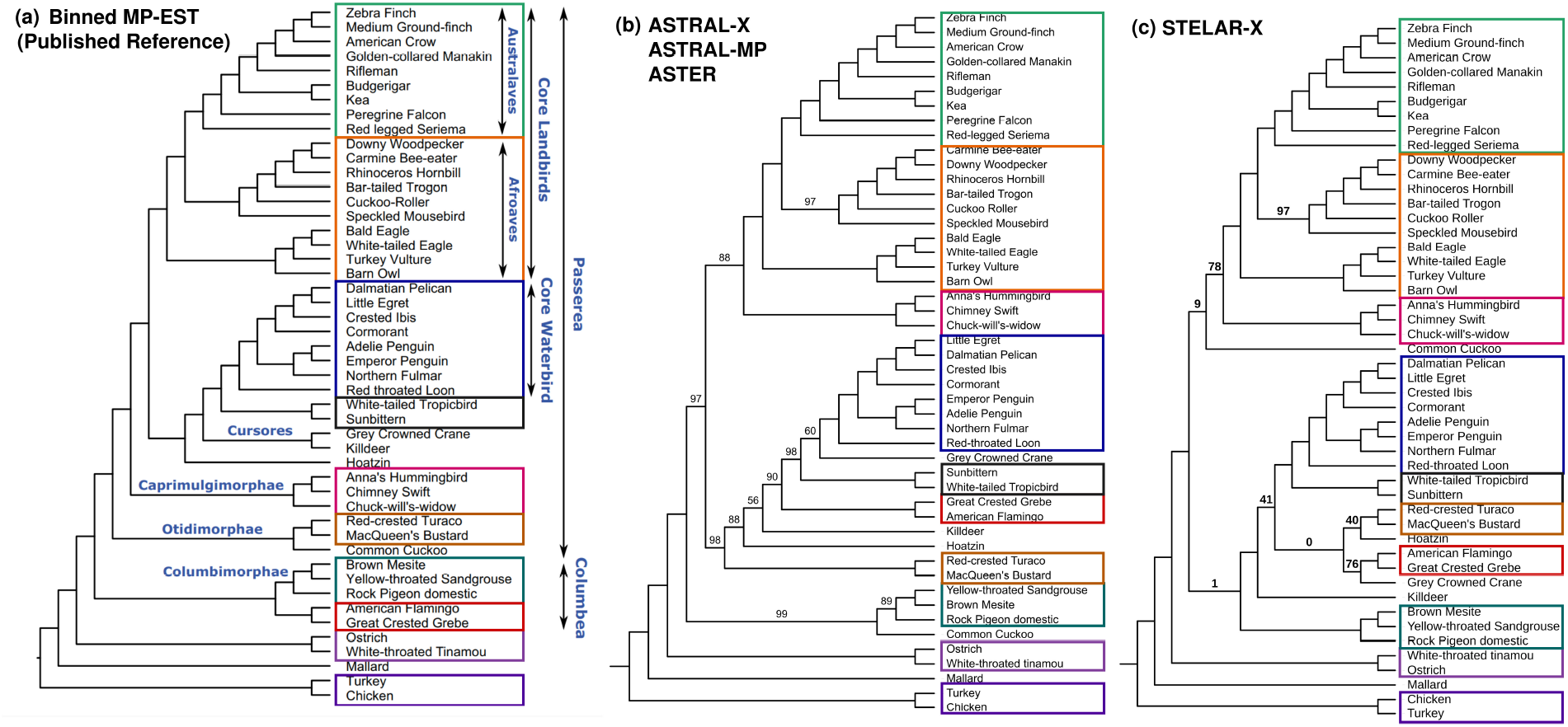
(a)-(c) Species trees inferred by Binned MP-EST (published reference), ASTRAL-X, ASTRAL-MP, ASTER, and STELAR-X on the avian dataset comprising 48 taxa and 14,446 gene trees. Identical clades are indicated with colors. The trees inferred by ASTRAL-X, ASTRAL-MP, and ASTER were topologically identical. Branch supports (BS) based on quartet-based local posterior probabilities [32] are shown on branches. All BS values are 100% except where noted.

The placement of Common Cuckoo was particularly conflicting among the inferred trees. In the published reference tree, the Common Cuckoo was placed within Otidimorphae, whereas ASTRAL-X, ASTRAL-MP, and ASTER placed it as sister to Columbimorphae with high branch support (99%). In contrast, STELAR-X placed the Common Cuckoo as sister to the clade comprising Core Waterbirds and Caprimulgimorphae. This alternative placement, however, received only 9% branch support, indicating uncertainty in this region of the tree inferred by STELAR-X. The trees inferred by ASTRAL-X, ASTRAL-MP, and ASTER each achieved a quartet score of 2,463,985,656.

Table 8 compares the running time and memory usage of ASTRAL-X, STELAR-X, ASTRAL-MP, and ASTER on this dataset. In this dataset, ASTRAL-X was run in cross-tree-augmented mode, and the intersection counts were computed using bitwise operations because the number of taxa was small. ASTRAL-X completed the analysis in only 46 seconds, achieving an approximately 7.7× speedup over ASTRAL-MP. STELAR-X completed the analysis faster, within 12 seconds, and used approximately 1.6 GB of CPU RAM, compared with 2.4 GB used by ASTRAL-X.

**Table 8.** Comparison of running time and memory usage among ASTRAL-X, STELAR-X, ASTRAL-MP, and ASTER on avian and extended avian datasets.

| Dataset | $n$ | $k$ | Method | Running Time (s) | Max CPU (MB) | Max GPU (MB) |
| --- | --- | --- | --- | --- | --- | --- |
| avian-48 | 48 | 14446 | ASTRAL-X | 46.319 | 2378.352 | 580.608 |
|  |  |  | STELAR-X | 11.361 | 1568.988 | 547.840 |
|  |  |  | ASTRAL-MP | 357.537 | 3156.684 | 176.248 |
|  |  |  | ASTER | 143.313 | 1659.660 | 0.000 |
| avian-363 | 363 | 63430 | ASTRAL-X | 432.975 | 44196.742 | 936.960 |
|  |  |  | STELAR-X | 332.775 | 28335.602 | 825.344 |
|  |  |  | ASTRAL-MP | 42007.093 | 100356.637 | 835.584 |
|  |  |  | ASTER | 41463.458 | 53667.383 | 0.000 |

#### 3.5.3 Analysis of the extended avian dataset

To further evaluate the performance of ASTRAL-X on empirical datasets with a large number of gene trees, we reanalyzed the extended avian dataset of Stiller et al. [30], comprising 363 taxa and 63,430 genes. The tree inferred by ASTRAL-X was highly consistent with the published reference tree and the re-analysis by STELAR-X and it was topologically identical to the trees inferred by ASTRAL-MP and ASTER (Figure 12). The tree reported by the STELAR-X re-analysis is presented in the Supplementary material (Figure S1). ASTRAL-X completed this analysis in only 7.2 minutes, running approximately 97× faster than ASTRAL-MP and 96× faster than ASTER (Table 8). STELAR-X completed the analysis in approximately 5.5 minutes, while also using less CPU memory. For this experiment, we first completed the incomplete gene trees and then ran ASTRAL-X in tree-local mode on the resulting completed trees, enabling this highly efficient performance. To evaluate the effect of a broader search space, we also ran ASTRAL-X in cross-tree-augmented mode. In this case, the analysis completed in 3.55 hours, remaining 3.28× faster than ASTRAL-MP. Notably, the species tree inferred in the cross-tree-augmented search space was identical to that obtained using only tree-local transitions.

**Figure 12.**
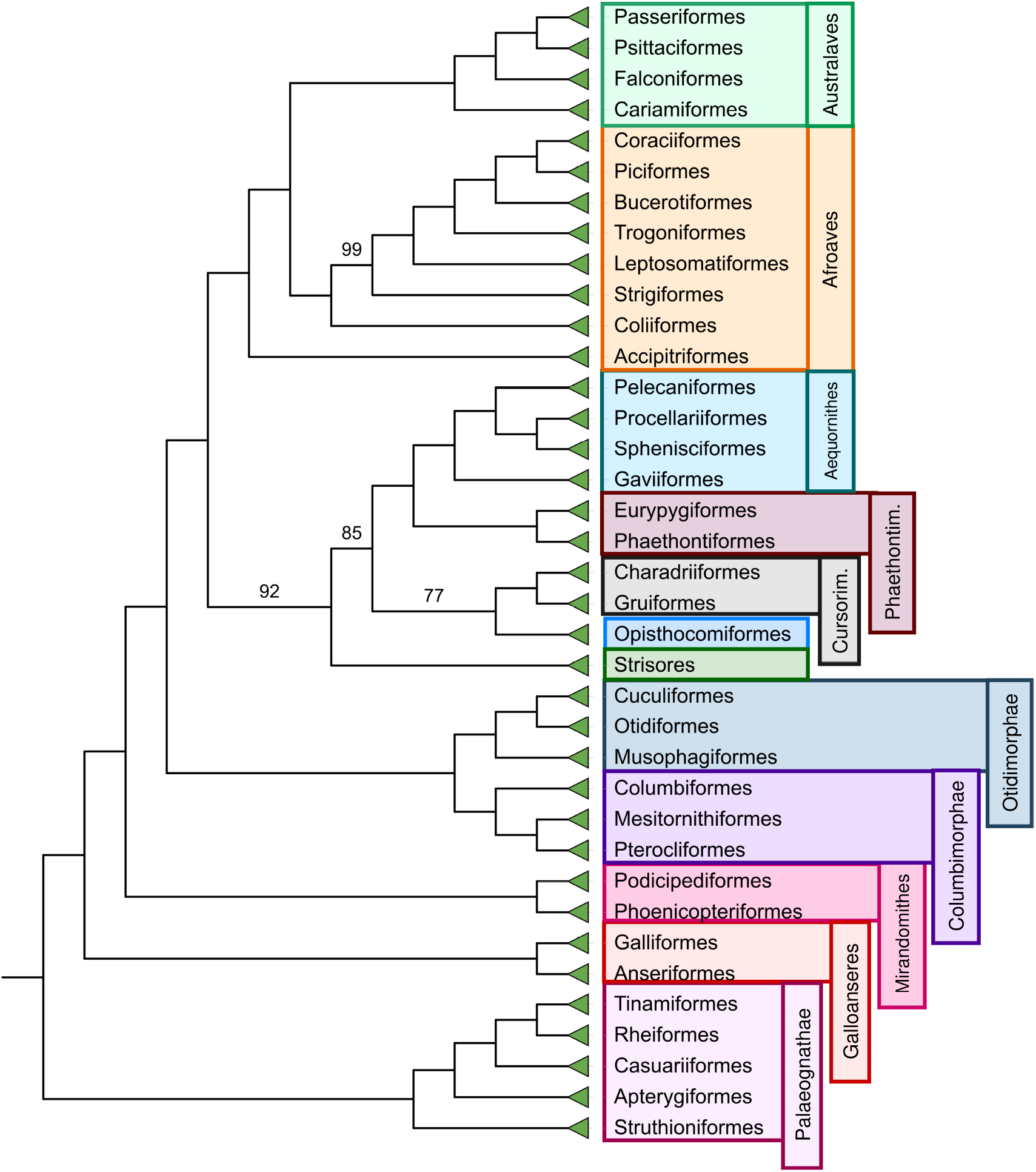
Species tree inferred by ASTRAL-X, ASTRAL-MP, and ASTER on the extended avian dataset with 363 taxa and 63,430 genes (intergenic loci). The trees inferred by the three methods were topologically identical. Branch supports (BS) based on quartet-based local posterior probabilities [32] are shown on branches. All BS values are 100% except where noted.

The tree inferred by ASTRAL-X reconstructed 11 of the 12 major avian clades, with the only exception being the placement of Accipitriformes (Figure 12). The inferred inter-clade relationships were also highly consistent with both the original study [30] and the recent reanalysis by STELAR-X [23]. Remarkably, the inferred tree achieved a quartet score of 61,832,824,956,556, exceeding the score of 61,828,805,831,222 reported for the published reference tree.

##### ASTRAL-X recovered major phylogenetic relationships

ASTRAL-X correctly reconstructed all 218 families and all 37 orders reported in the extended avian dataset [30]. It also recovered the major clades Columbimorphae, Mirandornithes, Cursorimorphae, Aequornithes, Phaethontimorphae, Australaves, and Afroaves. The only exception involved Accipitriformes, which ASTRAL-X placed as sister to all other Telluraves. This differs from previous reconstructions, where Accipitriformes was placed within Afroaves [23, 30].

##### Mirandornithes was placed as sister to all other Neoaves

Consistent with Stiller et al. [30], ASTRAL-X placed Mirandornithes as sister to all other Neoaves. This placement differs from both the STELAR-X reanalysis [23] and an earlier avian phylogenomic study [29], where Mirandornithes was placed as sister to Columbimorphae within Columbea. ASTRAL-X also placed Columbimorphae as sister to Otidimorphae, matching the relationship presented in Stiller et al. [30].

##### Waterbirds were placed deep within a diverse neoavian clade

ASTRAL-X placed the waterbird clades Aequornithes and Phaethontimorphae deep within a broader clade containing Strisores, Cursorimorphae, and Opisthocomiformes. This topology is consistent with both the original extended avian study [30] and the STELAR-X reanalysis [23]. In contrast, some earlier studies placed waterbirds as sister to landbirds [29, 40].

##### Rheiformes was placed with Tinamiformes within Palaeognathae

Within Palaeognathae, ASTRAL-X placed Rheiformes as sister to Tinamiformes. This placement agrees with the original extended avian study [30] and the STELAR-X reanalysis [23]. It, however, contrasts with the topology reported by Cloutier et al. [41], where Rheiformes was placed as sister to the clade containing Apterygiformes and Casuariiformes.

##### Musophagiformes was placed within Otidimorphae

Consistent with Stiller et al. [30], ASTRAL-X placed Musophagiformes with Cuculiformes and Otidiformes within Otidimorphae, with high support. This differs from the STELAR-X reanalysis, where Musophagiformes was placed as sister to Strisores [23].

## 4 Discussion

We presented ASTRAL-X, which dramatically extends the practical limits of statistically consistent species tree estimation. By fundamentally redesigning the computational foundations of the dominant quartet-based framework ASTRAL, ASTRAL-X reconstructs species trees containing 300,000 taxa in a few hours of computation using only 100 GB of memory, whereas previous ASTRAL implementations encounter prohibitive memory and runtime limitations well below this scale. Notably, ASTRAL-X achieves this without sacrificing ASTRAL’s statistical guarantees and empirical accuracy. This demonstrates that computational scalability and statistical rigor need not be competing objectives. Instead, careful algorithmic redesign can simultaneously preserve theoretical foundations that have made coalescent-based methods central to modern phylogenomics while dramatically expanding practical applicability.

Importantly, the improvements achieved by ASTRAL-X are not simply incremental. Nor is ASTRAL-X a straightforward adaptation of the computational techniques introduced in STELAR-X [23]. Although both STELAR-X (which was designed for rooted gene trees) and ASTRAL-X share the goal of overcoming scalability barriers through algorithmic redesign, the substantially greater complexity of optimization over unrooted quartets required fundamentally new data structures and algorithmic techniques. Previous generations of ASTRAL largely focused on improving computational efficiency within the existing algorithmic framework. In contrast, ASTRAL-X fundamentally redesigns the underlying computational representations and workflow through new data structures, scalable algorithms for candidate cluster generation and tripartition scoring, efficient precomputation strategies, heterogeneous CPU–GPU execution, and several complementary mathematical, graph-theoretic, and algorithmic innovations that together overcome the computational bottlenecks of the ASTRAL family of methods. Notably, ASTRAL-X achieves the asymptotically optimal memory complexity of *O*(*nk*) in the tree-local search space mode, essentially matching the input size and allowing analyses to remain feasible as long as the input trees fit in memory. Because reducing memory consumption often comes at the cost of increased running time as they generally involve competing design trade-offs, simultaneously attaining an asymptotically optimal memory footprint and dramatically reducing runtime represents a remarkable algorithmic advance.

The ability to reconstruct statistically consistent species trees from unrooted gene trees at this scale has important implications for emerging phylogenomic initiatives. Large collaborative efforts, such as the Angiosperm Tree of Life [19,42] and the Earth BioGenome Project [20,43], are rapidly generating genomic data for hundreds of thousands of species. Until now, applying statistically consistent coalescent-based methods directly to unrooted gene trees at this scale has been largely impractical. By making analyses at this scale feasible using readily available computational resources, ASTRAL-X enables comprehensive phylogenomic studies for ambitious community initiatives to resolve the tree of life across diverse lineages, and substantially broadens the scope of questions that can be addressed using genome-scale data.

Our results also suggest a broader lesson for computational phylogenomics. The long-standing scalability limitations of statistically consistent summary methods have often been regarded as an inherent consequence of the complexity of the underlying models and optimization problems. The success of ASTRAL-X indicates that these limitations are, to a considerable extent, algorithmic rather than statistical. Together, STELAR-X and ASTRAL-X reflect a broader opportunity to rethink the computational foundations of coalescent-based phylogenomics. We anticipate that many existing summary methods may benefit from similar redesigns and that these principles will help guide the next generation of scalable phylogenomic inference algorithms.

This study can be extended in several important directions. One natural direction is to further broaden the candidate search space while retaining the computational efficiency of ASTRAL-X, particularly for difficult datasets with substantial gene tree error or missing taxa. The species tree inferred by ASTRAL-X could also be used as a high-quality starting point for local tree-search procedures based on moves such as NNI or SPR, allowing the topology to be further refined beyond the original candidate search space. Such post-inference refinement may help recover improved solutions while preserving the scalability of the main ASTRAL-X inference framework. Another important direction is to extend ASTRAL-X to multi-copy gene family trees so that it can account for paralogous sequences and gene duplication and loss. These directions would further expand the applicability of ASTRAL-X to increasingly large and complex phylogenomic datasets.

## Supporting information

Supplementary Methods, Results, Figures, and Tables

## 5 Code Availability

The source code for ASTRAL-X, along with the scripts used to run the experiments, is publicly available on GitHub at https://github.com/aaniksahaa/ASTRAL-X.

