## Supplementary Methods, Results, Figures, and Tables for "ASTRAL-X: Scaling Coalescent-Based Species Tree Inference to 300,000 Taxa"

### Supplementary material for ASTRAL-X: Scaling Coalescent-Based Species Tree Inference to 300,000 Taxa

#### 1 Example: Tuple Representation of Tripartitions and Computation of $\mathrm{QI}(T, M)$

We use 0-based array indexing throughout this example, with both endpoints of each subarray included. Consider the taxon set

$$L = \{A, B, C, D, E\}$$

and two unrooted binary gene trees

$$g_1 = \big((A, B),\, C,\, (D, E)\big), \qquad g_2 = \big((A, B),\, E,\, (C, D)\big).$$

We use a tripartition $M$ induced by $g_1$ as a gene-tree tripartition and a tripartition $T$ induced by $g_2$ as a candidate tripartition. The example first illustrates their compact tuple representations and then computes the intersection counts between them using the four methods supported by ASTRAL-X: prefix sums, smaller-side traversal, postorder propagation, and bitwise operations.

**Tuple representation of the gene-tree tripartition.** We root $g_1$ at the edge separating the $\{A, B\}$ subtree from the remainder. The resulting rooted topology and postorder leaf array are

$$\big((A, B),\, (C, (D, E))\big), \qquad \mathcal{A}_1 = [A, B, C, D, E].$$

Consider the non-root internal node whose child subtrees contain $\{C\}$ and $\{D, E\}$. The taxa outside this node form the third side $\{A, B\}$, yielding

$$M = (M_1 \mid M_2 \mid M_3) = (\{C\} \mid \{D, E\} \mid \{A, B\}).$$

Its three sides are represented as

$$\begin{aligned} M_1 &= \mathcal{A}_{1,2,2}, && (1, 2, 2, 0),\\ M_2 &= \mathcal{A}_{1,3,4}, && (1, 3, 4, 0),\\ M_3 &= L \setminus \mathcal{A}_{1,2,4}, && (1, 2, 4, 1). \end{aligned}$$

The final component is the complement flag: 0 denotes the specified subarray, whereas 1 denotes its complement in the leaf set of the source gene tree.

**Tuple representation of the candidate tripartition.** We similarly root $g_2$ at the edge separating the $\{A, B\}$ subtree from the remainder. This gives

$$\big((A, B),\ (E, (C, D))\big), \qquad \mathcal{A}_2 = [A, B, E, C, D].$$

The non-root internal node with child subtrees $\{E\}$ and $\{C, D\}$ induces the tripartition

$$\{E\} \mid \{C, D\} \mid \{A, B\}.$$

Since the order of the three sides is immaterial, we write the candidate tripartition as

$$T = (P \mid Q \mid R) = (\{A, B\} \mid \{C, D\} \mid \{E\}).$$

Its sides are represented in $\mathcal{A}_2$ as

$$\begin{aligned} P &= L \setminus \mathcal{A}_{2,2,4}, && (2, 2, 4, 1),\\ Q &= \mathcal{A}_{2,3,4}, && (2, 3, 4, 0),\\ R &= \mathcal{A}_{2,2,2}, && (2, 2, 2, 0). \end{aligned}$$

Thus, every tuple above represents an actual cluster induced by its source gene tree.

**Core intersection counts.** To evaluate $\mathrm{QI}(T, M)$, we first require the four core intersections between the two explicitly stored sides of $T$ and those of $M$:

$$a_1 = |P \cap M_1|, \qquad a_2 = |P \cap M_2|, \qquad b_1 = |Q \cap M_1|, \qquad b_2 = |Q \cap M_2|.$$

We now show how each intersection method obtains these same four values.

**Computation using prefix sums.** We record membership in $P$ and $Q$ along the postorder array of $g_1$:

$$\mathrm{ind}_{P,1} = [1, 1, 0, 0, 0], \qquad \mathrm{ind}_{Q,1} = [0, 0, 1, 1, 0].$$

For $C \in \{P, Q\}$, define

$$\mathrm{pfx}_{C,1}[\ell] = \sum_{j=0}^{\ell-1} \mathrm{ind}_{C,1}[j].$$

The resulting prefix arrays are

| $\ell$ | 0 | 1 | 2 | 3 | 4 | 5 |
|---|---|---|---|---|---|---|
| $\mathrm{pfx}_{P,1}[\ell]$ | 0 | 1 | 2 | 2 | 2 | 2 |
| $\mathrm{pfx}_{Q,1}[\ell]$ | 0 | 0 | 0 | 1 | 2 | 2 |

Since $M_1$ occupies positions $[2, 2]$ and $M_2$ occupies positions $[3, 4]$, the four counts follow from prefix differences:

$$\begin{aligned} a_1 &= \mathrm{pfx}_{P,1}[3] - \mathrm{pfx}_{P,1}[2] = 2 - 2 = 0,\\ a_2 &= \mathrm{pfx}_{P,1}[5] - \mathrm{pfx}_{P,1}[3] = 2 - 2 = 0,\\ b_1 &= \mathrm{pfx}_{Q,1}[3] - \mathrm{pfx}_{Q,1}[2] = 1 - 0 = 1,\\ b_2 &= \mathrm{pfx}_{Q,1}[5] - \mathrm{pfx}_{Q,1}[3] = 2 - 1 = 1. \end{aligned}$$

The terminal prefix values also give the row totals

$$|P \cap L| = \mathrm{pfx}_{P,1}[5] = 2, \qquad |Q \cap L| = \mathrm{pfx}_{Q,1}[5] = 2.$$

**Computation using smaller-side traversal.** The smaller-side method traverses the smaller cluster in each pair and tests membership in the other cluster through the inverse postorder index of its source tree. The relevant inverse indices are

$$\pi_1(A) = 0, \quad \pi_1(B) = 1, \quad \pi_1(C) = 2, \quad \pi_1(D) = 3, \quad \pi_1(E) = 4,$$
$$\pi_2(A) = 0, \quad \pi_2(B) = 1, \quad \pi_2(E) = 2, \quad \pi_2(C) = 3, \quad \pi_2(D) = 4.$$

For $P \cap M_1$, we traverse the singleton $M_1 = \{C\}$. The tuple for $P$ represents the complement of positions $[2, 4]$ in $\mathcal{A}_2$. Since $\pi_2(C) = 3$ lies inside this range, $C \notin P$, and hence

$$|P \cap M_1| = 0.$$

For $P \cap M_2$, we traverse $P = \{A, B\}$. Neither $\pi_1(A) = 0$ nor $\pi_1(B) = 1$ lies in the range $[3, 4]$ representing $M_2$, giving

$$|P \cap M_2| = 0.$$

Similarly, $\pi_2(C) = 3$ lies in the range $[3, 4]$ representing $Q$, so

$$|Q \cap M_1| = 1.$$

Finally, for $Q \cap M_2$, we traverse $Q = \{C, D\}$. Here, $\pi_1(C) = 2$ lies outside $[3, 4]$, whereas $\pi_1(D) = 3$ lies inside it. Therefore,

$$|Q \cap M_2| = 1.$$

The smaller-side method thus gives

$$(a_1, a_2, b_1, b_2) = (0, 0, 1, 1).$$

**Computation using postorder propagation.** The postorder-propagation method obtains the same counts by combining states along $g_1$. For each node $u$, recall that

$$\mathbf{s}_T(u) = \big(|P \cap L_1(u)|,\ |Q \cap L_1(u)|,\ |L_1(u)|\big).$$

The relevant leaf states are

$$\mathbf{s}_T(C) = (0, 1, 1), \qquad \mathbf{s}_T(D) = (0, 1, 1), \qquad \mathbf{s}_T(E) = (0, 0, 1).$$

Let $u_3$ be the internal node with children $D$ and $E$. Its propagated state is

$$\mathbf{s}_T(u_3) = \mathbf{s}_T(D) + \mathbf{s}_T(E) = (0, 1, 2).$$

The node inducing $M$ has children $C$ and $u_3$. Its first child therefore contributes

$$|P \cap M_1| = 0, \qquad |Q \cap M_1| = 1,$$

while its second child contributes

$$|P \cap M_2| = 0, \qquad |Q \cap M_2| = 1.$$

Hence, postorder propagation again yields

$$(a_1, a_2, b_1, b_2) = (0, 0, 1, 1).$$

**Computation using bitwise operations.** For the bitset method, let the bit positions follow the global taxon order $(A, B, C, D, E)$. The relevant clusters are represented by

$$\mathbf{B}(P) = (1, 1, 0, 0, 0), \quad \mathbf{B}(Q) = (0, 0, 1, 1, 0),$$
$$\mathbf{B}(M_1) = (0, 0, 1, 0, 0), \quad \mathbf{B}(M_2) = (0, 0, 0, 1, 1).$$

Their bitwise intersections are

$$\mathbf{B}(P) \,\&\, \mathbf{B}(M_1) = (0, 0, 0, 0, 0), \quad \mathbf{B}(P) \,\&\, \mathbf{B}(M_2) = (0, 0, 0, 0, 0),$$
$$\mathbf{B}(Q) \,\&\, \mathbf{B}(M_1) = (0, 0, 1, 0, 0), \quad \mathbf{B}(Q) \,\&\, \mathbf{B}(M_2) = (0, 0, 0, 1, 0).$$

Counting the set bits gives

$$|P \cap M_1| = 0, \qquad |P \cap M_2| = 0, \qquad |Q \cap M_1| = 1, \qquad |Q \cap M_2| = 1.$$

Thus, all four methods recover the same core intersection counts:

$$(a_1, a_2, b_1, b_2) = (0, 0, 1, 1).$$

They differ only in how these values are computed.

**Recovering the remaining intersections.** Because $g_1$ is complete over $L$, the row totals are

$$|P \cap L| = |P| = 2, \qquad |Q \cap L| = |Q| = 2.$$

The third-column entries for $P$ and $Q$ are therefore

$$a_3 = |P \cap M_3| = 2 - a_1 - a_2 = 2,$$

$$b_3 = |Q \cap M_3| = 2 - b_1 - b_2 = 0.$$

The entries involving $R$ follow from the column totals:

$$\begin{aligned} c_1 &= |R \cap M_1| = |M_1| - a_1 - b_1 = 1 - 0 - 1 = 0, \\ c_2 &= |R \cap M_2| = |M_2| - a_2 - b_2 = 2 - 0 - 1 = 1, \\ c_3 &= |R \cap M_3| = |M_3| - a_3 - b_3 = 2 - 2 - 0 = 0. \end{aligned}$$

The complete intersection matrix is consequently

| | $M_1 = \{C\}$ | $M_2 = \{D, E\}$ | $M_3 = \{A, B\}$ |
|---|---|---|---|
| $P = \{A, B\}$ | 0 | 0 | 2 |
| $Q = \{C, D\}$ | 1 | 1 | 0 |
| $R = \{E\}$ | 0 | 1 | 0 |

whose row and column sums agree with the corresponding cluster sizes.

**Computing** $\mathrm{QI}(T, M)$**.** We have

$$(a_1, a_2, a_3) = (0, 0, 2), \qquad (b_1, b_2, b_3) = (1, 1, 0), \qquad (c_1, c_2, c_3) = (0, 1, 0).$$

Using Equation (4) in the main text,

$$\mathrm{QI}(T, M) = \sum_{(i,j,k) \in S_3} \frac{a_i + b_j + c_k - 3}{2} a_i b_j c_k.$$

Every term vanishes except the one corresponding to $(i, j, k) = (3, 1, 2)$. Hence,

$$\begin{aligned} \mathrm{QI}(T, M) &= \frac{a_3 + b_1 + c_2 - 3}{2} a_3 b_1 c_2 \\ &= \frac{2 + 1 + 1 - 3}{2} \cdot 2 \cdot 1 \cdot 1 \\ &= 1. \end{aligned}$$

This contribution corresponds to the quartet $\{A, B, C, E\}$. In both tripartitions, $A$ and $B$ occur together, while $C$ and $E$ occupy the two remaining sides. Both therefore induce the quartet topology

$$AB \mid CE.$$

According to Equation 5 in the main text, the contribution of $M$ to the weight of $T$ is

$$\frac{1}{2} f(M)\, \mathrm{QI}(T, M) = \frac{f(M)}{2},$$

where $f(M)$ is the frequency of $M$ among the input gene-tree tripartitions.

### 2 Running Time and Memory Complexity of Consensus-guided Cluster Expansion

**Theorem 2.1** (Complexity of consensus-guided cluster expansion)**.** *Suppose that ASTRAL-X uses a constant number of greedy-consensus frequency thresholds and a constant number of representative-sampling rounds for resolving each consensus polytomy. Then the consensus-guided expansion of the candidate cluster set $X$ introduces at most $O(n)$ additional clusters and requires $O(nk)$ additional memory.*

*The running time of this procedure is*

$$O\big(n^2 k + nk\log(nk) + n^2 \log n\big)$$

*in the worst case. When the input gene trees and the evolving consensus hierarchies have logarithmic height, the running time becomes*

$$O\big(nk\log^2 n + nk\log(nk) + n^2 \log n\big)\,.$$

.

*Proof.* Across $k$ gene trees on at most $n$ taxa, there are $O(nk)$ gene-tree clusters in total and therefore at most $O(nk)$ distinct clusters whose frequencies need to be considered. Their frequencies can be counted using the hash signatures maintained by ASTRAL-X in expected $O(nk)$ time. Ordering the distinct clusters in decreasing order of frequency requires $O(nk\log(nk))$ time using comparison sorting. Since ASTRAL-X uses only a constant number of consensus thresholds, this ordering and the subsequent scans of the candidate clusters contribute the same asymptotic cost across all consensus trees.

We first consider the cost of constructing the greedy consensus hierarchies. Starting from an unresolved tree, ASTRAL-X processes the candidate clusters in decreasing order of frequency and inserts a cluster only when it is compatible with the clusters accepted earlier. At every stage, the accepted clusters form a laminar hierarchy on at most $n$ taxa. For a proposed cluster, the algorithm follows the relevant ancestor paths in this hierarchy, identifies the smallest region containing the cluster, and determines whether the proposed cluster is a union of complete child subtrees.

In the worst case, the evolving hierarchy may have linear height, and a single compatibility test may therefore inspect $O(n)$ nodes. Since at most $O(nk)$ distinct gene-tree clusters are proposed, and the number of consensus thresholds is constant, the total cost of testing compatibility is bounded by

$$O(n)\cdot O(nk) = O(n^2 k).$$

A lower asymptotic bound follows when the trees are balanced. Consider first the clusters contributed by one balanced binary gene tree. At depth $\ell$, there are $O(2^\ell)$ clusters, each containing $O(n/2^\ell)$ taxa. Hence, the total cardinality of all clusters at that depth is $O(n)$. Since a balanced tree has $O(\log n)$ levels, the sum of the cardinalities of all of its clusters is $O(n\log n)$. Across $k$ gene trees this becomes

$$O(nk\log n).$$

If the evolving consensus hierarchy also has height $O(\log n)$, processing a taxon occurrence in a proposed cluster requires following at most $O(\log n)$ relevant ancestor levels. Consequently, the total ancestor-path traversal over all proposed clusters is bounded by

$$O(nk\log n)\cdot O(\log n) = O(nk\log^2 n).$$

Thus the compatibility component improves from $O(n^2 k)$ in the worst case to $O(nk\log^2 n)$ when the relevant tree hierarchies are balanced.

We next consider the additional clusters obtained by resolving polytomies in the greedy consensus trees. Let the degrees of the polytomies in one such consensus tree be $d_1, d_2, \ldots, d_r$. Since a phylogenetic tree on $n$ taxa contains $O(n)$ nodes and edges, the sum of the degrees of its internal nodes, and hence the sum of the degrees of its polytomies, is linear:

$$\sum_{i=1}^{r} d_i = O(n).$$

For a polytomy of degree $d_i$, the UPGMA procedure produces a binary resolution of its $d_i$ incident arms. Any binary tree on $d_i$ leaves contains only $O(d_i)$ nontrivial clusters, so this resolution can add at most $O(d_i)$ clusters to $X$. Similarly, one representative-sampling round selects one taxon from each of the $d_i$ arms and constructs a tree on only these $d_i$ representatives. Such a tree also contains only $O(d_i)$ clusters, and expanding each representative cluster back to the corresponding union of complete polytomy arms does not change their number. Therefore, with a constant number of representative-sampling rounds, the total number of clusters introduced while resolving this polytomy is $O(d_i)$.

Summing over all polytomies in one consensus tree gives

$$\sum_{i=1}^{r} O(d_i) = O\left(\sum_{i=1}^{r} d_i\right) = O(n).$$

Because the number of consensus thresholds is constant, the entire consensus-guided expansion introduces only $O(n)$ additional clusters. In particular, starting from the $O(nk)$ clusters obtained from the gene trees, this procedure does not change the asymptotic bound on the size of the candidate cluster set:

$$|X| = O(nk).$$

The running time required for the UPGMA resolutions can also be bounded over all polytomies. A UPGMA resolution on $d_i$ arms can be computed in $O(d_i^2 \log d_i)$ time. Therefore,

$$\sum_{i=1}^{r} O(d_i^2 \log d_i) \leq O\left(\log n \sum_{i=1}^{r} d_i^2\right).$$

Using

$$\sum_{i=1}^{r} d_i^2 \leq \left(\sum_{i=1}^{r} d_i\right)^2 = O(n^2),$$

the total UPGMA cost for one consensus tree is $O(n^2 \log n)$, and remains of the same order across a constant number of consensus thresholds.

The representative-based resolutions give analogous bounds. For a degree-$d_i$ polytomy, the restricted gene trees contain at most $d_i$ sampled taxa each. Applying the same evolving-hierarchy compatibility procedure to these restricted trees requires $O(kd_i^2)$ time in the worst case. Summing over all polytomies,

$$O\left(k \sum_{i=1}^{r} d_i^2\right) = O(n^2 k).$$

When the restricted trees and their evolving consensus hierarchies have logarithmic height, the corresponding bound becomes

$$O\left(k \sum_{i=1}^{r} d_i \log^2 d_i\right) \leq O\left(k \log^2 n \sum_{i=1}^{r} d_i\right) = O(nk \log^2 n).$$

A constant number of representative-sampling rounds does not change either asymptotic bound.

Combining frequency ordering, consensus-hierarchy construction, UPGMA resolution, and representative-based resolution therefore gives

$$O\big(n^2 k + nk \log(nk) + n^2 \log n\big)$$

time in the worst case, and

$$O\big(nk \log^2 n + nk \log(nk) + n^2 \log n\big)$$

when the relevant gene-tree and consensus hierarchies are balanced.

Finally, we consider the additional memory required by this stage. The frequency map contains at most $O(nk)$ distinct gene-tree clusters, each represented by a constant-size signature. An evolving consensus hierarchy contains only $O(n)$ nodes, and the temporary structures used for a representative-based resolution contain at most $O(nk)$ total gene-tree information at any one time. As shown above, only $O(n)$ new clusters are generated by the consensus-guided expansion itself. Since consensus thresholds and polytomies can be processed sequentially, these structures do not accumulate multiplicatively. Hence the additional memory required by the entire consensus-guided expansion is

$$O(nk).$$

□

### 3 Constant-time computation of quartet similarity

We now describe the constant-time computation of the pairwise quartet similarity used in the similarity-matrix construction. A direct computation would require considering all quartets containing a queried pair of taxa. Instead, we exploit the structure of the path connecting the two taxa in a gene tree. The relevant quartet counts can be summarized by a small set of node-level quantities that are precomputed once for each gene tree and then reused for all pairwise queries.

Let $g_i$ be a gene tree with leaf set $L_i$, and let $m_i = |L_i|$. For two distinct taxa $x, y \in L_i$, there are

$$\binom{m_i - 2}{2}$$

quartets containing both $x$ and $y$, corresponding to the choice of the remaining two taxa. Let $q_i(x, y)$ denote the number of these quartets in which $x$ and $y$ occur on the same side of the induced quartet split. The similarity computation ultimately reduces to evaluating this quantity efficiently.

To avoid enumerating these quartets, we first count how many leaves lie in different parts of the rooted gene tree. For every node $v$ of $g_i$, let $\lambda_i(v)$ denote the number of leaves descended from $v$. Thus, $\lambda_i(v) = 1$ for a leaf, while for an internal node,

$$\lambda_i(v) = \sum_{c \in \mathrm{child}(v)} \lambda_i(c).$$

These values are obtained by a single bottom-up traversal.

The next quantity captures the size of the component branching away from a root-to-leaf path. For a non-root node $v$ with parent $p(v)$, define

$$h_i(v) = \lambda_i(p(v)) - \lambda_i(v).$$

Since the rooted representation is binary, $h_i(v)$ is precisely the number of leaves in the sibling subtree of $v$. Such sibling subtrees will form the components attached to the path between a queried pair of taxa. We associate with the edge $p(v) \to v$ the value

$$\phi_i(v) = h_i(v)\big(m_i - 2 - h_i(v)\big).$$

As shown below, this is exactly the contribution of that component when counting quartets in which the queried taxa lie on opposite sides.

Rather than summing these contributions separately for every query, we accumulate them along root-to-node paths. Let $r_i$ denote the root of $g_i$ and define

$$F_i(r_i) = 0, \qquad F_i(v) = F_i(p(v)) + \phi_i(v).$$

Once the descendant counts $\lambda_i(v)$ have been computed bottom-up, all prefix values $F_i(v)$ can be obtained in a single top-down traversal. Thus, the information required for later pairwise queries is prepared once for the entire gene tree.

We now consider a particular pair $x, y \in L_i$. Let

$$w = \mathrm{lca}_i(x, y),$$

and let $c_x$ and $c_y$ be the two children of $w$ lying on the paths from $w$ to $x$ and $y$, respectively. We write

$$a = \lambda_i(c_x), \qquad b = \lambda_i(c_y), \qquad z = m_i - a - b.$$

Here, $a$ and $b$ give the sizes of the two descendant sides of $w$ containing $x$ and $y$, while $z$ counts the leaves outside the subtree rooted at $w$. In particular, $z = 0$ when $w$ is the root.

It is convenient to first count the complementary set of quartets. Let $Q_i(x, y)$ denote the number of quartets containing $x$ and $y$ in which the two taxa occur on opposite sides of the quartet split. Along the path from $c_x$ to $x$, the relevant component contributions are accumulated by $F_i(x) - F_i(c_x)$. The corresponding contribution on the other side is $F_i(y) - F_i(c_y)$. The only remaining component consists of the $z$ leaves outside the subtree of $w$, and its contribution is

$$z(m_i - 2 - z) = z(a + b - 2) = (a - 1)z + (b - 1)z.$$

Combining the three parts gives

$$2Q_i(x, y) = F_i(x) - F_i(c_x) + F_i(y) - F_i(c_y) + (a - 1)z + (b - 1)z.$$

Since every quartet containing $x$ and $y$ places them either on the same side or on opposite sides, we therefore obtain

$$q_i(x, y) = \binom{m_i - 2}{2} - \frac{1}{2}\left(F_i(x) - F_i(c_x) + F_i(y) - F_i(c_y) + (a - 1)z + (b - 1)z\right).$$

Once the LCA and the corresponding node-level values have been retrieved, this expression requires only a constant number of arithmetic operations.

We briefly justify the counting argument underlying this expression. Consider the unique unrooted path between $x$ and $y$. After excluding $x$ and $y$ themselves, the remaining leaves can be partitioned according to the component through which they attach to this path. Let the sizes of these components be

$$h_1, h_2, \ldots, h_t, \qquad \sum_{j=1}^{t} h_j = m_i - 2.$$

Two additional taxa $u$ and $v$ induce the quartet $xy \mid uv$ exactly when they are drawn from the same component. Hence,

$$q_i(x, y) = \sum_{j=1}^{t} \binom{h_j}{2}.$$

Conversely, $x$ and $y$ occur on opposite sides when the other two taxa are drawn from two different components. Therefore,

$$Q_i(x, y) = \sum_{j<\ell} h_j h_\ell = \frac{1}{2} \sum_{j=1}^{t} h_j\big(m_i - 2 - h_j\big).$$

This form explains the definition of $\phi_i(v)$. Each sibling subtree along the $x$–$y$ path contributes one term of the form $h_j(m_i - 2 - h_j)$, and the prefix values $F_i$ allow all such terms on each side of the LCA to be recovered by a single subtraction. The component outside the LCA subtree contributes the remaining term involving $z$. Thus the precomputed quantities give exactly the desired quartet count without explicit quartet enumeration.

When gene trees are incomplete, not every tree contains a queried pair of taxa. We therefore aggregate information only over gene trees containing both $x$ and $y$ and having at least four leaves. The final similarity is

$$\mathcal{S}(x, y) = \frac{\displaystyle\sum_{\substack{i:\, x,y \in L_i \\ m_i \geq 4}} q_i(x, y)}{\displaystyle\sum_{\substack{i:\, x,y \in L_i \\ m_i \geq 4}} \binom{m_i - 2}{2}}.$$

If the denominator is zero, we set $\mathcal{S}(x, y) = 0$ for $x \neq y$, while $\mathcal{S}(x, x) = 1$. This aggregation weights each gene tree according to the number of informative quartets it contributes, rather than giving equal weight to gene trees of different sizes.

Finally, we need to retrieve the LCA and the associated node-level quantities efficiently for each pair. For this purpose, every gene tree is preprocessed using an Euler tour together with a sparse range-minimum-query structure. The LCA $\mathrm{lca}_i(x, y)$ can then be obtained in $O(1)$ time from the first Euler-tour occurrences of $x$ and $y$. The descendant counts and prefix values associated with the relevant nodes are stored during preprocessing, so all remaining quantities in the expression for $q_i(x, y)$ are also available in constant time. Using a sparse RMQ table, the preprocessing requires $O(m_i \log m_i)$ time and space for gene tree $g_i$, after which each pairwise similarity query takes $O(1)$ time.

### 4 Simulation Parameters

We follow the simulation parameters used in the STELAR-X study. The simulation parameters for the ILS-L2 condition are presented in Table S1. To generate the four ILS levels (ILS-L1 to ILS-L4), we varied the upper bound of the uniform distribution for effective population size from 150000 to 300000.

**Table S1:** Simulation parameters used to generate gene trees and species trees using SimPhy.

| **Parameter Name** | **Parameter Value** |
|---|---|
| Speciation rate | 0.000001 |
| Extinction rate | 0 |
| Duplication rate | 0 |
| Horizontal Gene Transfer rate | 0 |
| Ingroup divergence to the ingroup ratio | 1.0 |
| Generations | LogN(1.470055e+01,2.500000e-01) |
| Haploid effective population size | Uniform(100000,200000) |
| Global substitution rate | LogN(-1.727461e+01,6.931472e-01) |
| Species Tree Height | LogN(16.2,1) |
| Seed | 42 |

An example simulation command is presented below.

```
./simphy_lnx64 \
  -sb f:0.000001 \
  -ld f:0 \
  -lb f:0 \
  -lt f:0 \
  -rs 5 \
  -rl f:<NUM_GENE_TREES> \
  -rg 1 \
  -o <OUTPUT_DIR> \
  -sp u:100000,<SP_MAX> \
  -su ln:-17.27461,0.6931472 \
  -sg f:1 \
  -sl f:<NUM_TAXA> \
  -st ln:16.2,1 \
  -om 1 \
  -v 2 \
  -od 1 \
  -op 1 \
  -oc 1 \
  -on 1 \
  -cs 42
```

Here, <NUM_TAXA> denotes the number of taxa, <NUM_GENE_TREES> denotes the number of gene trees, and <SP_MAX> specifies the upper bound of the uniform distribution for the effective population size. We used <SP_MAX> values of 150,000, 200,000, 250,000, and 300,000 for ILS-L1, ILS-L2, ILS-L3, and ILS-L4, respectively.

### 5 Additional Results

#### 5.1 Additional Results on Biological Datasets

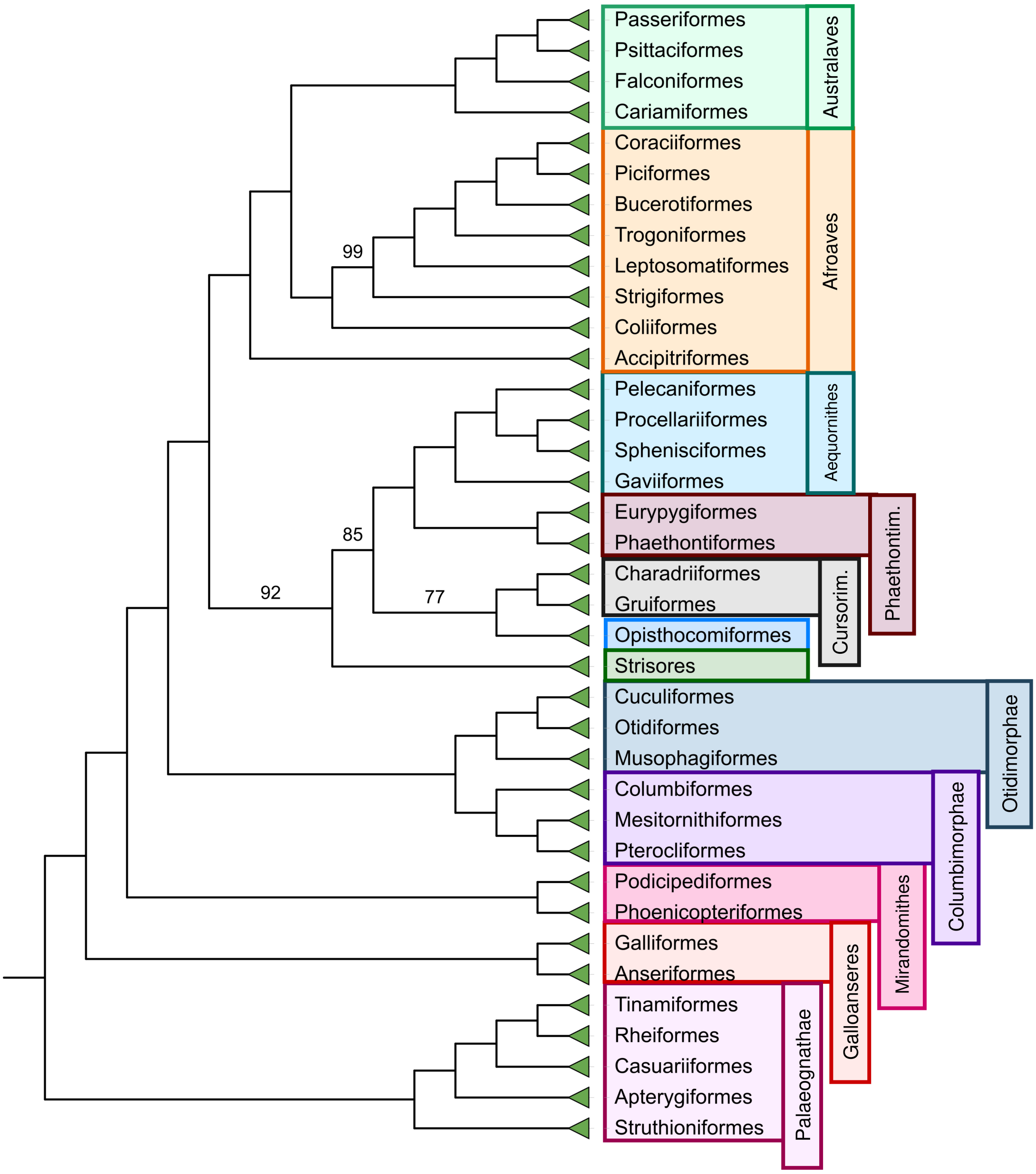


**Figure S1:** Species tree inferred by STELAR-X on the extended avian dataset with 363 taxa and 63,430 genes (intergenic loci). All branch support values are 100% except where noted.
